# Streamlined screening platforms lead to the discovery of pachysiphine synthase from *Tabernanthe iboga*

**DOI:** 10.1101/2024.06.30.601415

**Authors:** Mohamed O. Kamileen, Yoko Nakamura, Katrin Luck, Sarah Heinicke, Benke Hong, Maite Colinas, Benjamin R. Lichman, Sarah E. O’Connor

## Abstract

Plant-specialized metabolism is largely driven by the oxidative tailoring of key chemical scaffolds catalyzed by cytochrome P450 (CYP450s) enzymes. The monoterpene indole alkaloids tabersonine and pseudo-tabersonine, found in the medicinal plant *Tabernanthe iboga*, are extensively modified by oxidative reactions. Here we developed a streamlined screening strategy to screen the activity of *T. iboga* CYP450s in *Nicotiana benthamiana.* Using multigene constructs encoding the biosynthesis of tabersonine and pseudo-tabersonine scaffolds, we set out to uncover the CYP450s responsible for oxidative transformations of these scaffolds. Our approach identified two *T. iboga* cytochrome P450 enzymes: pachysiphine synthase (PS) and 16-hydroxy-tabersonine synthase (T16H). These enzymes catalyze an epoxidation and site-specific hydroxylation of tabersonine to produce pachysiphine and 16-OH-tabersonine, respectively. We further demonstrated that these genes produced the expected products when expressed in *Catharanthus roseus* flowers. This work provides new insights into the biosynthetic pathways of MIAs and underscores the utility of *N. benthamiana* and *C. roseus* as platforms for the functional characterization of plant enzymes.

## Introduction

Many natural products are derivatized by CYP450 enzymes, which enables extensive chemical diversity to be generated from a small set of starting scaffolds (Bathe & Tissier, 2019; Hansen *et al*., 2021; Nguyen & Dang, 2021). The medicinal plant *Tabernanthe iboga* is most widely known to produce the anti-addiction agent ibogaine, an MIA belonging to an iboga-type scaffold. However, in addition to ibogaine, this plant produces dozens of alkaloids derived from aspidosperma and pseudo-aspidosperma type scaffolds (Fig. 1). Many of these derivatives are generated through the action of CYP450s (Fig. 1).

**Fig. 1.**
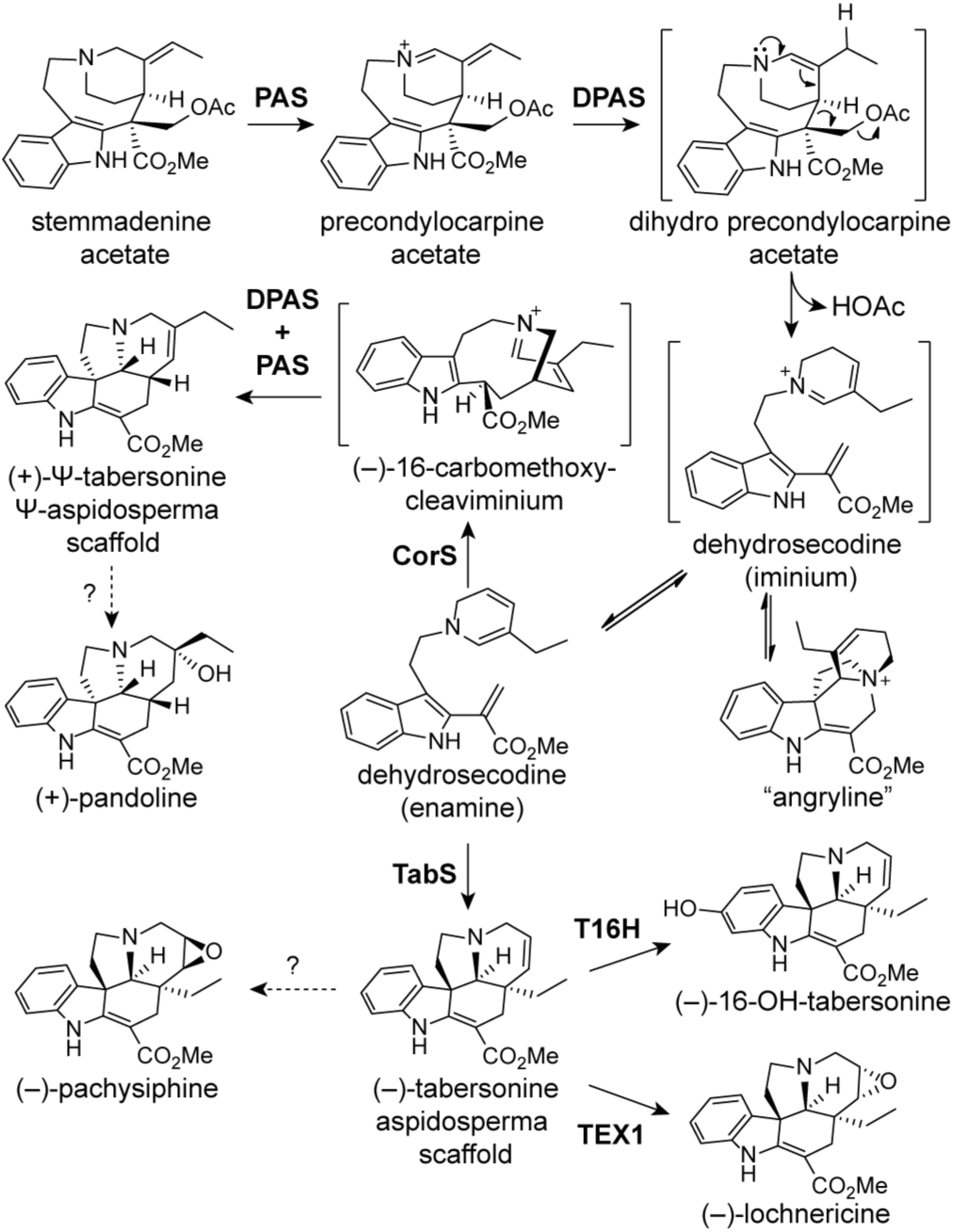
The enzymatic reactions involved in aspidosperma and pseudo-aspidosperma alkaloid biosynthesis. Stemmadenine acetate is a precursor to generate (–)-tabersonine (aspidosperma) and (+)-pseudo-tabersonine (pseudo-aspidosperma) alkaloids for further derivatization with known (solid arrows) and unknown (dashed arrows) oxidases (Farrow *et al*., 2019; Kamileen *et al*., 2022).

As part of a larger effort to elucidate the alkaloid pathways of *T. iboga*, we wanted to discover the CYP450s from this plant that catalyze the derivatization of these aspidosperma scaffolds. Discovery of the biosynthetic genes responsible for such derivatization steps requires screening of many candidate genes and the use of large quantities of valuable alkaloid starting substrates. Here we report a streamlined method for combinatorial screening of *T. iboga* CYP450s from the large CYP71 family. Using this method, we report here the discovery of two new CYP450 genes, pachysiphine synthase, and tabersonine 16-hydroxylase, from this plant. The resulting product is a predicted biosynthetic intermediate of several pharmacologically important dimeric alkaloids (Han-ya *et al*., 2011; Walia *et al*., 2020).

## Materials and Methods

### Chemicals

Stemmadenine acetate was synthesized from stemmadenine as previously described (Caputi *et al*., 2018). (±)-pseudo-tabersonine was synthesized, and (+)-pseudo-tabersonine was enzymatically generated as previously reported (Kamileen *et al*., 2022). (–)-tabersonine (cat no. T003900, CAS 4429-63-4) was purchased from Toronto Research Chemicals. Ajmaline (cat no. COMH93D5CE61, CAS 4360-12-7) was purchased from Sigma-Aldrich. (–)-[^13^C]-tabersonine was synthesized from (–)-tabersonine, as reported (Auriola *et al*., 1991).

### Plant growth conditions

*Tabernanthe iboga* plants were maintained as previously described (Kamileen *et al*., 2022). *Catharanthus roseus* var. Little Bright Eyes (LBE) was germinated in a standard soil mix in a climate chamber under 16 h light/8 h dark photoperiod at 23 °C, and 60% relative humidity. *C. roseus* var. LBE plants aged 6 – 8 months were used for flower petal transformation. *Nicotiana benthamiana* plants used for transient gene expression were grown in a greenhouse on a low-nutrient F1 compost soil mix under a 16 h light/8 h dark photoperiod at 22 °C, and 55% relative humidity. Plants were grown for 3-4 weeks before *Agrobacterium* infiltration.

### Cloning methods

Cytochrome P450 enzymes characterized in this study were amplified by PCR from a cDNA library generated from *T. iboga,* as previously described (Farrow *et al*., 2019). Known *C. roseus* cytochromes P450 genes (Table S1) were amplified from a cDNA library generated from *C. roseus* var. LBE. RNA. Primers with overhangs (Table S2) compatible with In-Fusion HD cloning kit (Takara Bioscience) were used to clone full-length genes into pESC-HIS (GenBank: AF063850.1, Agilent) vector for yeast expression and 3Ω1 (Sarrion-Perdigones *et al*., 2013) vector for plant transient expression (*N. benthamiana* or *C. roseus*) in the case of single transcriptional unit (STU) constructs. Multi-transcriptional unit constructs (MTU) for plant transient expression in *N. benthamiana* or *C. roseus* were assembled using the GoldenBraid 2.0 kit (Sarrion-Perdigones *et al*., 2013) (Addgene kit # 11000000076) (Fig. S1). For yeast expression, pESC-HIS gene constructs were transformed into yeast (*Saccharomyces cerevisiae*) strain WAT11 (*ade2*; contains the *Arabidopsis thaliana* cytochrome P450 reductase I gene, *ATR1*) (Urban *et al*., 1997) using the Frozen-EZ Yeast Transformation II Kit (Zymo Research). For plant transient expression, all STU and MTU constructs were transformed into *Agrobacterium tumefaciens* GV3101 cells (Goldbio) by electroporation as previously described (Kamileen *et al*., 2022). Detailed cloning methods are provided in Methods S1.

### Transient expression in *N. benthamiana*

The *Agrobacterium* strains harbouring gene constructs of interest were grown overnight at 28°C, 220 rpm. Cells were harvested by centrifugation at 4,000 × g for 10 min, and the supernatant was discarded. The pellet was gently resuspended in infiltration buffer (10 mM MES, pH 5.6, 10 mM magnesium chloride, 200 μM acetosyringone). Cells were re-harvested by centrifugation, and the supernatant was discarded. Cell pellets were resuspended in 10 ml of infiltration buffer and incubated in the dark at room temperature with gentle rocking for 2 hours. When multiple gene constructs were tested in combinations, strains were pooled in equal cell density. Each strain had an OD_600_ of 0.4 and infiltrated into the abaxial side of a 3-week-old *N. benthamiana* leaf using a 1 ml needleless syringe. When pooling for combinations of ≥ 10 constructs, the OD_600_ was set to 0.15. Each pooled combination consisted of a minimum of three biological replicates, where individual leaves of three different plants were infiltrated to randomize batch effects. Appropriate controls: wild-type, P19, and GFP were included in the experimental setup to be tested with and without substrate. Leaves were harvested four days post-*Agrobacterium* infiltration for plant disk assays.

### Transient expression in *C. roseus* flower petals

Transformation of *C. roseus* var. LBE flower petals were performed as previously described (Colinas *et al*., 2021; Colinas & Goossens, 2022). All open flowers were removed from the plant two days before infiltration. The *Agrobacterium* strains were prepared as outlined for transient expression in *N. benthamiana* to a final OD_600_ of 0.4. For each experiment, a minimum of three biological replicates from different individual plants were selected. Flower petals were pierced once with a sterile needle, and the *Agrobacterium* mixture was infiltrated using a 1 ml needleless syringe. Appropriate controls: wild-type, P19, and GFP were included in the experimental setup to be tested with and without substrate. Petals were harvested three days post-*Agrobacterium* infiltration for plant disk assays.

### Plant disk assays of *Agrobacterium* infiltrated *N. benthamiana* leaf and *C. roseus* flower petals

At harvest, 10 mm diameter leaf disks of *N. benthamiana* leaves, or *C. roseus* flower petals were cut using a borer. For each biological replicate, three leaf disks or petal disks were harvested. Each plant disk was then placed in an individual well of a 48-well plate with 300 μl of 50 mM HEPES buffer at pH 7.5. Where necessary, the substrate was added to a final concentration of 25 μM. Plates were closed and sealed with parafilm to avoid evaporation. Plates were incubated at 24 °C overnight (16-18 hours). After incubation, disks were collected, placed in a 2 ml safe-lock tube, and snap-frozen in liquid nitrogen. The disks were homogenized using a Tissue-Lyser II (Qiagen) at 25 Hz for 1 min with 3 mm tungsten beads. Metabolites were extracted from the powdered tissue with 300 μl of extraction solution (70% methanol in water + 0.1% formic acid) and sonicated for 10 min at room temperature. When needed, internal standard ajmaline (final concentration 20 μM) was spiked into the extraction solution before sample extraction. The samples were then centrifuged at 20,000 × g for 10 min to pellet cell debris, and the extract was filtered through 0.22 μm PTFE syringe filters into analytical vials for LC-MS analysis (Fig. S2).

### Heterologous expression in yeast and microsome preparation

Yeast cells (WAT11) were cultured in 100 ml of SD-His medium (6.7 g l^-1^ yeast nitrogen base without amino acids, 2 g l^-1^ drop-out mix without histidine, 74 mg l^-1^ adenine hemisulfate) containing 2% glucose (w/v) at 30 °C, 200 rpm for 28-34 hours. Cells were harvested and resuspended in 100 ml of SD-His medium + 1.8% galactose (w/v) + 0.2 % glucose (w/v) for protein production at 30 °C, 200 rpm for 18-24 hours. Yeast cells were harvested, and microsomes were prepared as reported (Pompon *et al*., 1996). Detailed yeast expression and microsome preparation methods are provided in Methods S1.

### Analytical-scale reactions with purified yeast microsomes

All analytical-scale *in vitro* yeast microsome reactions were carried out in 50 mM HEPES buffer (pH 7.5) in a volume of 100 μl. Each reaction consisted of 50 μg of total microsomal protein (0.50 μg μl^-1^ microsomal protein), 25 μM substrate, and 1 mM NADPH. Empty vector (EV) reactions were performed as a negative control. Reactions were initiated by adding microsomes and incubating at 30 °C and 600 rpm for 60 min, after which the reactions were quenched by adding 100 μl of extraction solution. Quenched reactions were centrifuged at 20,000 × g at room temperature for 10 min and filtered through 0.22-μm PTFE filters into analytical vials for LC-MS analysis.

### Semi-preparative-scale reactions with purified yeast microsomes

Reactions scaled to 10 ml volumes were set up to generate (–)-pachysiphine and (–)-16OH-tabersonine, from (–)-tabersonine and (–)-16OH-pachysiphine from (–)-pachysiphine. Reactions consisted of 50 mM HEPES pH 7.5 buffer, 2 mg substrate [(–)-tabersonine or (–)-pachysiphine] and, NADPH generation system [1.0 mM NADP^+^, 2.0 U ml^-1^ glucose-6-phosphate dehydrogenase (G6PDH), and 3.0 mM glucose-6-phosphate (G6P) along with 0.2 mM NADPH]. Reactions were initiated by the addition of 3 mg yeast microsomes (0.3 μg μl^-1^ microsomal protein), TiPS for (–)-pachysiphine or TiT16H for (–)-16OH-tabersonine and incubated at 30 °C, 200 rpm for 6 hours. To generate (–)-16OH-pachysiphine, a crude workup of 2 mg of (–)-pachysiphine reconstituted in 5% methanol (aqueous) was used as substrate with 3 mg (0.3 μg μl^-1^) of TiT16H microsomal protein preparation and incubated as described above. The reactions were terminated by the addition of 10 ml ethyl acetate. The aqueous reaction mixture was extracted thrice with ethyl acetate and evaporated to dryness. The dried extract was reconstituted in 5 ml of 10% methanol (aqueous) and subjected to SPE (solid phase extraction) as described in Methods S1.

### Semi-preparative HPLC isolation

Semi-preparative-scale product isolation was performed using high-performance liquid chromatography (HPLC). Reaction workups of (–)-pachysiphine, (–)-16OH-tabersonine, (–)-16OH-pachysiphine, and *N. benthamiana* extract of pseudo-tabersonine were subjected to HPLC isolation. An Agilent 1260 Infinity II HPLC instrument connected to an autosampler, diode array detector (DAD), and fraction collector for compound detection and isolation. Chromatography was monitored at UV 254 nm and 328 nm (Hisiger & Jolicoeur, 2007). Chromatographic separation was performed using a Phenomenex Kinetex XB-C18 (5.0 μm, 100 Å, 100 × 2.1 mm) column maintained at 40 °C under gradient elution using reversed phase conditions. Detailed HPLC methods are provided in Methods S1.

### Extraction of pseudo-tabersonine from *N. benthamiana*

To isolate pseudo-tabersonine, *N. benthamiana* (30 plants) transiently expressing the pS-psTab multigene construct were infiltrated with 400 μM stemmadenine acetate, 4 days post *Agrobacterium* infiltration. After 18 h, the substrate-infiltrated portions of the leaves were harvested for processing. Leaf material 20 g fresh weight was lyophilized using a freeze dryer. Dehydrated leaves were crushed and extracted with 50 ml of extraction solution at room temperature for 1 h with constant stirring. The extract was filtered through a miracloth and filter paper (MN 615, Macherey-Nagel) and evaporated to dryness. The dried extract was reconstituted in 10 ml of 10% methanol (aqueous) and subjected to SPE (solid phase extraction) as described in Methods S1.

### LC-MS analysis

LC-MS was performed using an UltiMate 3000 (Thermo Scientific) ultra-high performance liquid chromatography (UHPLC) system coupled to an Impact II UHR-Q-ToF (Ultra-High-Resolution Quadrupole-Time-of-Flight) mass spectrometer (Bruker Daltonics). LC-MS measurements were performed as previously reported (Kamileen *et al*., 2022). Data analysis was performed using Bruker Compass Data Analysis (Version 5.3) software. Detailed LC-MS methods are described in Methods S1.

### Chiral LC-MS analysis

Chiral separation was performed using the LC-MS instrumentation described above with changes to chromatographic conditions. A Phenomenex Lux Amylose-1 (150 x 3 mm, 3 µm particle) chiral LC column set at 30 °C was used to separate (±)-pseudo-tabersonine enantiomers. The column was used in reverse phase with mobile phase A (10 mM ammonium acetate) and B (acetonitrile). An isocratic gradient of 95% B and 0.6 ml min^-1^ flow rate was used.

### Structural elucidation by NMR analysis

NMR spectra were measured on a 700 MHz Bruker Advance III HD spectrometer (Bruker Biospin GmbH). Bruker TopSpin (version 3.6.1) was used for spectrometer control and data processing. CDCl_3_ and MeOH-*d_3_* were used as *d*-solvents and referenced to the residual solvent signals at δ_H_ 3.31 and δ_C_ 49.0 for CDCl_3_, and δ_H_ 3.31 and δ_C_ 49.0 for MeOH-*d_3_* respectively. For [^13^C]-tabersonine, NMR spectra were measured on a 500 MHz Bruker Advance III HD spectrometer (Bruker Biospin GmbH) using MeOH-*d_3_*.

### ECD spectra measurements of pachysiphine

The electronic circular dichroism (ECD) spectrum of (–)-pachysiphine was measured at 25°C on a JASCO J-810 spectropolarimeter (JASCO cooperation) using a 350 μl cell. Spectrometer control and data processing were accomplished using JASCO Spectra Manager II. The spectrum was measured in methanol at 0.35 mg ml^-1^ concentration. Detailed structure optimization and ECD spectral calculation for (–)-pachysiphine are described in Methods S1.

## Results

### Gene expression in *Nicotiana benthamiana* leaf disks

Transient expression of genes in *Nicotiana benthamiana* leaves is a well-established method for screening the biochemical activity of plant enzymes (Christ *et al*., 2019; Hong *et al*., 2022; Nett *et al*., 2023; Jiang *et al*., 2024; Martin *et al*., 2024). This process entails infiltrating the *N. benthamiana* leaf with a culture of *Agrobacterium tumefaciens* that harbors an expression plasmid containing the gene of interest. Within 3-5 days after infiltration, the protein is expressed in the leaf, and the resulting tissue can be analyzed by mass spectrometry to monitor what product(s) have formed as a result of the gene expression. If the expected substrate of the gene product is not naturally found in *N. benthamiana*, this substrate can be infiltrated into the leaf after protein expression using a needleless syringe (Fig. S2). However, relatively large amounts of substrate, usually 1 ml of a 0.1-1 mM solution, are required.

To minimize the amount of substrate needed for screening of protein activity, we modified the *N. benthamiana* protocol. The desired protein is transiently expressed in the *N. benthamiana* leaf using standard *Agrobacterium* transformation protocols. The leaf is allowed to remain attached to the plant for four days, allowing protein expression from the infiltrated gene. At this point, a cork borer is used to punch out disks of leaf tissue in the transfected leaf. These leaf disks are placed in a 24-well microplate, in which each well contains 300 µl of buffer with 25 µM of the substrate of interest. Leaf disks are incubated in this buffer solution for approximately 16 hours, allowing the substrate to diffuse into the disk. Then, the leaf disks are harvested, frozen in liquid nitrogen, ground, and extracted in methanol, and the methanolic extract is analyzed by LC-MS (Fig. S2). This method ensures substrates are effectively delivered into the leaf tissues using minimum substrate quantities, eliminating technical issues associated with substrate wastage and wash-off that commonly occur with traditional *N. benthamiana* syringe-based substrate infiltration. Additionally, the uniform fresh weight of the leaf disks and consistent substrate concentrations streamline the downstream sample preparation process for LC-MS analysis and data normalization.

### Gene stacked plasmids for expression of (–)-tabersonine and (+)-pseudo-tabersonine in *Nicotiana benthamiana*

Elucidation of multi-step natural product biosynthetic pathways often requires that more than one gene be expressed simultaneously in the *N. benthamiana* leaf. Typically, liquid cultures of individual *Agrobacterium* strains, each harboring a plasmid with a different gene of interest, are cultured separately, and then each culture is diluted to a defined optical density. The individual *Agrobacterium* cultures are then mixed together, and this mixture is then infiltrated into *N. benthamiana* leaves. However, stacking all pathway genes within a single plasmid has been shown in some cases to generate the final product of the enzyme sequence in higher amounts compared to co-infiltration with many *Agrobacterium* strains (Forestier *et al*., 2021; Kruse *et al*., 2024). Therefore, to screen for CYP450s that would act on either (–)-tabersonine and (+)-pseudo-tabersonine, we designed and built two multigene constructs to produce these alkaloid scaffolds from the starting substrate stemmadenine acetate using the Goldenbraid 2.0 kit (Sarrion-Perdigones *et al*., 2013). The three biosynthetic genes that are responsible for producing (–)-tabersonine and (+)-pseudo-tabersonine scaffolds from stemmadenine acetate have been previously reported (Fig. 1) (Farrow *et al*., 2019; Kamileen *et al*., 2022).

The first construct, containing the biosynthetic genes TiPAS3, TiDPAS1, and TiTabS, along with silencing suppressor P19, was designed to produce (–)-tabersonine (pS-Tab). The second construct, an analogous multigene construct containing the biosynthetic genes TiPAS3, TiDPAS1, and TiCorS, along with P19, was designed to produce (+)-pseudo-tabersonine (pS-psTab) (Fig. 2a, Fig. S1). Each transcriptional unit in these multigene constructs was regulated by the ubiquitin10 promoter (pUbq10) and terminator (tUbq10) from tomato (*Solanum lycopersicum*) (Table S1), elements known to be more stable and drive high gene expression (Grefen *et al*., 2010; Kumar *et al*., 2022).

**Fig. 2.**
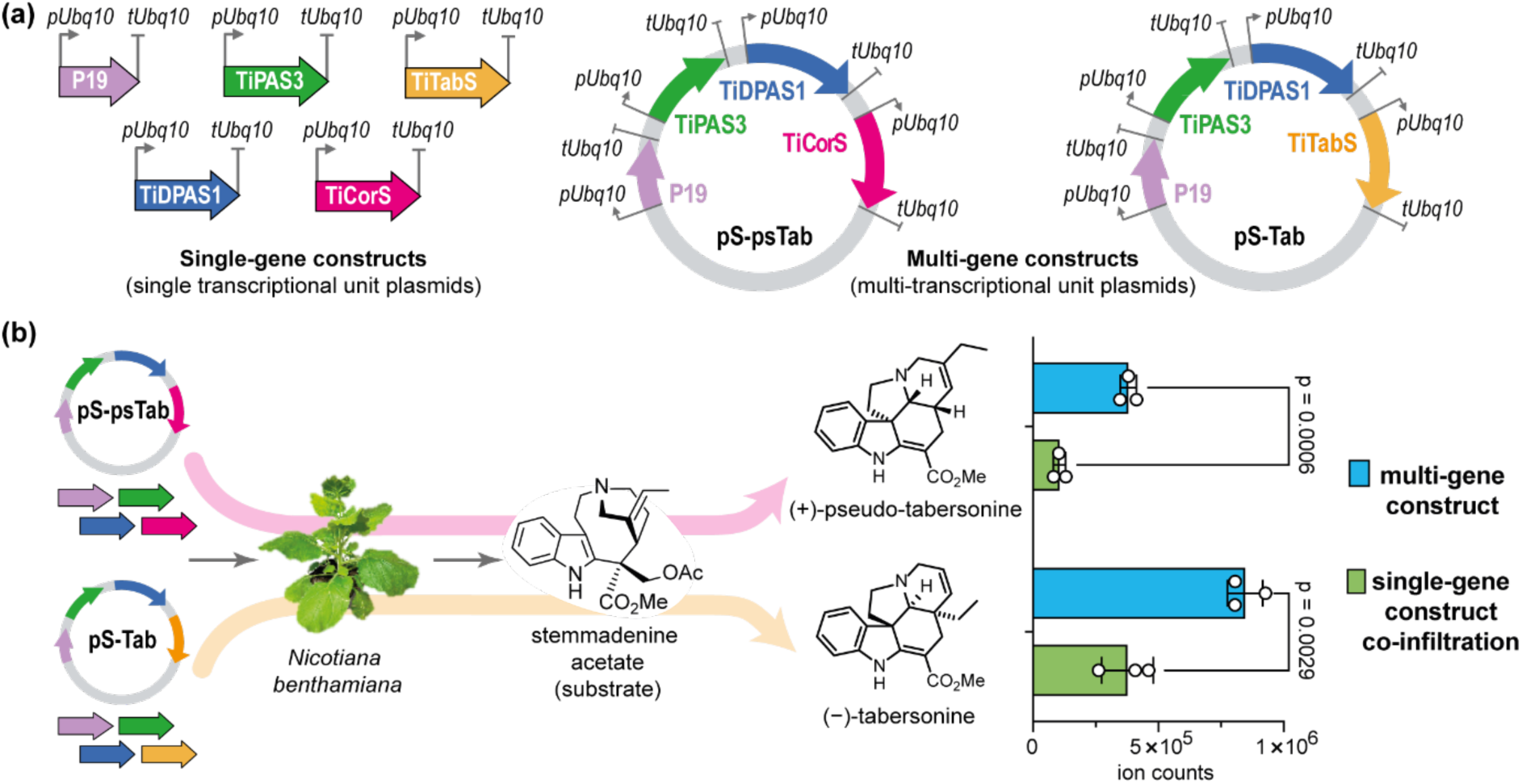
Multigene constructs for (–)-tabersonine and (+)-pseudo-tabersonine production via transient expression in *N. benthamiana*. (a) Multi-transcriptional unit constructs and single transcriptional unit constructs designed to generate (–)-tabersonine and (+)-pseudo-tabersonine in *N. benthamiana* using the exogenous substrate stemmadenine acetate. (b) Comparison of (–)-tabersonine and (+)-pseudo-tabersonine product observed in *N. benthamiana* when single transcriptional unit constructs (green bar) and the multi-transcriptional unit construct (blue bar) are used. The values represent mean ± SD (n = 3), two-tailed Student’s *t*-test.

We compared the levels of (–)-tabersonine and (+)-pseudo-tabersonine production that resulted from using these multigene constructs with levels that resulted from the conventional method in which we co-infiltrated four *Agrobacterium* strains harboring the corresponding single gene constructs (Fig 2a). For each experiment, leaf disks were incubated with 25 µM stemmadenine acetate, the starting substrate, in 300 µl buffer (pH 7.5). Notably, we recorded significantly higher levels of (–)-tabersonine and pseudo-tabersonine when using the multigene constructs (Fig. 2b Fig. S3). Since the stereochemistry of pseudo-tabersonine produced by this enzyme system had only been demonstrated *in vitro* (Kamileen *et al*., 2022), we isolated the pseudo-tabersonine produced by the pS-psTab multigene construct from *N. benthamiana* by analytical HPLC. We performed chiral HPLC with authentic standards of (+)-pseudo-tabersonine and a racemic (±)-pseudo-tabersonine. This confirmed that the product generated in *N. benthamiana* by the pS-psTab construct is indeed the expected (+)-pseudo-tabersonine isomer (Fig. S4).

### Screening of P450 genes using gene-stacked plasmids

These stacked plasmids were used to screen for CYP450 genes that encoded enzymes that oxidizes aspidosperma scaffolds. Each CYP450 gene candidate would be cloned on separate plasmids and then combined with the gene-stacked plasmid to create a flexible screening system. As a positive control to test this approach, we used the pS-TabS multigene construct to test the activity of previously characterized CYP450 genes known to derivatize (–)-tabersonine in *N. benthamiana* leaf disks (Fig. 3a). Using this system, we successfully demonstrated that the expected hydroxylated product (*m/z* 353) of (–)-tabersonine (*m/z* 337) catalyzed by the *C. roseus* CYP450 genes of the CYP71 family, T16H1, T16H2, T19H, TEX1, TEX2, and T3O could be detected (Fig. 3b). Furthermore, we observed that when these CYP71s were combined, the expected double-hydroxylated product (*m/z* 369) could also be detected (Fig. 3c).

**Fig. 3.**
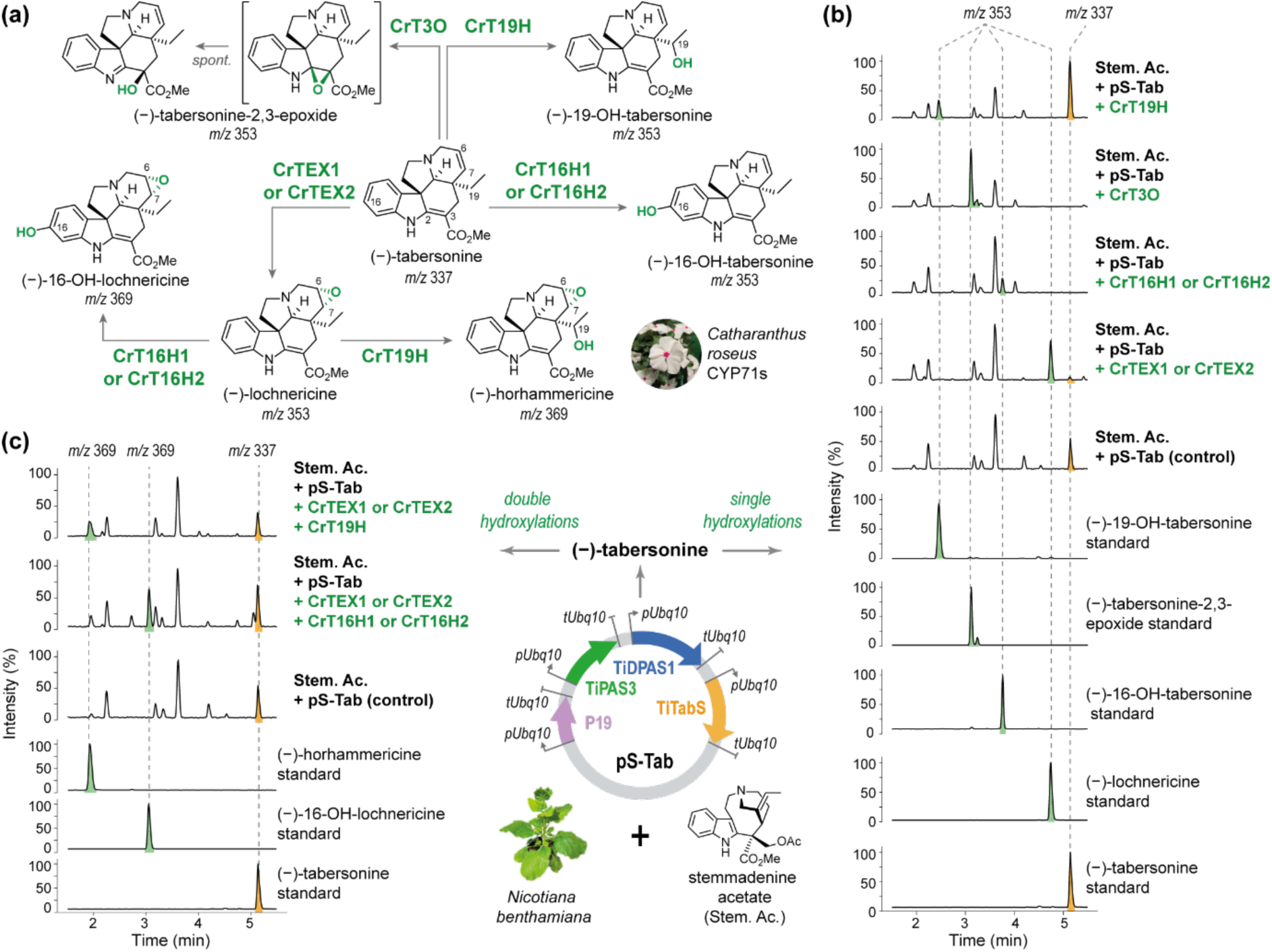
Hydroxylations of aspidosperma alkaloid (–)-tabersonine catalyzed by *Catharanthus roseus* CYP71 enzymes. (a) Functionally characterized *C. roseus* CYP71 P450 enzymes involved in (–)-tabersonine hydroxylation(s). (b) LC-MS profiles of the products resulting from a single hydroxylation of (–)-tabersonine in *N. benthamiana* leaf disk assays. (c) LC-MS profiles of the products resulting from double hydroxylations of tabersonine with combinations of selected *C. roseus* CYP71 genes in *N. benthamiana* leaf disk assays. Single and double hydroxylation products of (–)-tabersonine are confirmed by standards and MS^2^ spectra. LC-MS profiles are presented as total ion chromatograms, and *m/z* is presented as [M+H]^+^ of the parent ion.

### Discovery and functional characterization of new P450 genes

With this streamlined assay in hand, we inspected the *T. iboga* transcriptome to identify CYP450-encoding gene sequences (Farrow *et al*., 2019). Sequences assigned as a CYP450 were curated using transcriptome annotations and phylogenetic analysis (Fig S5). We chose to focus on the *T. iboga* CYP71 family, since many members of this enzyme family have been shown to derivatize monoterpene indole alkaloids in other plant species (Giddings *et al*., 2011; Besseau *et al*., 2013; Kellner *et al*., 2015; Tatsis *et al*., 2017; Carqueijeiro *et al*., 2018; Dang *et al*., 2018; Franke *et al*., 2019; Williams *et al*., 2019; Hong *et al*., 2022; Wang *et al*., 2022). A phylogenetic tree of *T. iboga* CYP450 sequences, which included CYP71 genes previously shown to be involved in MIA biosynthesis, was generated (Fig. S5). Using this tree, we curated a list of 30 CYP71 encoding genes from *T. iboga* in a phylogenetically informed manner (Fig. 4a).

**Fig. 4.**
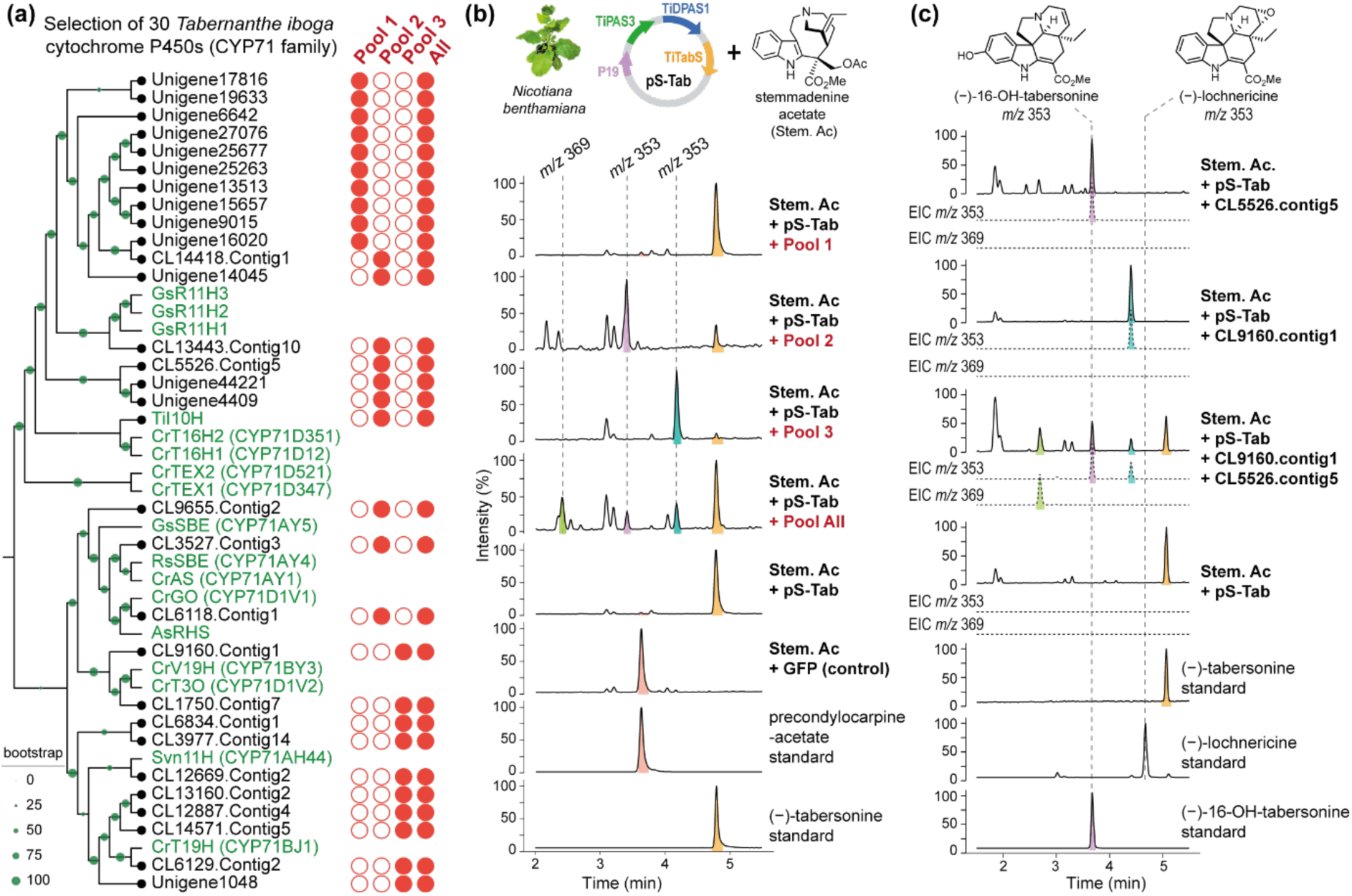
Screening of *T. iboga* CYP71 for enzymatic activity in *N. benthamiana*. (a) Phylogenetic tree of the 30 *T. iboga* CYP71 candidates (black circles) selected for pooled screening (red shaded circles). (b) LC-MS profiles of *T. iboga* CYP71 pools tested for hydroxylation activity in combination with the tabersonine-producing multigene construct pS-Tab and assayed in leaf disks with the substrate stemmadenine acetate Stemmadenine acetate is oxidized to precondylocarpine acetate by an endogenous *N. benthamiana* enzyme (Caputi *et al*., 2018). (c) Deconvolution of CYP71 pools 2 and 3 to identify gene candidate(s) responsible for the hydroxylation(s) of tabersonine observed in (b). LC-MS profiles in solid lines are presented as total ion chromatograms, and dotted lines as extracted ion chromatograms (EIC) with *m/z* presented as [M+H]^+^ of the parent ion.

Each CYP71 gene was cloned and transformed into an individual strain of *Agrobacterium*. For the initial screen, we tested the 30 *T. iboga* CYP71 candidates in pools of 10. Each set of *Agrobacterium* strains harboring the 10 genes of the pool were combined and then co-infiltrated into *N. benthamiana* leaves along with an *Agrobacterium* strain harboring either the pS-TabS or pS-psTab multigene construct. Infiltration was followed by leaf disk assays with the substrate stemmadenine acetate. When the *T. iboga* CYP71 pools were tested with the pS-psTab multigene construct, none of the candidates appeared to act on (+)-pseudo-tabersonine (Fig. S6). However, when the pS-TabS multigene construct was used, two of the three CYP71 candidate pools resulted in the production of a new, more hydrophilic product with a *m/z* 353 indicative of hydroxylation (Δ *m/z* 16) of (–)-tabersonine (*m/z* 337) (Fig. 4b). Additionally, when all three CYP71 candidate pools were combined, a third new product with a *m/z* 369 indicative of double hydroxylation (Δ *m/z* 32) of (–)-tabersonine (*m/z* 337) was observed along with the two *m/z* 353 products found in pool 2 and pool 3 (Fig. 4b). To identify the specific *T. iboga* CYP71 candidate(s) responsible for the hydroxylation(s) of (–)-tabersonine, we tested the individual candidates from *T. iboga* CYP71 pool 2 and 3 by co-infiltrating them with the pS-TabS multigene construct in the presence of stemmadenine acetate. We identified the two *T. iboga* CYP71 candidates responsible for the hydroxylation of (–)-tabersonine as CL5526.contig5 and CL9160.contig1 (Fig. 4c). Candidate CL5526.contig5 produced a hydroxylated product (*m/z* 353) that co-eluted and matched with identical MS^2^ spectra to the known compound 16OH-tabersonine and was therefore named *T. iboga* tabersonine-16-hydroxylase (TiT16H, assigned as CYP71BE297) (Fig. 4c). Additionally, we confirmed the structure of 16-OH-tabersonine by NMR, isolated from an *in vitro* enzymatic reaction containing (–)-tabersonine and yeast microsomes containing TiT16H protein (Fig. 5a, Fig. S7a, S8).

**Fig. 5.**
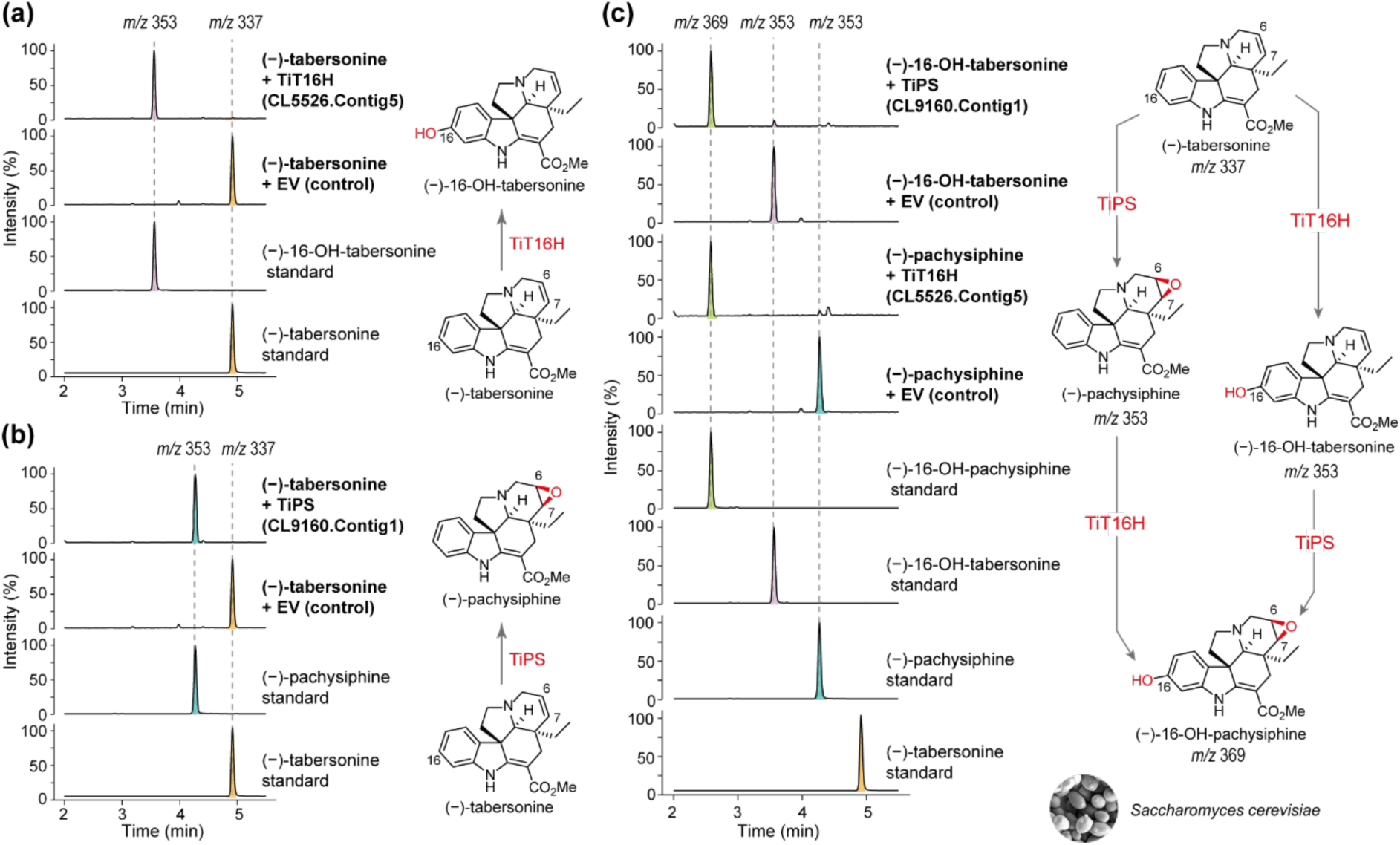
Characterization of *T. iboga* CYP450s in yeast. (a) Functional characterization of *T. iboga* tabersonine-16-hydroxylase (TiT16H) in yeast for the structure elucidation of 16-hydroxy-tabersonine (16-OH-tabersonine). (b) Functional characterization of *T. iboga* pachysiphine synthase (TiPS) in yeast for the structure elucidation of pachysiphine. (c) Elucidation of 16-hydroxy-pachysiphine (16-OH-pachysiphine) biosynthesis catalyzed by TiT16H and TiPS in yeast for structure validation. LC-MS profiles are presented as total ion chromatograms with *m/z* presented as [M+H]^+^ of the parent ion.

Interestingly, the MS^2^ spectrum of the hydroxylated product (*m/z* 353) of (–)-tabersonine observed from candidate CL9160.contig1 showed a fragmentation pattern similar to that of the epoxide (–)-lochnericine, which is known to be produced by the previously reported CYP71 TEX1 (Carqueijeiro *et al*., 2018) (Fig. 1, Fig. S9). To determine the product produced by candidate CL9160.contig1, we expressed this gene in yeast, purified yeast microsomes, and incubated the microsomes with (–)-tabersonine to isolate the product. NMR analysis revealed that the product was a tabersonine-6,7-ꞵ-epoxide known as pachysiphine, which is structurally identical to lochnericine except for the stereochemistry of the epoxide (Fig. 5b, Fig. S7b, S10). Consequently, CL9160.contig1 was named *T. iboga* pachysiphine synthase (TiPS, assigned as CYP71D821). We performed CD spectroscopy of pachysiphine and assigned the stereochemistry as (–)-pachysiphine (Fig. S11). Surprisingly, (–)-pachysiphine has never been reported in *T. iboga*.

### Double oxidation of tabersonine by TiT16H and TiPS

Having discovered and functionally characterized TiPS and TiT16H, we combined these two genes with the pS-TabS multigene construct in *N. benthamiana* (Fig. 4c). We observed the formation of a double hydroxylated product corresponding to 16-OH-pachysiphine (*m/z* 369), a compound that has not been reported to be present in *T. iboga*. As described above for (–)-pachysiphine, we expressed both genes in yeast, prepared microsomes that contained TiPS and TiT16H, and then performed *in vitro* reactions with the substrates (–)-pachysiphine and (–)-16-OH-tabersonine. These reactions resulted in the formation of what appeared to be a hydroxylated pachysiphine product, and the yield was unaffected by the order in which the enzymes were added (Fig. 5c). To obtain the compound in sufficient quantities for NMR analysis, we scaled up the reaction containing (–)-pachysiphine with TiT16H protein and isolated the resulting product by HPLC. NMR analysis confirmed that the product was the expected 16OH-pachysiphine (Fig. S7c, S12). We carefully examined the metabolite profile of *T. iboga* tissue for the presence of pachysiphine and its hydroxylated derivatives. Although neither pachysiphine nor 16-OH-pachysiphine has been reported to be present in *T. iboga,* we observed the presence of both compounds in methanolic extracts of the young leaf of this plant (Fig. S13).

### *Catharanthus roseus* flowers as a screening platform

The *N. benthamiana* leaf disk assay minimizes the quantities of exogenous substrate required for testing biosynthetic enzymes and this led to the streamlined identification of TiPS and TiT16H. However, it can be challenging to obtain even small amounts of some chemically complex substrates involved in MIA biosynthesis (*e.g.* stemmadenine acetate) (Caputi *et al*., 2018). We reasoned that since many MIA-producing plants produce this upstream biosynthetic pathway intermediate if any of these plants could be transformed with these multi-gene constructs, the substrate could be accessed in situ. *T. iboga* is slow to germinate, and efforts to transform this plant have not been successful. However, the MIA-producing plant *Catharanthus roseus* is fast-growing and readily available, and although *C. roseus* leaves are recalcitrant to transient expression, the flowers of *C. roseus* have been successfully transfected to express transcription factors (Schweizer *et al*., 2018; Colinas *et al*., 2021). We surveyed the alkaloid profile of the flowers of the *C. roseus* cultivar “Little Bright Eyes” (LBE) using untargeted LC-MS and observed the presence of MIA precursors stemmadenine acetate and precondylocarpine acetate upon *Agrobacterium* infiltration (Fig. S14), suggesting that the substrate utilized by the genes encoded in the pS-Tab and pS-psTab multi-gene constructs is present in this tissue.

To test whether we could access the upstream stemmadenine acetate pools within flower tissue and heterologously produce active enzymes in flower petals, we transfected the petals with multigene constructs designed for tabersonine (pS-Tab) and pseudo-tabersonine (pS-psTab) without the addition of any exogenous substrate. After three days of infection, increased levels of tabersonine were observed in petals transfected with the tabersonine-producing multigene construct (pS-Tab) (Fig 6a). Tabersonine is present as a natural product in *C. roseus* LBE flowers in low abundance, as confirmed by comparison with a standard. However, when the flower was transfected with pS-TabS, the levels of tabersonine were substantially higher. Notably, *C. roseus* does not natively produce and accumulate pseudo-tabersonine (Fig. S14). However, in the case of the pseudo-tabersonine multigene construct (pS-psTab), we observed the formation of this compound in the flower petals (Fig 6a). The observation of both tabersonine over-production and pseudo-tabersonine accumulation demonstrates that the upstream stemmadenine acetate pools can be utilized within the flower and that multiple active enzymes can be heterologously expressed to catalyze the formation of native and non-native metabolites in the flower petals to explore downstream biosynthesis.

**Fig. 6.**
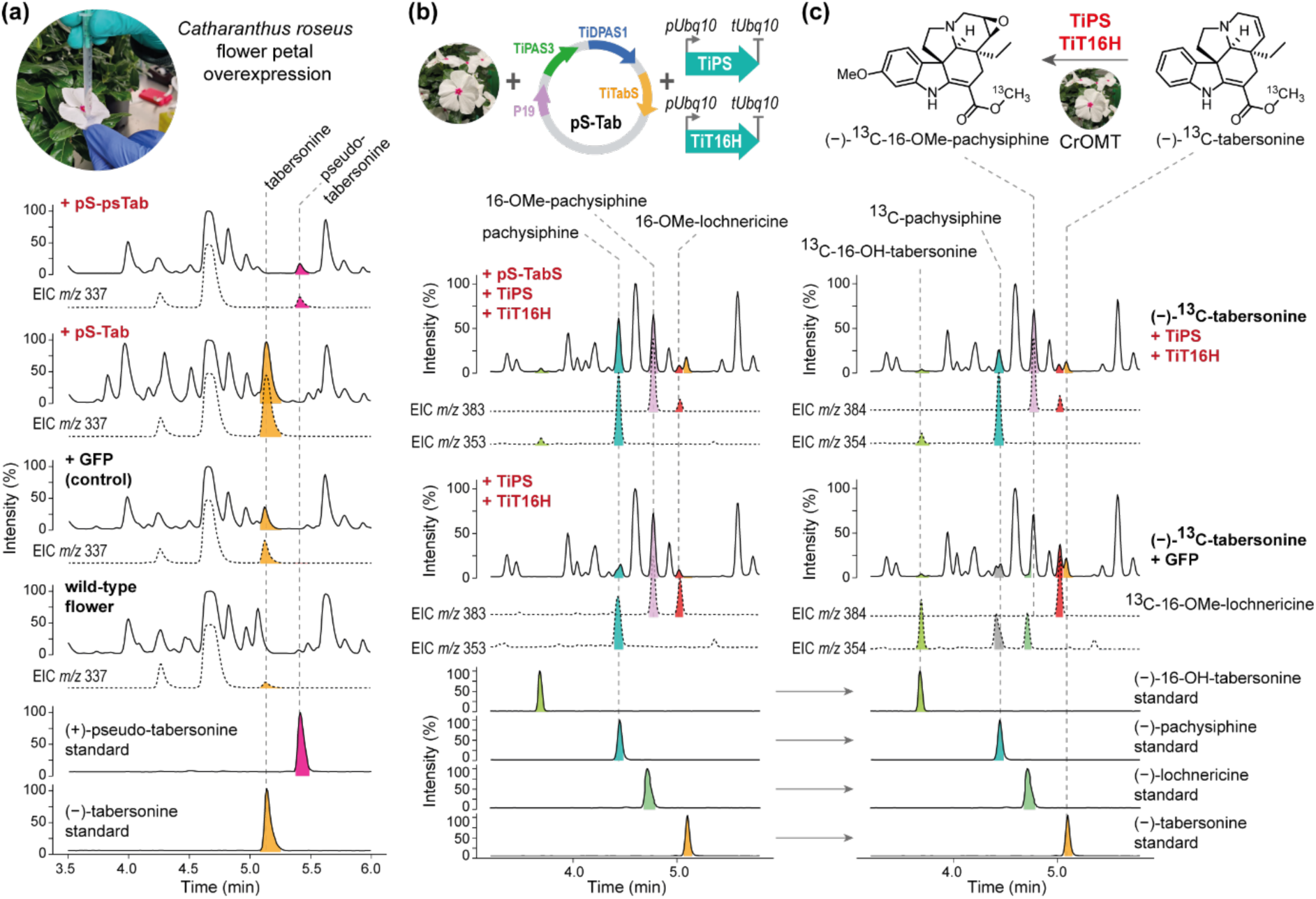
Transient overexpression of *T. iboga* enzymes in *C. roseus* var. LBE flowers. (a) LC-MS profiles of the products resulting from the transient expression of multigene constructs pS-Tab (tabersonine-producing) and pS-psTab (pseudo-tabersonine-producing) in *C. roseus* var. LBE flower petals. Expression of pS-Tab and pS-psTab results in the formation of tabersonine and pseudo-tabersonine, respectively in *C. roseus* flowers without the addition of exogenous substrate. (b) LC-MS profiles of the products generated by the co-expression of TiPS and TiT16H in combination with and separately from pS-Tab in *C. roseus* flowers. The expected product pachysiphine is increased when pS-Tab is co-expressed with TiPS and TiT16H alongside putative 16-OMe-pachysiphine catalyzed by endogenous enzyme(s). (c) LC-MS profiles of labeled pachysiphine and 16-OH-tabersonine generated by transient co-expression of TiPS and TiT16H in *C. roseus* flowers assayed with ^13^C-labeled tabersonine substrate. Consequent labeling of the putative 16-OMe-pachysiphine is also observed due to the endogenous catalytic activity of *C. roseus* enzymes in the flowers. LC-MS profiles in solid lines are presented as total ion chromatograms, dotted lines as extracted ion chromatograms (EIC) with *m/z* presented as [M+H]^+^ of the parent ion.

When the pS-TabS multigene construct was transfected into the flower petals of *C. roseus*, in addition to the product tabersonine, we unexpectedly observed the formation of its derivatized products 16-OH-tabersonine, lochnericine, and 16-OMe-tabersonine (Fig. S15). These modifications were likely catalyzed by endogenous *C. roseus* enzymes with specificity for tabersonine. To investigate the endogenous enzymatic activity catalyzed by *C. roseus* petals, we incubated wild-type flower petal disks in buffer (300 µl) with 25 µM ^13^C labeled (Fig. S16) and non-labeled tabersonine, the product of the multigene construct (pS-Tab). The tabersonine substrate was converted into the downstream alkaloids 16-hydroxy-tabersonine (16-OH-tabersonine), lochnericine, 16-methoxy-tabersonine (16-OMe-tabersonine), and 16-O-methoxy-tabersonine (16-OMe-lochnericine) as observed by the formation of equivalent ^13^C labeled products (Fig. S17). This confirmed the presence of active enzymes in this tissue able to derivatize tabersonine. In contrast, when the pS-psTab multigene construct was expressed, no downstream derivatization products of pseudo-tabersonine were detected. This lack of further modification is not surprising, as *C. roseus* does not naturally produce pseudo-tabersonine, and therefore, this plant would be expected to lack endogenous enzymes with specificity for this substrate.

Having established the native activity of *C. roseus* flower petals on the precursor tabersonine, we transfected flowers simultaneously with the multigene construct pS-Tab and two strains of *Agrobacterium* containing TiPS and TiT16H. After three days, the methanolic extracts of the transformed flower petals were analyzed by LC-MS for metabolic changes. We observed the heterologous production of pachysiphine and a compound with a mass corresponding to 16-O-methoxy pachysiphine (16-OMe-pachysiphine), both natural products not found natively in the *C. roseus* flower (Fig. 6b, Fig. S18). To corroborate this result, we incubated flower disks transfected simultaneously with TiPS and TiT16H in a buffer (300 µl) containing 50 µM ^13^C-labeled tabersonine. As expected, the transformed disks showed the formation of ^13^C-labeled pachysiphine and ^13^C-labeled 16-OMe-pachysiphine (Fig. 6c Fig. S19).

## Discussion

The monoterpene indole alkaloids (–)-tabersonine (aspidosperma) and (+)-pseudo-tabersonine (pseudo-aspidosperma) are alkaloid scaffolds found in many plant species that are modified through various oxidation reactions to generate chemical diversity (Giddings *et al*., 2011; Besseau *et al*., 2013; Kellner *et al*., 2015; Carqueijeiro *et al*., 2018). Here, we report the discovery of two tabersonine derivatizing enzymes from *T. iboga*, TiPS, and TiT16H. We used a miniaturized assay that uses *N. benthamiana* leaf disks to screen for enzyme activity, allowing the use of only small quantities of precious starting substrate. Furthermore, we also demonstrate that flower petals of the MIA-producing plant *C. roseus* can be transformed with the desired genes and that after gene expression, the resulting enzymes can access the biosynthetic intermediates present in this tissue as a source of in situ substrates. For monoterpene indole alkaloid biosynthetic enzymes that utilize rare or difficult-to-access substrates, this is a convenient and efficient expression platform for biosynthetic gene candidate screening.

The activity of TiT16H, which catalyzes the hydroxylation of the 16 position of tabersonine, has been previously identified in the closely related plant *C. roseus* (CrT16H). Surprisingly, while both of these enzymes belong to the CYP71 family, these genes are found in different clades in the phylogenetic tree (Figure 4a). TiT16H and CrT16H share only 56% amino acid identity (Fig. S20, S21), suggesting that the enzymatic activity in these two closely related plants may have evolved independently. We also discovered the enzyme that catalyzes the formation of the beta epoxide moiety on tabersonine to form pachysiphine (TiPS). Although pachysiphine is not found in *C. roseus*, this compound is structurally similar to lochnericine, a natural product that is found in *C. roseus* roots (Carqueijeiro *et al*., 2018). The only difference between lochnericine and pachysiphine is the stereochemistry of the epoxide. In *C. roseus*, the formation of this alpha epoxide moiety on the (–)-tabersonine substrate is catalyzed by CrTEX1 or CrTEX2 in the presence of tabersonine. However, TiPS, which catalyzes the formation of the beta epoxide on (–)-tabersonine to yield pachysiphine, has only 46% amino acid identity to CrTEX1 (45% identity with CrTEX2). Therefore, it would appear that the stereoselectivity of this epoxidation reaction also may have evolved independently. Finally, we show that these two enzymes can act together to generate 16-OH-pachysiphine from tabersonine. Although pachysiphine has not been reported to be found in *T. iboga*, after the discovery of these enzyme activities, we carefully inspected the metabolite profile of *T. iboga* tissues and saw the presence of both pachysiphine and 16-OH-pachysiphine in young leaves (Fig. S13). We also examined the gene expression profile of TiPS and TiT16H from the transcriptomic data and observed TiPS to be expressed in the roots, flowers, and young leaves, whereas TiT16H was expressed in the leaves of the plant (Fig. S22). It appears that pachysiphine, unlike 16-OH-tabersonine, may be transported to young leaves for storage.

Pachysiphine and 16-OH-pachysiphine have no known biological function but are well known to serve as precursors for a variety of dimeric alkaloids, including melodine K (Walia *et al*., 2020), connophylline (Han-ya *et al*., 2011), Conolodinines A–D (Sim *et al*., 2019), and scandomelidine (Mehri & Plat, 1992) which display a variety of pharmacological activities. Therefore, the discovery of these biosynthetic genes from *T. iboga* provides the foundation for assay and discovery of the biosynthetic genes that make this structurally complex family of dimeric alkaloids. Overall, *T. iboga* is a rich source of oxidative derivatization enzymes, and the miniaturized screening platforms described here can streamline the identification of these enzymes.

## Acknowledgements

We thank Prashant D. Sonawanne for advice on Goldenbraid cloning and design, Maritta Kunert for assistance with mass spectrometry, Eva Rothe, Franz Kaltofen, and the members of the Max Planck Institute for Chemical Ecology Research Green House for providing and tending to the *Catharanthus roseus*, *Tabernanthe iboga*, and *Nicotiana benthamiana* plants. We thank David Nelson for his assistance in assigning the CYP family of TiPS and TiT16H. We gratefully acknowledge the Max Planck Society for funding.

## Competing interests

None declared.

## Author contributions

SEO and MOK conceived the study and wrote the manuscript. MOK designed and performed experiments. SEO, BRL, and MOK critically analyzed the data. YN performed ECD, NMR analysis, and compound characterization. KL performed RNA isolation. SH developed the chiral LC-MS method. BH synthesized (–)-[^13^C]-tabersonine. MC assisted with *C. roseus* flower petal transformation.

## Data availability

The data supporting this study’s findings are available within the article and in the Supporting Information file. Sequences for TiPS (XXXX) and TiT16H (XXXX) have been deposited in GenBank.

## Supporting Information)

**Fig. S1** Design of multi-transcriptional unit constructs

**Fig. S2** *N. benthamiana* leaf disk assay workflow

**Fig. S3** Testing multi-gene constructs in *N. benthamiana*.

**Fig. S4** Chiral LC-MS analysis of pseudo-tabersonine.

**Fig. S5** Phylogenetic tree of *Tabernanthe iboga* (Ti) cytochrome P450s.

**Fig. S6** Iboga CYP71s assayed with (+)-pseudo-tabersonine in *N. benthamiana*.

**Fig. S7** HPLC isolation of hydroxylated tabersonine products from yeast workup.

**Fig. S8** Structure elucidation of (–)-16OH-tabersonine by NMR.

**Fig. S9** MS^2^ spectral comparison of lochnericine and CL9160.contig1 product.

**Fig. S10** Structure elucidation of (–)-pachysiphine by NMR.

**Fig. S11** Electronic circular dichroism (ECD) spectra of (–)-pachysiphine.

**Fig. S12** Structure elucidation of (–)-16OH-pachysiphine by NMR.

**Fig. S13** LC-MS metabolite profiles of *T. iboga* tissues.

**Fig. S14** MIA precursors present in *C. roseus* var. LBE flowers.

**Fig. S15** Derivatization of tabersonine by endogenous enzymes present in *C. roseus* flowers.

**Fig. S16** Structure confirmation of (–)-[^13^C]-tabersonine.

**Fig. S17** Incubation of *C. roseus* flowers with ^13^C-labelled tabersonine.

**Fig. S18** Overexpression of TiPS and TiT16H in *C. roseus* flowers.

**Fig. S19** Overexpression of TiPS and TiT16H in *C. roseus* flowers with ^13^C-labeled tabersonine.

**Fig. S20** Percentage of identity shared by TiPS and TiT16H with *C. roseus* P450s.

**Fig. S21** Sequence alignment of TiPS and TiT16H.

**Fig. S22** Transcriptomics and metabolomics related to 16OH-pachysiphine biosynthesis in *T. iboga*.

**Fig. S23** LC-MS profiles and MS^2^ spectra of authentic standards.

**Table S1** Nucleotide sequences of genes described in this study.

**Table S2** Primers used in this study.

## Notes

### Competing Interest Statement

The authors have declared no competing interest.

