## Supporting Information for "Streamlined screening platforms lead to the discovery of pachysiphine synthase from *Tabernanthe iboga*"

The following Supporting Information is available for this article:

**Fig. S1** Design of multi-transcriptional unit constructs

**Fig. S2** *N. benthamiana* leaf disk assay workflow

**Fig. S3** Testing multi-gene constructs in *N. benthamiana.*

**Fig. S4** Chiral LC-MS analysis of pseudo-tabersonine.

**Fig. S5** Phylogenetic tree of *Tabernanthe iboga* (Ti) cytochrome P450s.

**Fig. S6** Iboga CYP71s assayed with (+)-pseudo-tabersonine in *N. benthamiana*.

**Fig. S7** HPLC isolation of hydroxylated tabersonine products from yeast workup.

**Fig. S8** Structure elucidation of (–)-16OH-tabersonine by NMR.

**Fig. S9** $MS^2$ spectral comparison of lochnericine and CL9160.contig1 product.

**Fig. S10** Structure elucidation of (–)-pachysiphine by NMR.

**Fig. S11** Electronic circular dichroism (ECD) spectra of (–)-pachysiphine.

**Fig. S12** Structure elucidation of (–)-16OH-pachysiphine by NMR.

**Fig. S13** LC-MS metabolite profiles of *T. iboga* tissues.

**Fig. S14** MIA precursors present in *C. roseus* var. LBE flowers.

**Fig. S15** Derivatization of tabersonine by endogenous enzymes present in *C. roseus* flowers.

**Fig. S16** Structure confirmation of (–)-[$^{13}$C]-tabersonine.

**Fig. S17** Incubation of *C. roseus* flowers with $^{13}$C-labelled tabersonine.

**Fig. S18** Overexpression of TiPS and TiT16H in *C. roseus* flowers.

**Fig. S19** Overexpression of TiPS and TiT16H in *C. roseus* flowers with $^{13}$C-labeled tabersonine.

**Fig. S20** Percentage of identity shared by TiPS and TiT16H with *C. roseus* P450s.

**Fig. S21** Sequence alignment of TiPS and TiT16H.

**Fig. S22** Transcriptomics and metabolomics related to 16OH-pachysiphine biosynthesis in *T. iboga*.

**Fig. S23** LC-MS profiles and $MS^2$ spectra of authentic standards.

**Table S1** Nucleotide sequences of genes described in this study.

**Table S2** Primers used in this study.

**Methods S1** Click here to enter text.

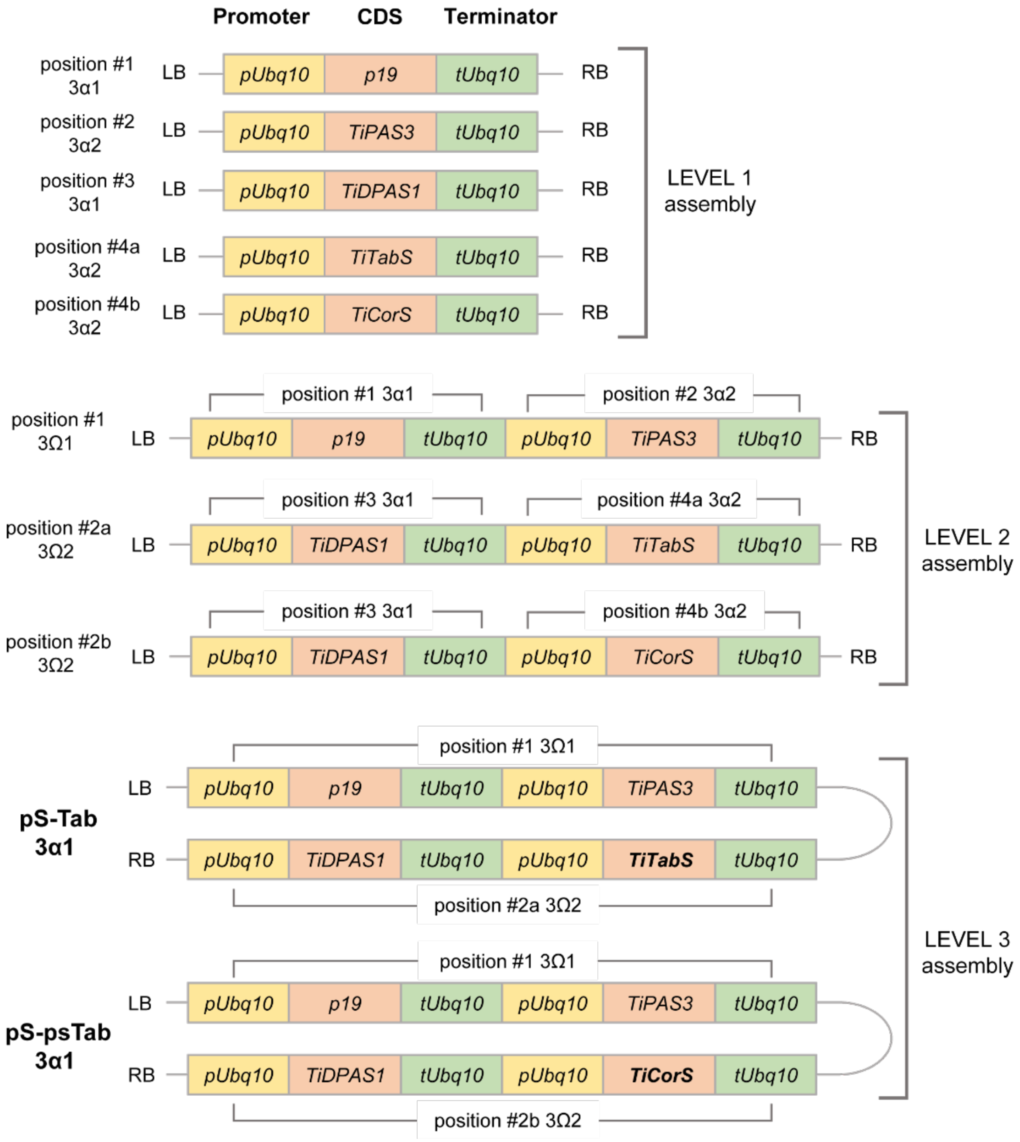


**Fig. S1** Design of multi-transcriptional unit constructs. Overview of the design of Level 1 constructs, and the assembly of Level 2 and Level 3 multi-transcription unit constructs encoding for tabersonine and pseudo-tabersonine biosynthesis for transient expression in *Nicotiana benthamiana*. Transcriptional units were assembled and constructed using the GoldenBraid 2.0 kit. Biosynthetic genes and P19 suppressor are assembled into 3α1 and 3α2 vectors as a transcriptional unit with a Ubiquitin10 promoter (pUbq10) and Ubiquitin10 terminator (tUbq10) at Level 1. Level 1 assemblies are paired into 3Ω1 and 3Ω2 vectors as Level 2 multi-transcriptional unit (multigene) constructs. Level 2 constructs are combined into the 3α1 vector to generate the final multi-gene (4 gene) pS-Tab and pS-psTab, encoding for transient gene expression of tabersonine and pseudo-tabersonine biosynthesis respectively, in *N. benthamiana*. LB and RB indicate the left and right border of the binary plasmid.

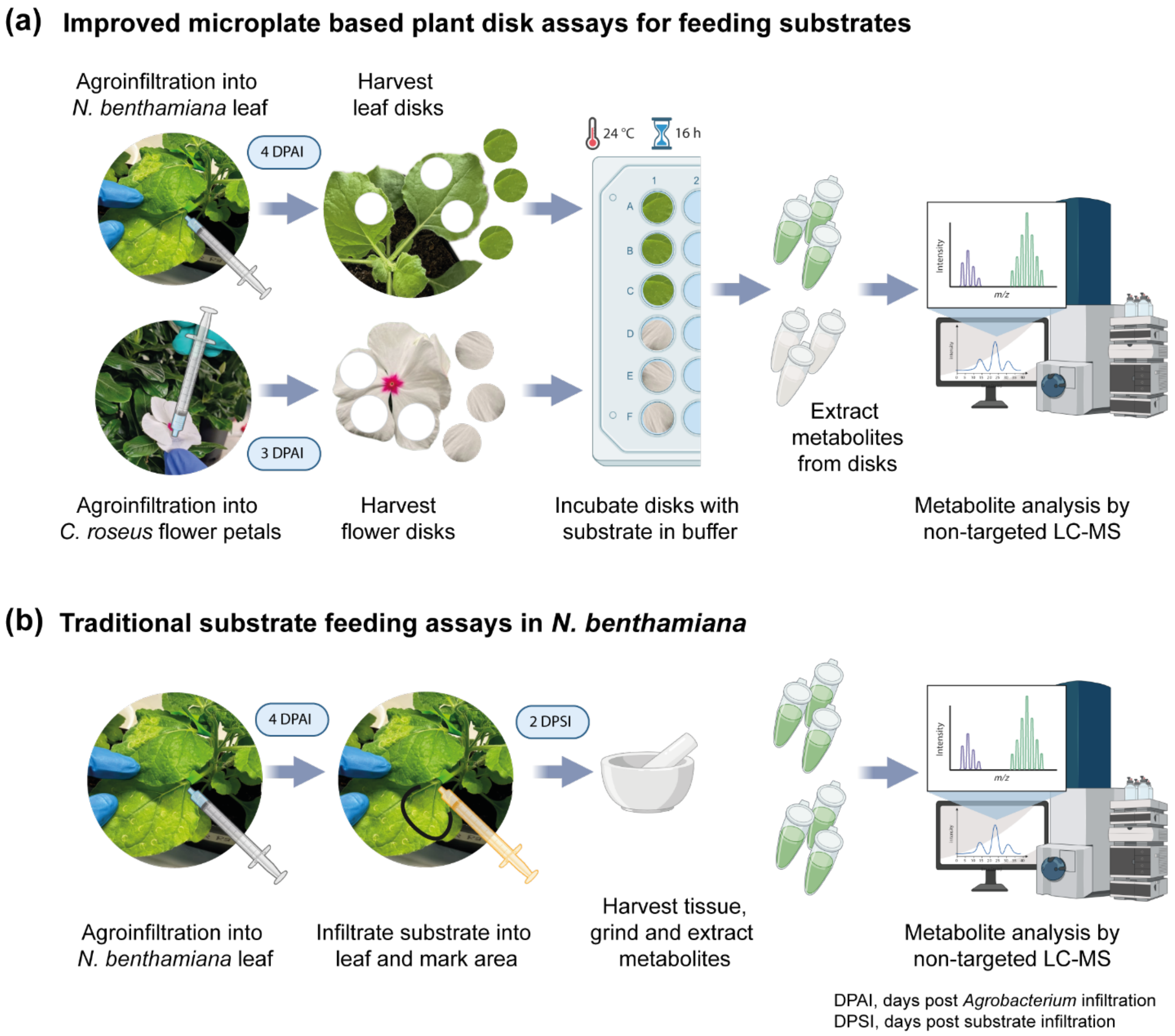


**Fig. S2** *N. benthamiana* leaf disk assay workflow. (a) Schematic of the plant disk-based screening system designed to test for enzyme activity of transiently expressed genes in *N. benthamiana* leaves or *Catharanthus roseus* flowers with exogenous substrate(s). Plant disks are harvested four days post *Agrobacterium* infiltration (DPAI) and incubated in a microplate containing buffer and substrate of known volume and concentration. After overnight incubation, the disks are collected, pulverized, and extracted for LC-MS analysis. (b) Workflow of conventional enzyme activity screening involving syringe-based substrate feeding in *N. benthamiana*. Drawbacks of this workflow include the requirement for larger quantities of substrate, wastage during syringe infiltration, and the lack of precise control over substrate delivery into the plant tissue.

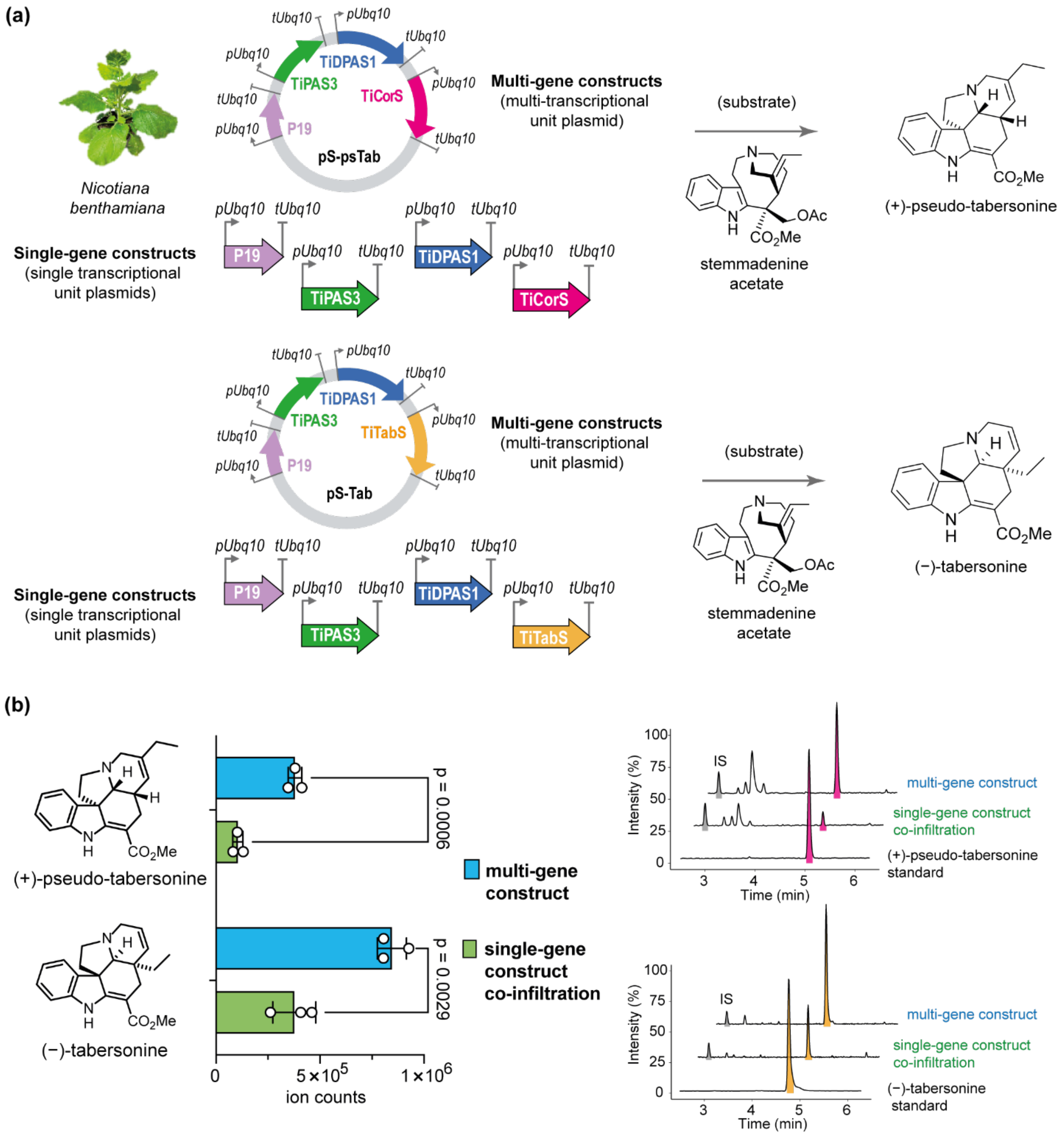


**Fig. S3** Testing multi-gene constructs in *N. benthamiana*. (a) Design of multi-gene constructs pS-Tab and pS-psTab to produce tabersonine and pseudo-tabersonine, respectively, through transient expression in *N. benthamiana* and substrate feeding with stemmadenine acetate. (b) Comparison of the qualitative biosynthetic yields of tabersonine and pseudo-tabersonine produced by the multi-gene constructs (blue bars) versus the equivalent single-gene co-infiltrated constructs (green bars). Representative LC-MS profiles (total ion chromatograms) show the production of tabersonine and pseudo-tabersonine qualitatively, with ajmaline as the internal standard (IS). The values represent mean ± SD (n = 3), two-tailed Student's t-test.

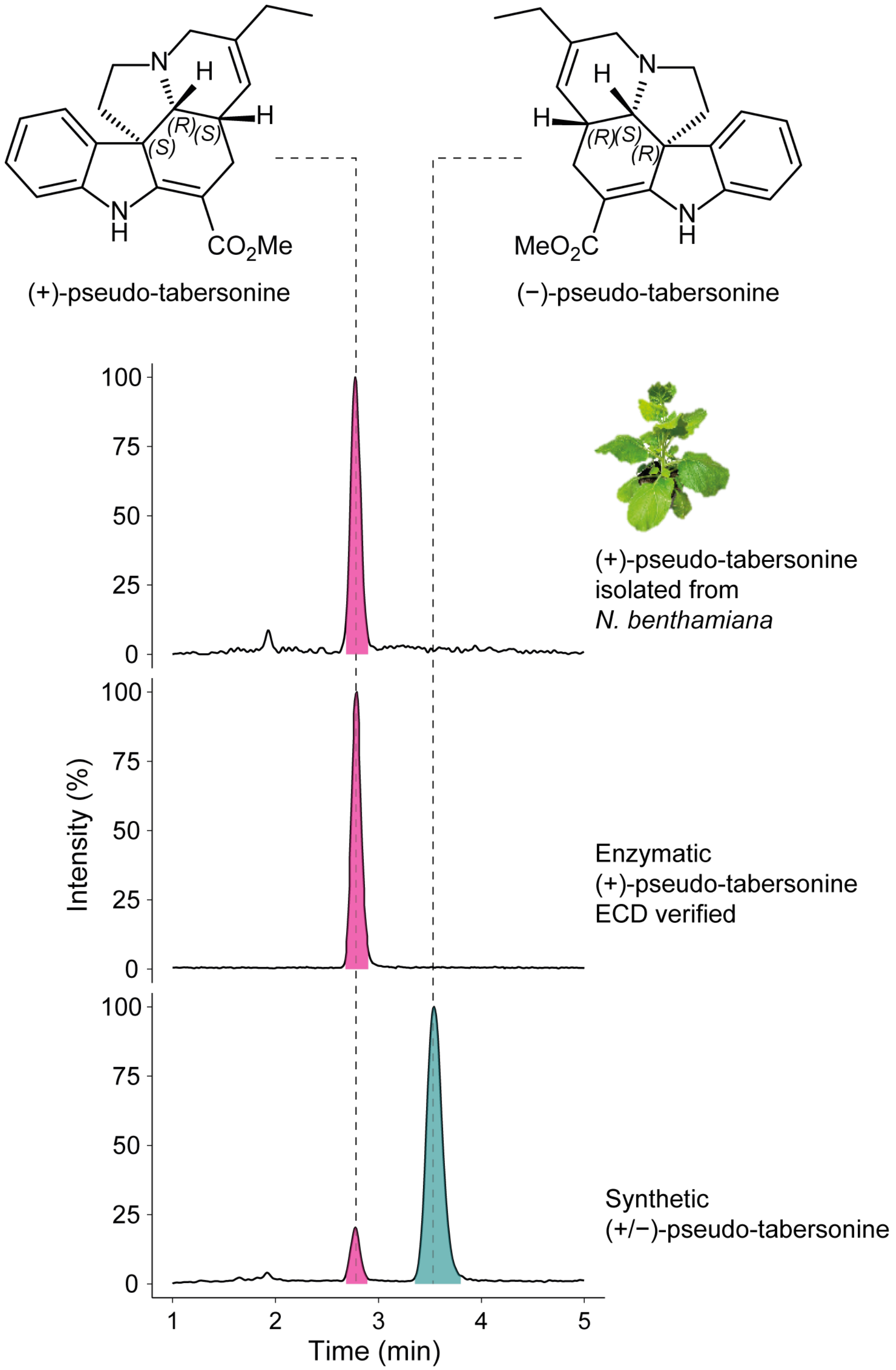


**Fig. S4** Chiral LC-MS analysis of pseudo-tabersonine. The biosynthetic product of pseudo-tabersonine generated in *N. benthamiana* is enantiomerically pure (+)-pseudo-tabersonine. LC-MS traces are presented as total ion chromatograms (solid lines).

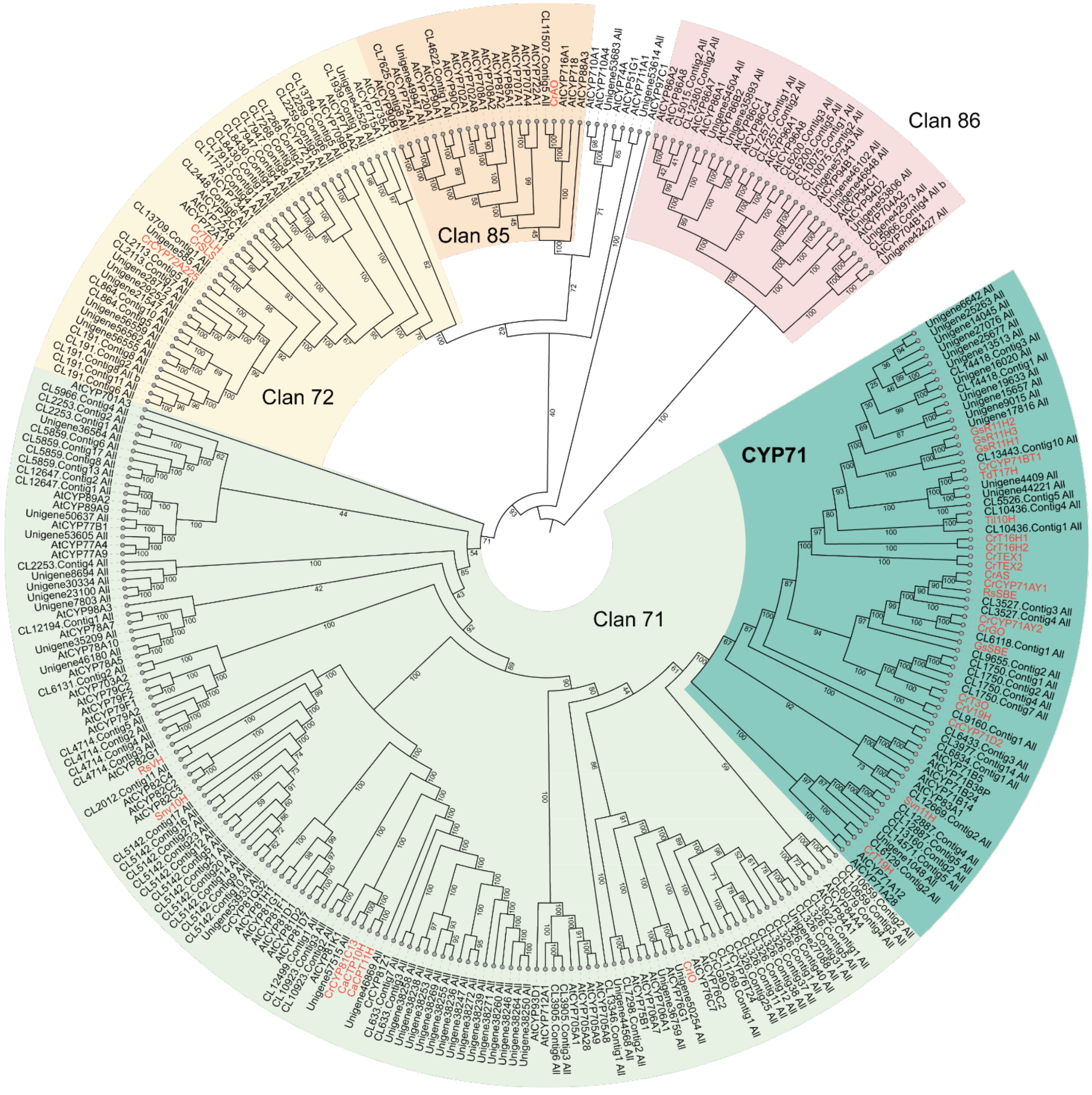


**Fig. S5** Phylogenetic tree of *Tabernanthe iboga* (Ti) cytochrome P450s. Full-length P450s were extracted from de novo assembled transcriptome. Selected *Arabidopsis thaliana* (At) CYPs were included in the analysis to define CYP clans and CYP71 based on (Hansen *et al.*, 2021). The maximum-likelihood phylogenetic tree was constructed using iQtree (Trifinopoulos *et al.*, 2016) with default parameters and 1000 bootstraps (shown out of 100). The CYP71 family and major P450 clans are highlighted and labelled. P450s functionally characterized to be involved in monoterpene indole alkaloid (MIA) biosynthesis are coloured in red. Clan 86 was used to root the tree. The phylogenetic tree was visualized and inferred using iToL (Letunic & Bork, 2024).

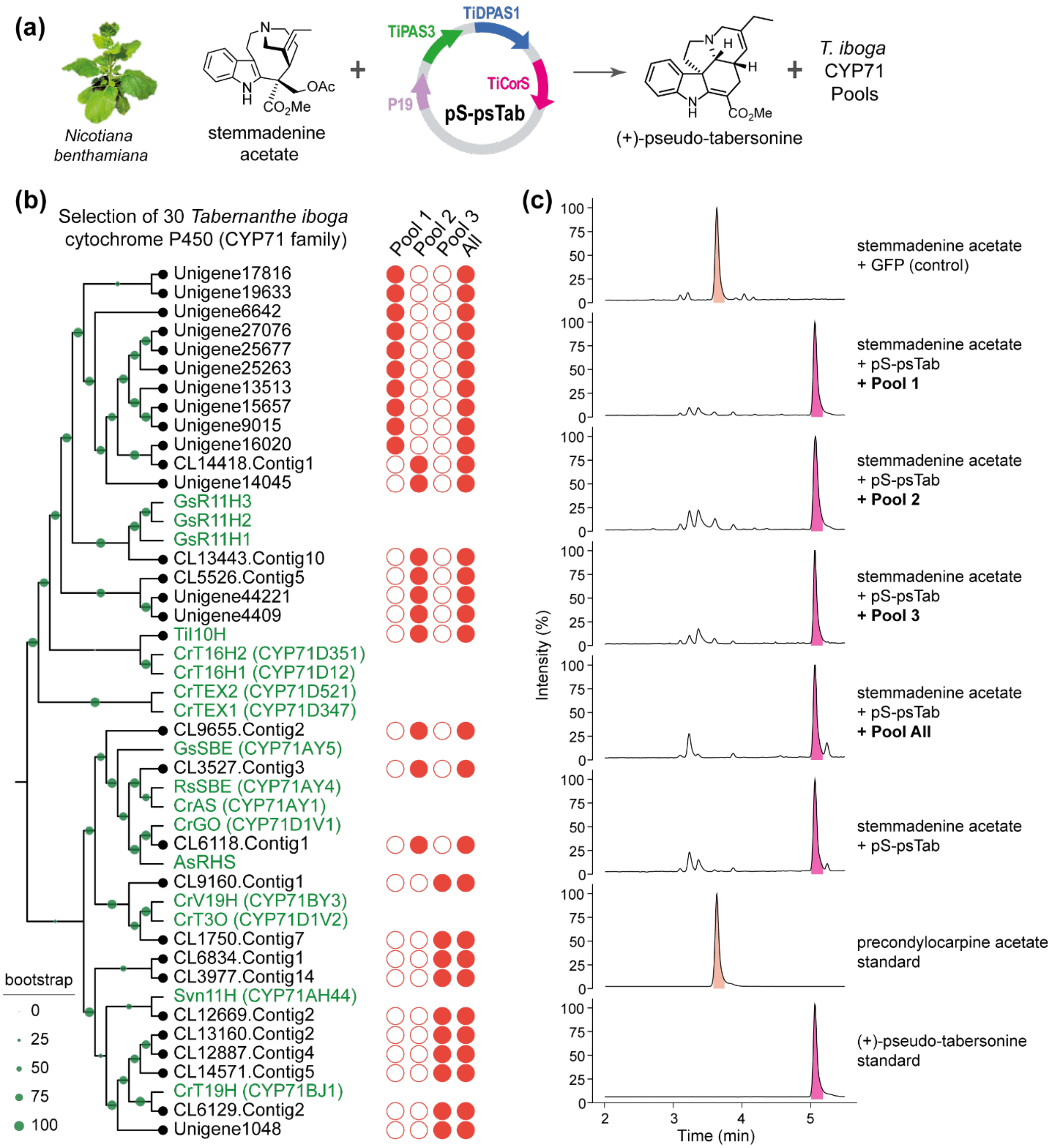


**Fig. S6** Iboga CYP71s assayed with (+)-pseudo-tabersonine in *N. benthamiana*. (a) Assay plan for screening CYP71 gene candidate pools with the (+)-pseudo-tabersonine producing multigene construct pS-psTa. (b) Phylogenetic tree of the iboga CYP71s and the candidate pools selected for screening in *N. benthamiana*. (c) LC-MS analysis of the iboga CYP71 candidate pools tested for activity in *N. benthamiana* with the multigene construct pS-psTab. LC-MS traces are presented as total ion chromatograms (solid lines). Stemmadenine acetate is oxidized to precondylocarpine acetate by an endogenous *N. benthamiana* enzyme.

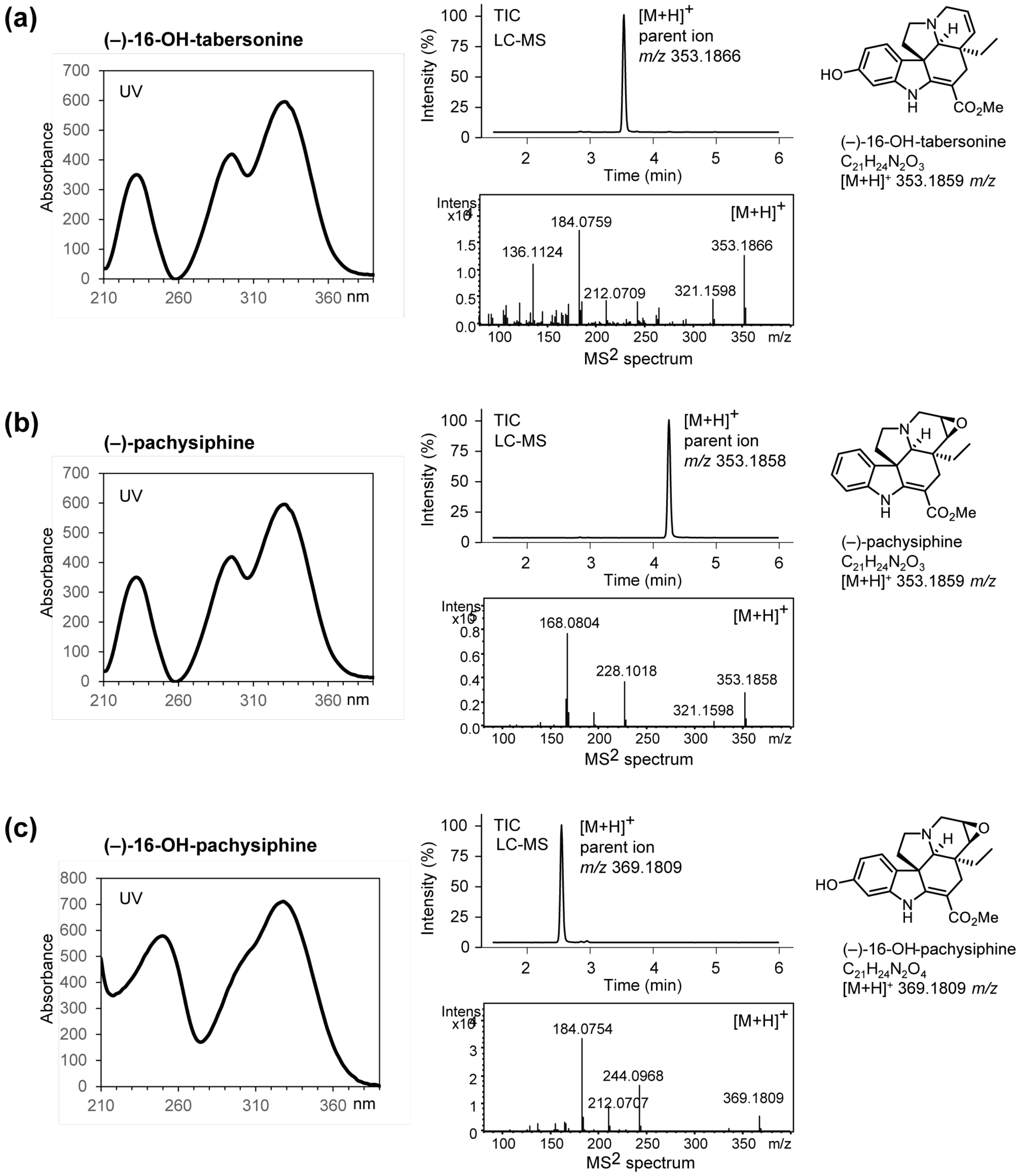


**Fig. S7** HPLC isolation of hydroxylated tabersonine products from yeast workup. (a) Isolated product (–)-16OH-tabersonine. (b) Isolated product (–)-pachysiphine. (c) Isolated product (–)-16OH-pachysiphine. Traces indicate UV spectrum, LC-MS chromatography profile, and $MS^2$ spectrum of parent ions of the isolated compounds. LC-MS traces are presented as total ion chromatograms.

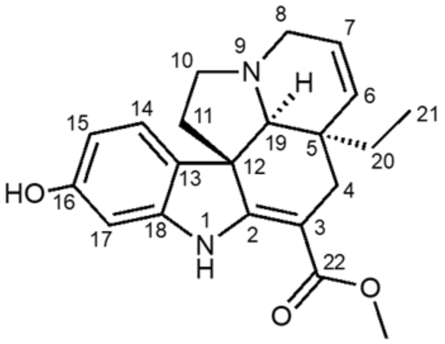


(–)-16-OH-tabersonine
$^{1}$H-NMR, MeOH-$d_3$
700 MHz, 25 °C

| pos. | $\delta_H$ | mult. | $J_{HH}$ | $\delta_C$ |
|---|---|---|---|---|
| 1 | 9.23 | *bs* | - | - |
| 2 | - | - | - | 168.1 |
| 3 | - | - | - | 91.5 |
| 4α | 2.58 | *bd* | 15.4 | 29.7 |
| 4β | 2.40 | *d* | 15.4 | 29.7 |
| 5 | - | - | - | 42.5 |
| 6α | 5.75 | *bd* | 10.0 | 134.4 |
| 7α | 5.82 | *bdd* | 10.0/4.5 | 125.4 |
| 8α | 3.34 | *bd* | 16.0 | 51.5 |
| 8β | 3.52 | *dd* | 16.0/4.5 | 51.5 |
| 10α | 2.85 | *m*** | - | 51.9 |
| 10β | 3.10 | *dd* | 7.5/6.4 | 51.9 |
| 11α | 1.78 | *dd* | 11.8/4.4 | 45.8 |
| 11β | 2.00 | *ddd* | 11.8/11.6/6.4 | 45.8 |
| 12 | - | - | - | 55.8 |
| 13 | - | - | - | 129.6 |
| 14 | 7.08 | *d* | 8.0 | 122.8 |
| 15 | 6.30 | *dd* | 8.0/1.8 | 108.2 |
| 16 | - | - | - | 159.3 |
| 17 | 6.45 | *d* | 1.8 | 99.2 |
| 18 | - | - | - | 146.1 |
| 19α | 2.86 | *bs* | - | 71.6 |
| 20a | 0.92 | *dq* | 14.5/7.4 | 28.0 |
| 20b | 1.03 | *dq* | 14.5/7.4 | 28.0 |
| 21 | 0.65 | *t* | 7.4 | 7.9 |
| 22 | - | - | - | 170.0 |
| OMe | 3.75 | *s* | - | 51.6 |

** overlapped signals J unresolved

**Fig. S8** Structure elucidation of (–)-16OH-tabersonine by NMR. (Continued to next page)

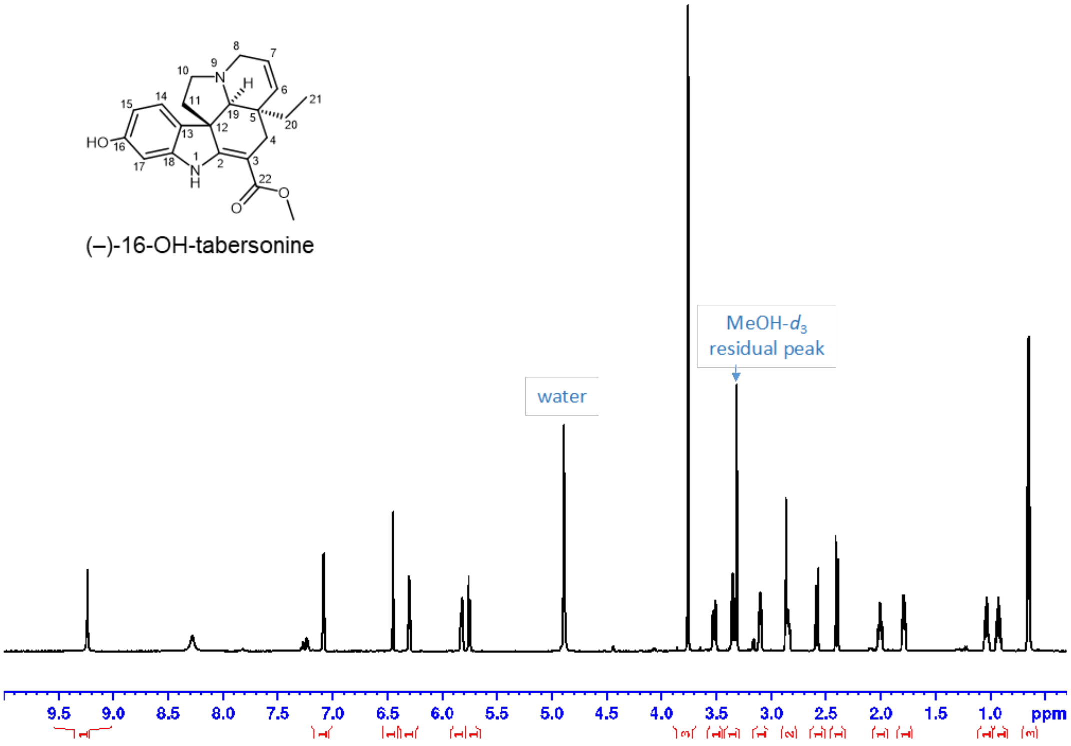


$^1$H NMR with water suppression (zgpr) full range in MeOH-$d_3$

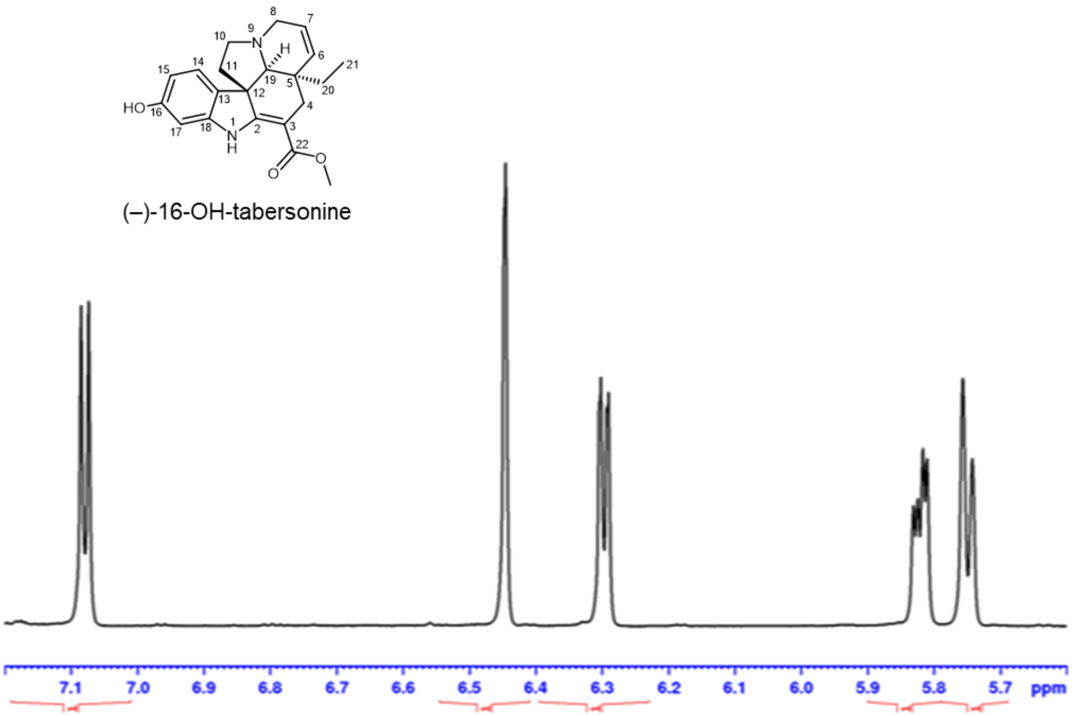


$^1$H NMR with water suppression (zgpr)
5.6-7.2 ppm in MeOH-$d_3$

**Fig. S8** Structure elucidation of (–)-16OH-tabersonine by NMR. (Continued to next page).

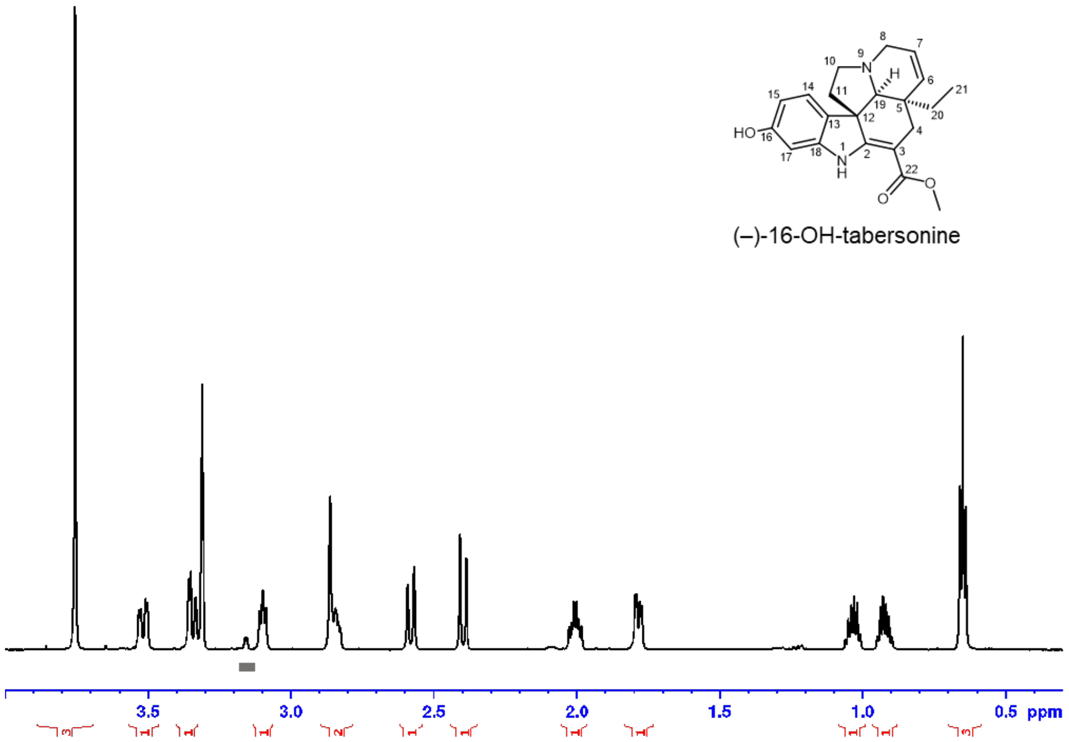


**$^1$H NMR with water suppression (zgpr)**
aliphatic range in MeOH-$d_3$. Shaded areas mark impurity

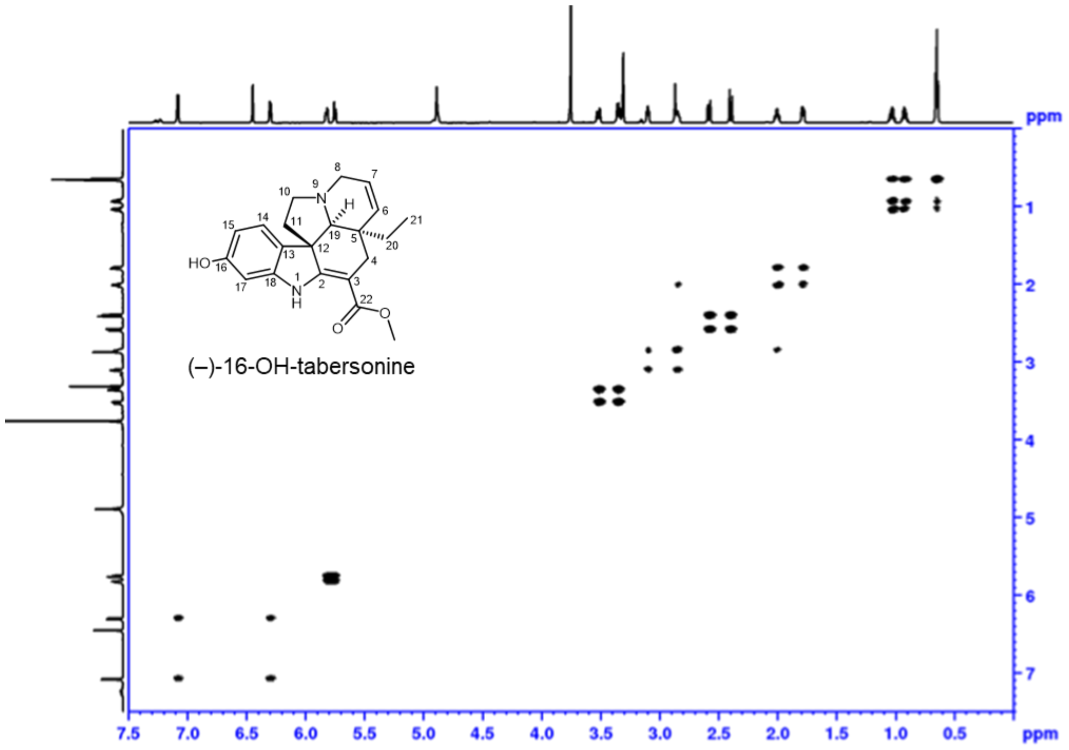


**$^1$H-$^1$H DQF COSY** with water suppression
full range in MeOH-$d_3$

**Fig. S8** Structure elucidation of (–)-16OH-tabersonine by NMR. (Continued to next page).

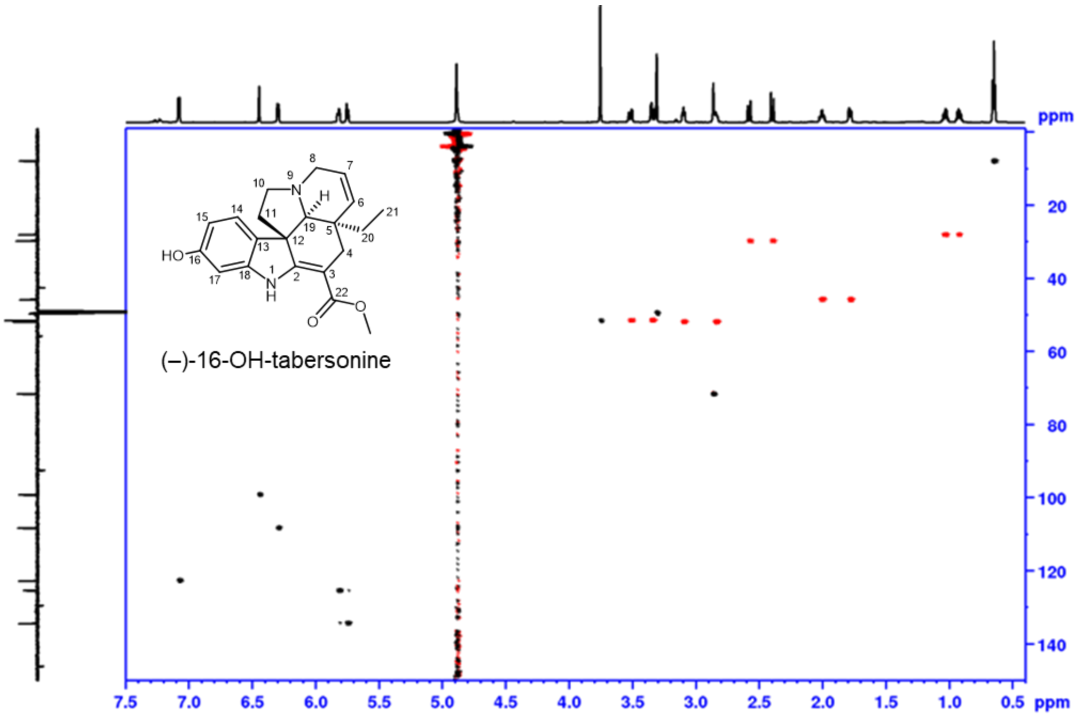


**phase sensitive HSQC**, full range in MeOH-$d_3$
red: $CH_2$, black: CH, $CH_3$

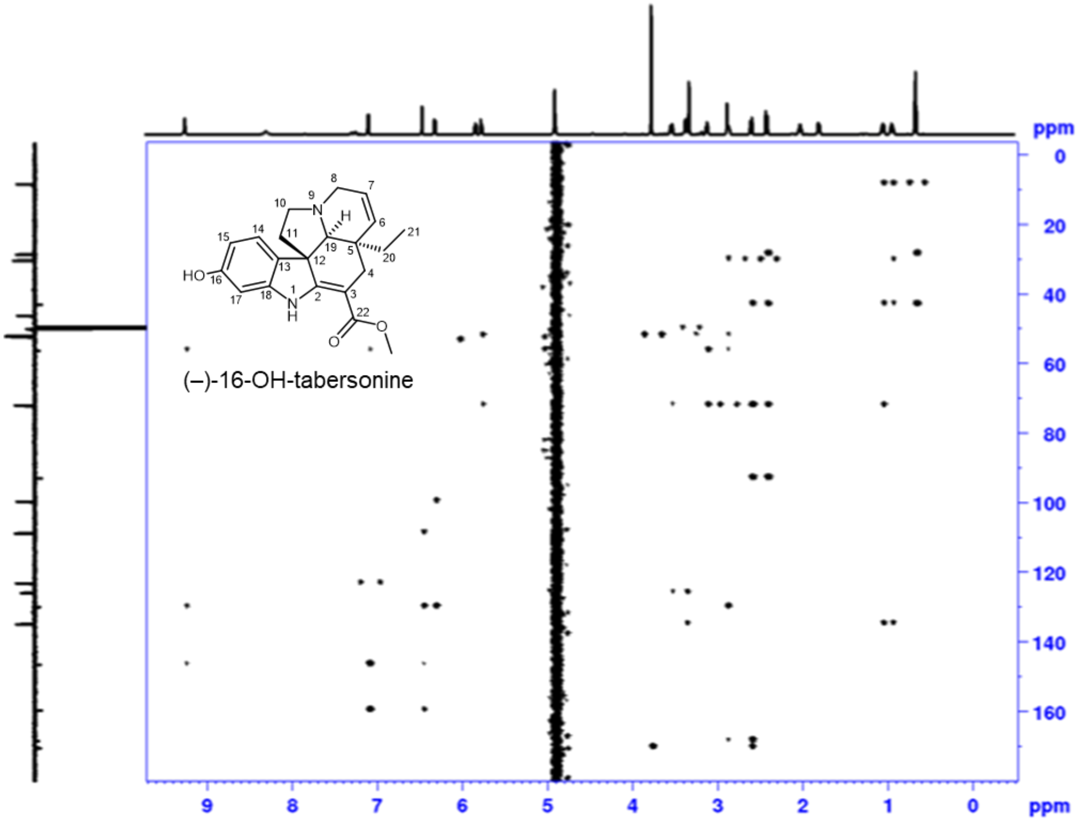


**HMBC**, full range in MeOH-$d_3$

**Fig. S8** Structure elucidation of (–)-16OH-tabersonine by NMR. (Continued to next page).

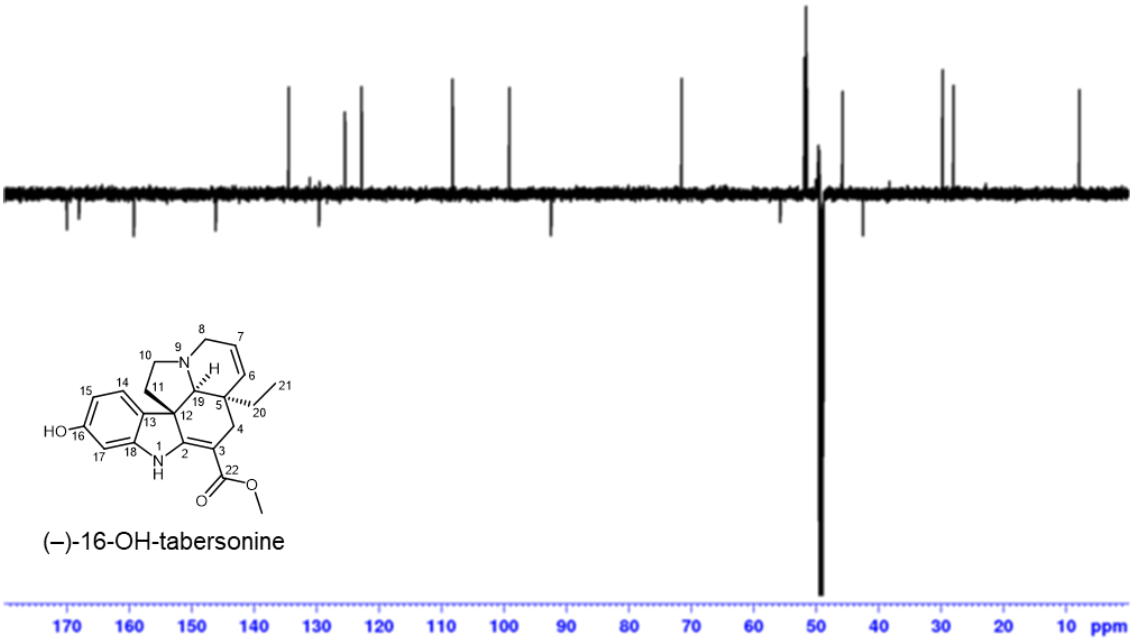


**DEPTQ**, full range in MeOH-$d_3$

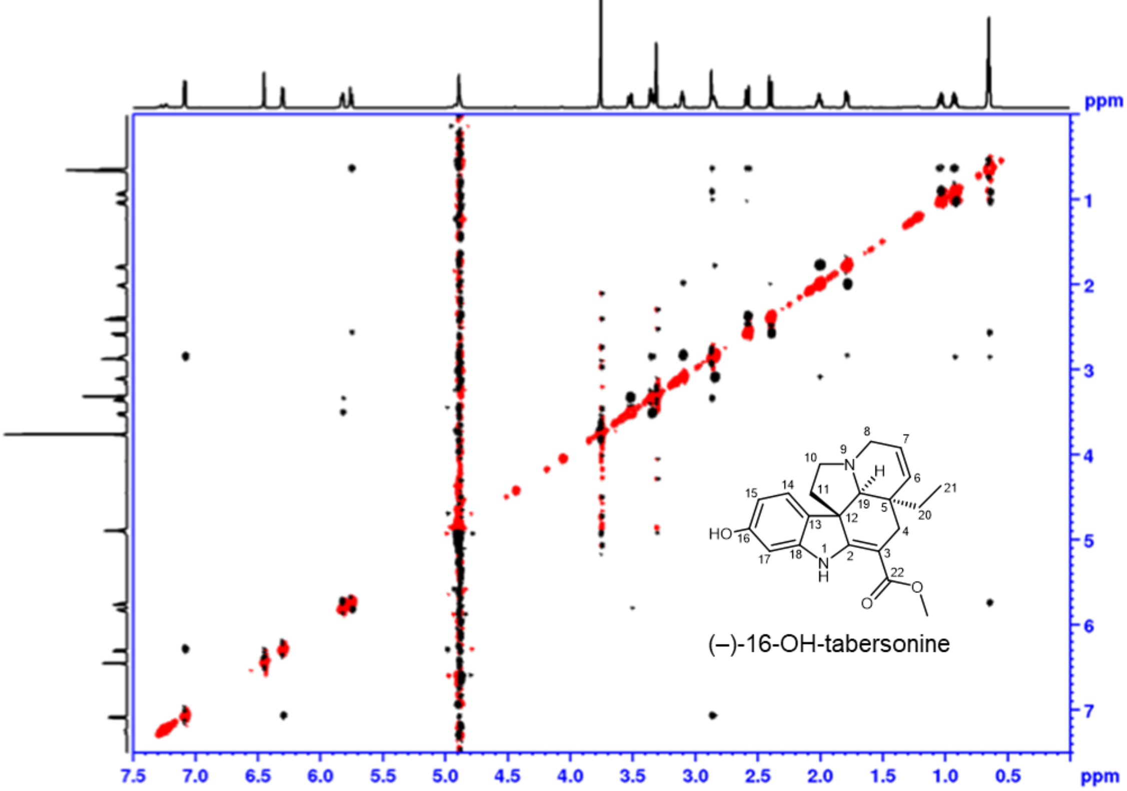


**$^1$H-$^1$H ROESY with water suppression**
full range in MeOH-$d_3$

**Fig. S8** Structure elucidation of (–)-16OH-tabersonine by NMR. (End).

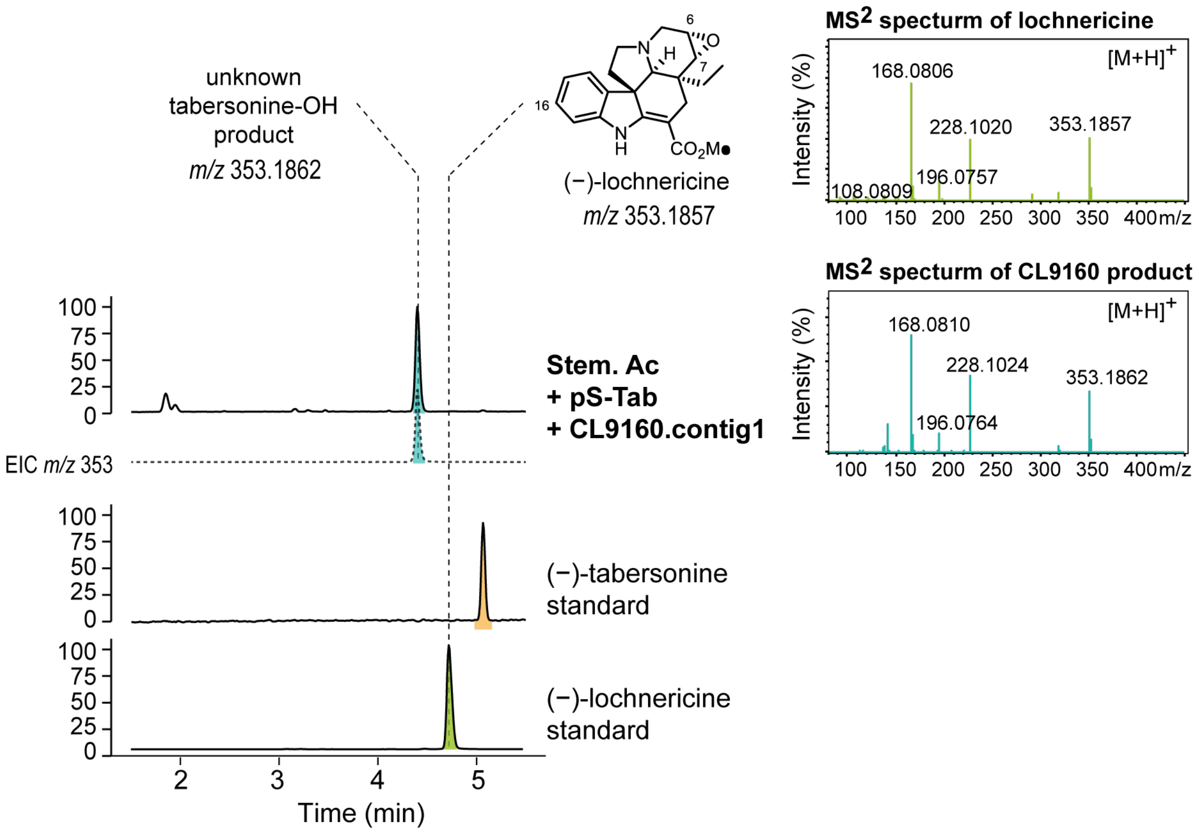


**Fig. S9** $MS^2$ spectral comparison of lochnericine and CL9160.contig1 product. LC-MS profiles and $MS^2$ spectra of lochnericine standard and the hydroxylated tabersonine product of CL9160.contig1. Comparison of $MS^2$ spectra of the parent ions *m/z* 353.18 shows similar fragmentation patterns between the two compounds, suggesting structural similarities.

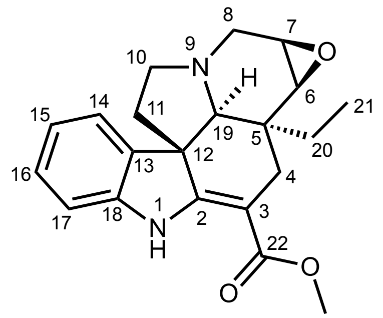


(–)-pachysiphine
[1]H-NMR, $CDCl_3$
700 MHz, 25 °C

| pos. | $\delta_H$ | mult. | $J_{HH}$ | $\delta_C$ |
|---|---|---|---|---|
| 1 | 8.97 | *bs* | - | - |
| 2 | - | - | - | 165.5 |
| 3 | - | - | - | 91.4 |
| 4α | 2.67 | *dd* | 15.4/1.6 | 23.6 |
| 4β | 2.52 | *d* | 15.4 | 23.6 |
| 5 | - | - | - | 37.4 |
| 6α | 3.05 | *d* | 3.9 | 56.6 |
| 7α | 3.24 | *ddd* | 3.9/1.1/0.4 | 52.4 |
| 8α | 2.87 | *bd* | 12.9 | 49.9 |
| 8β | 3.56 | *dd* | 12.9/1.1 | 49.9 |
| 10α | 2.69 | *ddd* | 11.5/8.2/4.4 | 51.4 |
| 10β | 3.01 | *dd* | 8.2/6.5 | 51.4 |
| 11α | 1.68 | *dd* | 11.6/4.4 | 44.3 |
| 11β | 2.06 | *ddd* | 11.6/11.5/6.5 | 44.3 |
| 12 | - | - | - | 54.9 |
| 13 | - | - | - | 137.9 |
| 14 | 7.15 | *bd* | 7.4 | 121.7 |
| 15 | 6.85 | *ddd* | 7.4/7.4/0.9 | 120.8 |
| 16 | 7.13 | *ddd* | 7.7/7.4/1.0 | 128.1 |
| 17 | 6.79 | *bd* | 7.7 | 109.6 |
| 18 | - | - | - | 143.3 |
| 19α | 2.44 | *bd* | 1.6 | 71.4 |
| 20a | 0.91 | *dq* | 14.7/7.4 | 26.7 |
| 20b | 0.96 | *dq* | 14.7/7.4 | 26.7 |
| 21 | 0.71 | *t* | 7.4 | 7.4 |
| 22 | - | - | - | 169.2 |
| OMe | 3.76 | *s* | - | 51.3 |

**Fig. S10** Structure elucidation of (–)-pachysiphine by NMR. (Continued to next page)

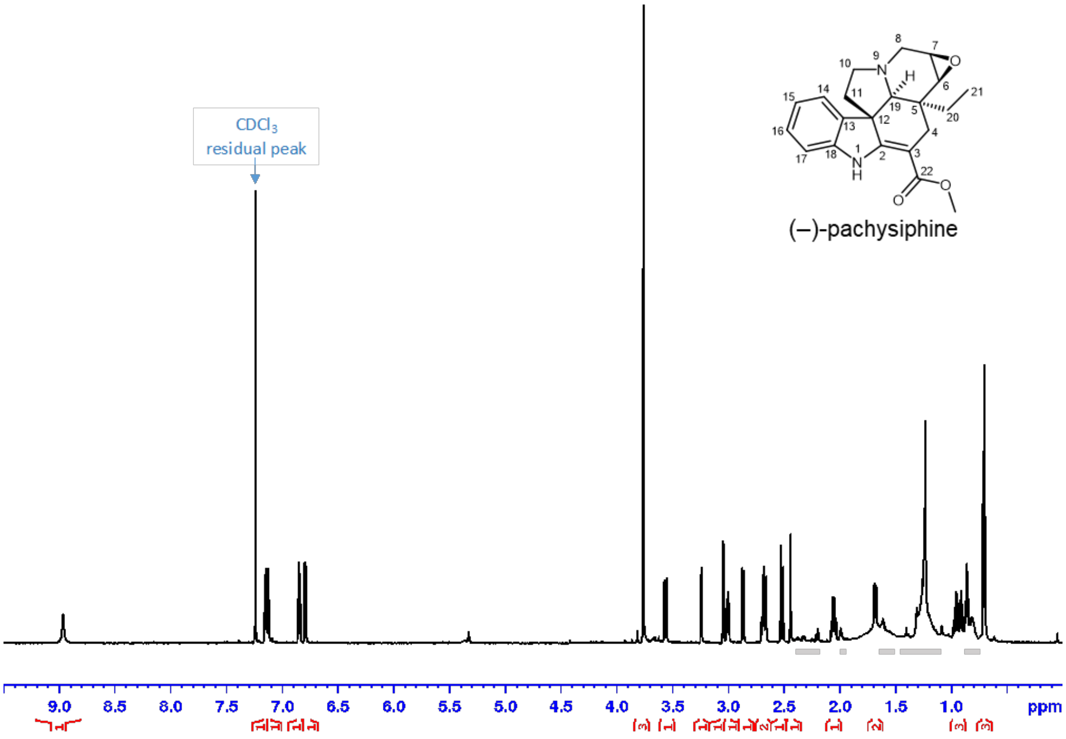


**¹H NMR with water suppression (zgpr)** full range in $CDCl_3$
Shaded areas mark impurity

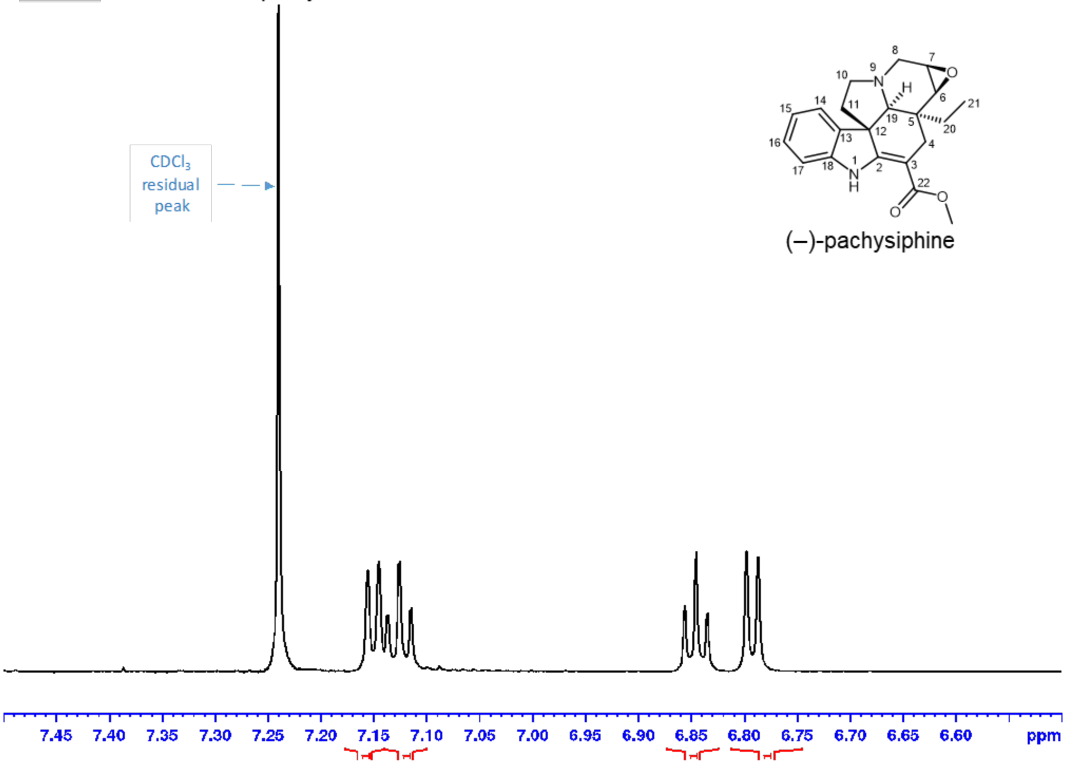


**¹H NMR with water suppression (zgpr)**
aromatic range in CDCl3

**Fig. S10** Structure elucidation of (–)-pachysiphine by NMR. (Continued to next page).

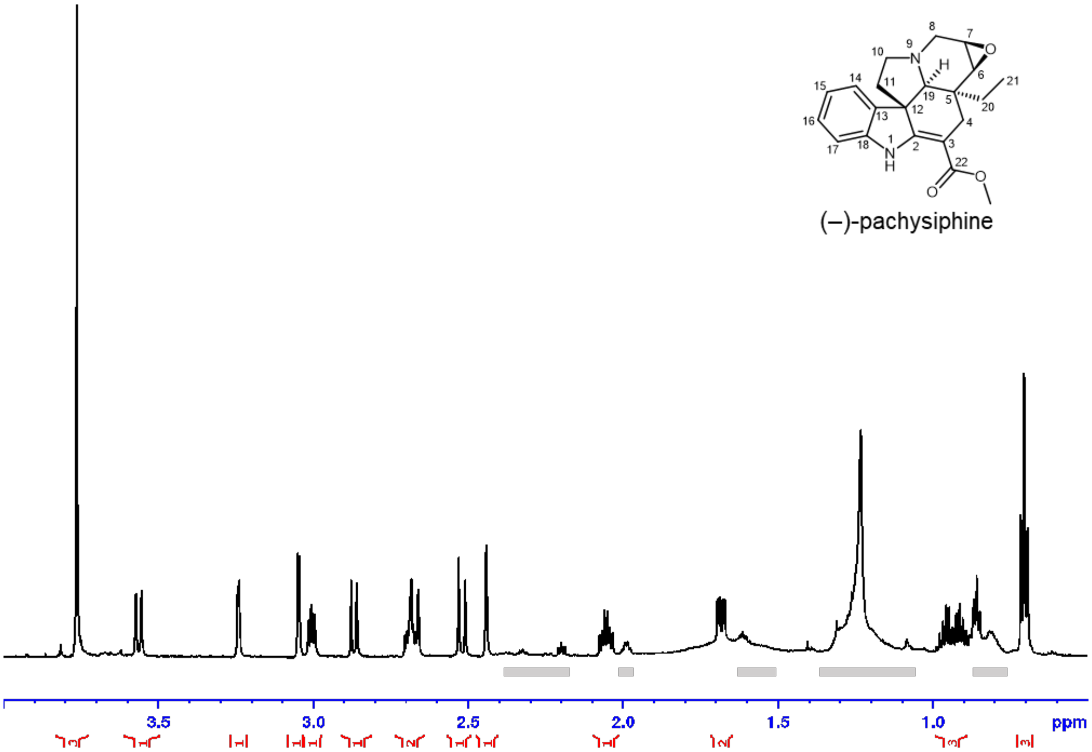


**$^1$H NMR with water suppression (zgpr)**
aliphatic range in $CDCl_3$. Shaded areas mark impurity.

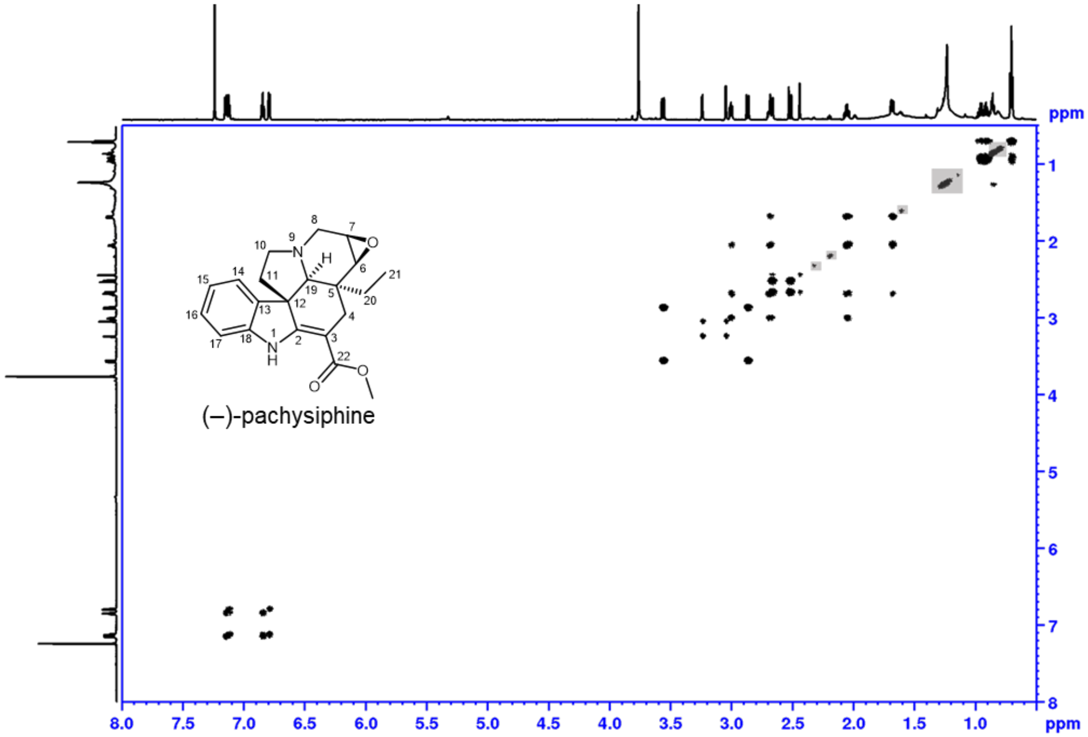


**$^1$H-$^1$H DQF COSY**, full range in $CDCl_3$
Shaded areas mark impurity

**Fig. S10** Structure elucidation of (–)-pachysiphine by NMR. (Continued to next page).

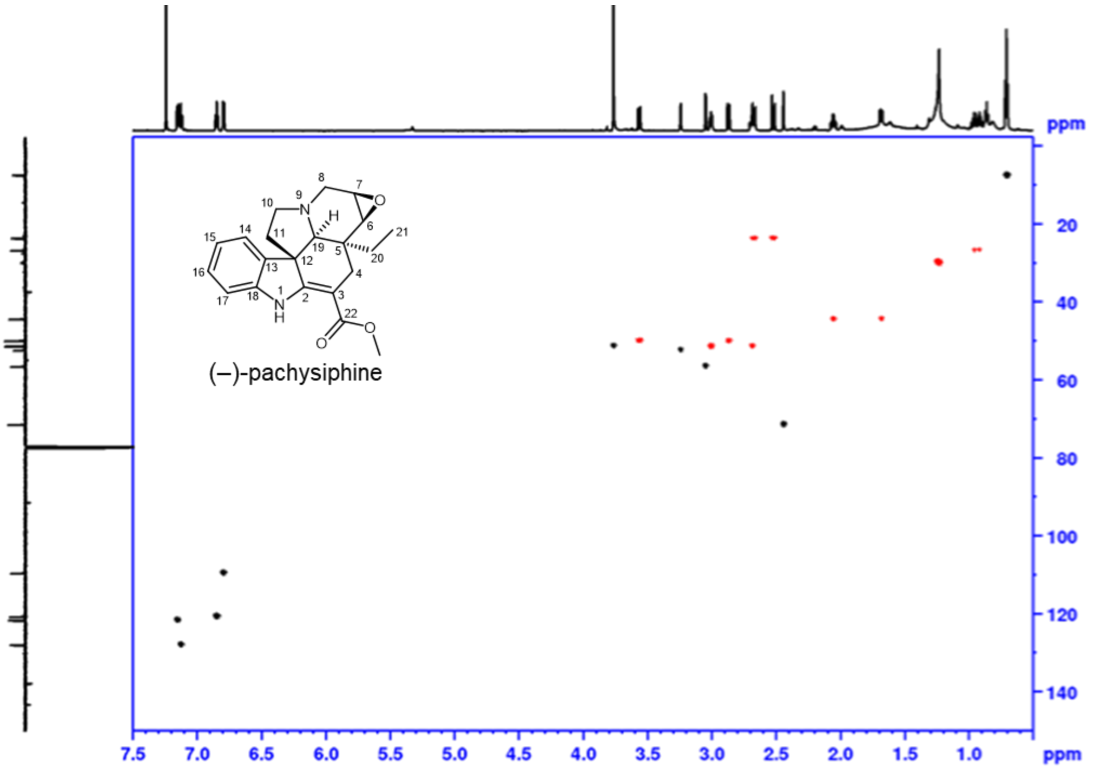


**phase sensitive HSQC**, full range in $CDCl_3$
red: $CH_2$, black: CH, $CH_3$

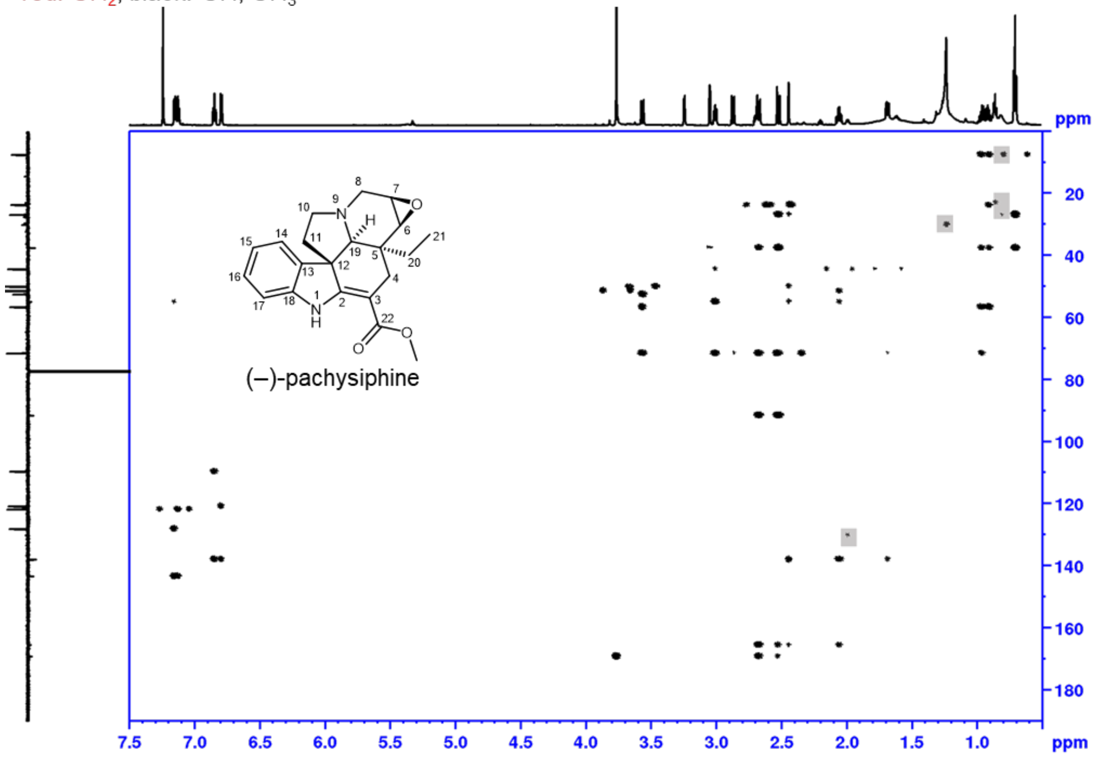


**HMBC**, full range in $CDCl_3$
Shaded areas mark impurity

**Fig. S10** Structure elucidation of (–)-pachysiphine by NMR. (Continued to next page).

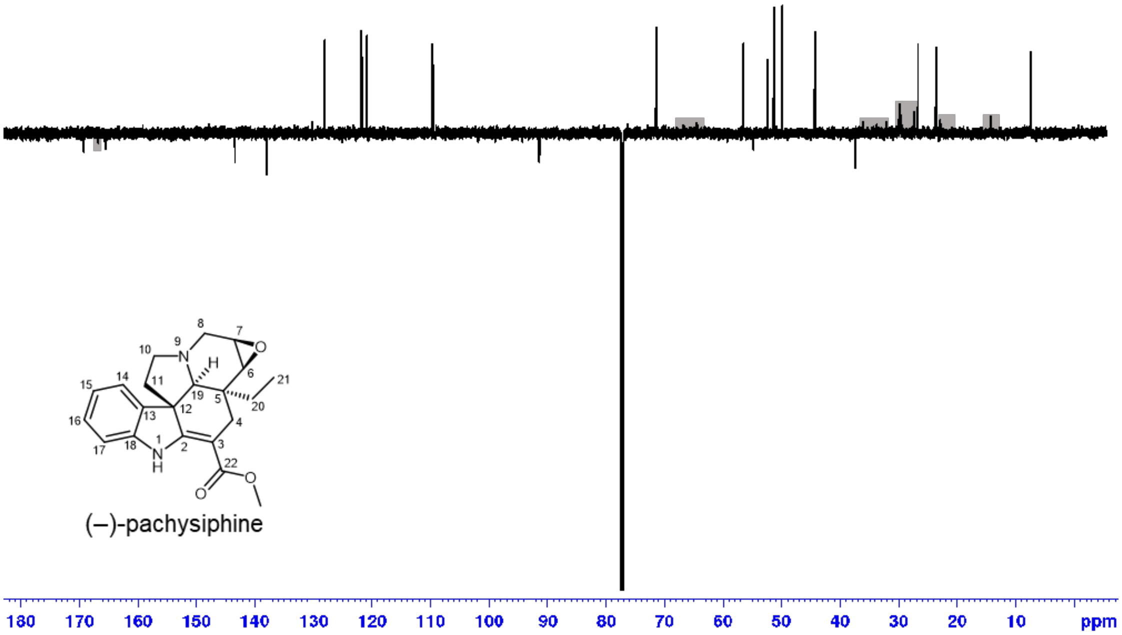


**DEPTQ**, full range in $CDCl_3$
Shaded areas mark impurity

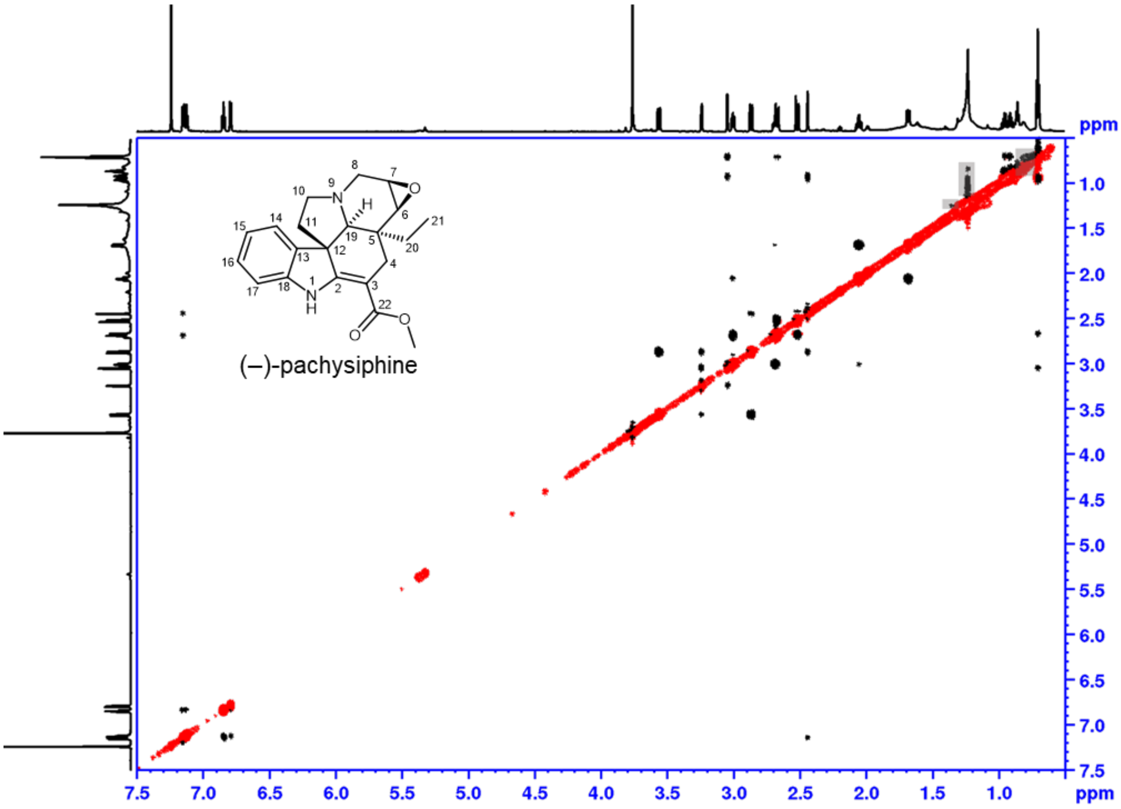


**$^1H$-$^1H$ ROESY with water suppression**
full range in $CDCl_3$. Shaded areas mark impurity

**Fig. S10** Structure elucidation of (–)-pachysiphine by NMR. (Continued to next page).

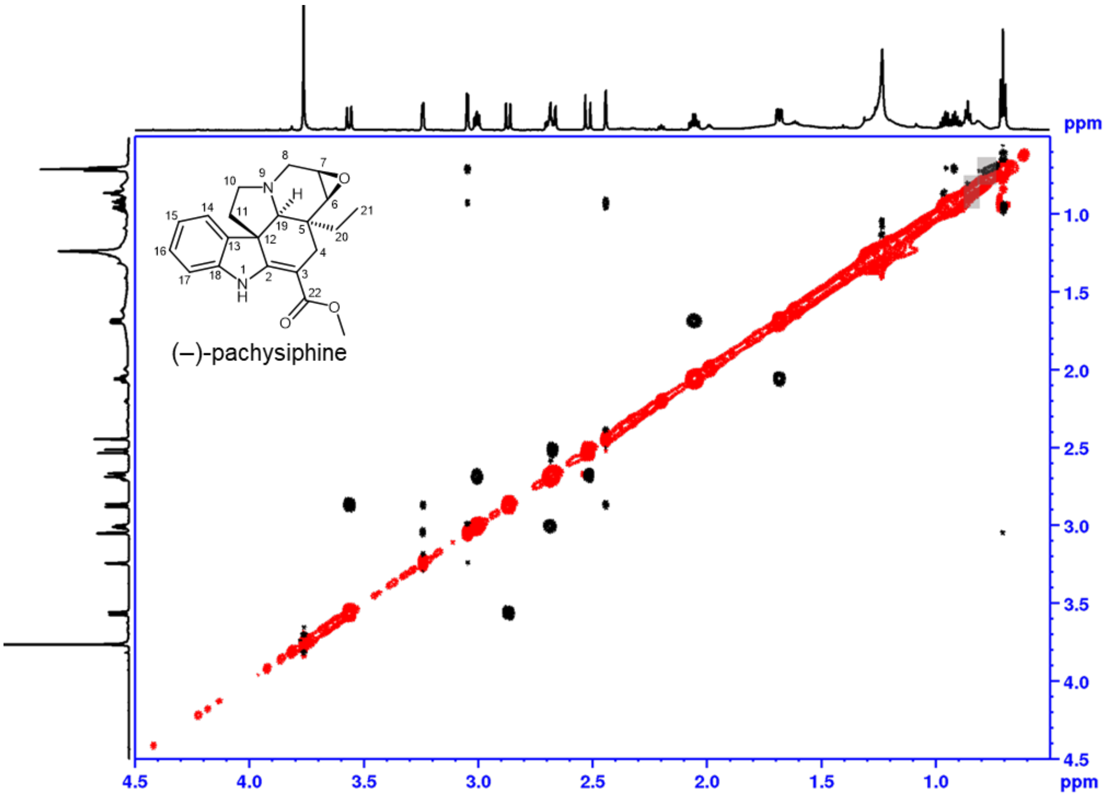


**$^1H$-$^1H$ ROESY with water suppression**
aliphatic range in $CDCl_3$. Shaded areas mark impurity

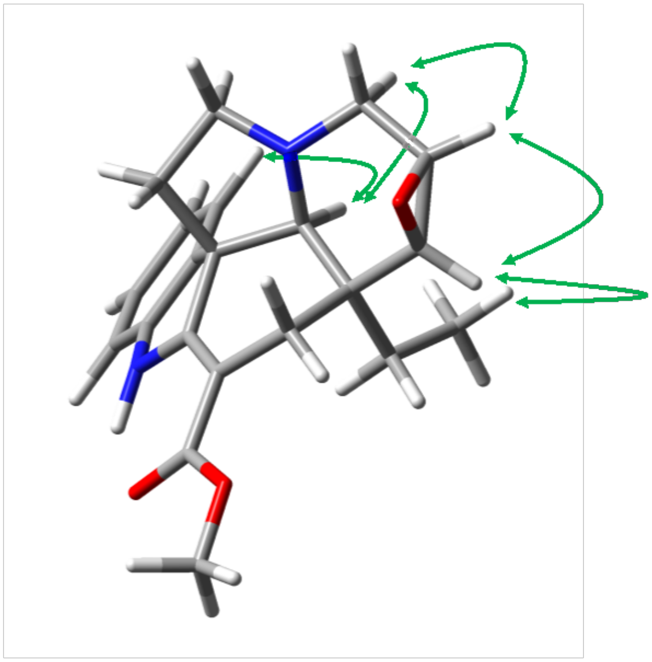

**ROESY correlations of (–)-pachysiphine**

Optimized using Gaussian 16W (DFT, APFD/6-311G+(2d,p), solvent MeOH). Important ROESY correlations are depicted in green.

**Fig. S10** Structure elucidation of (–)-pachysiphine by NMR. (End).

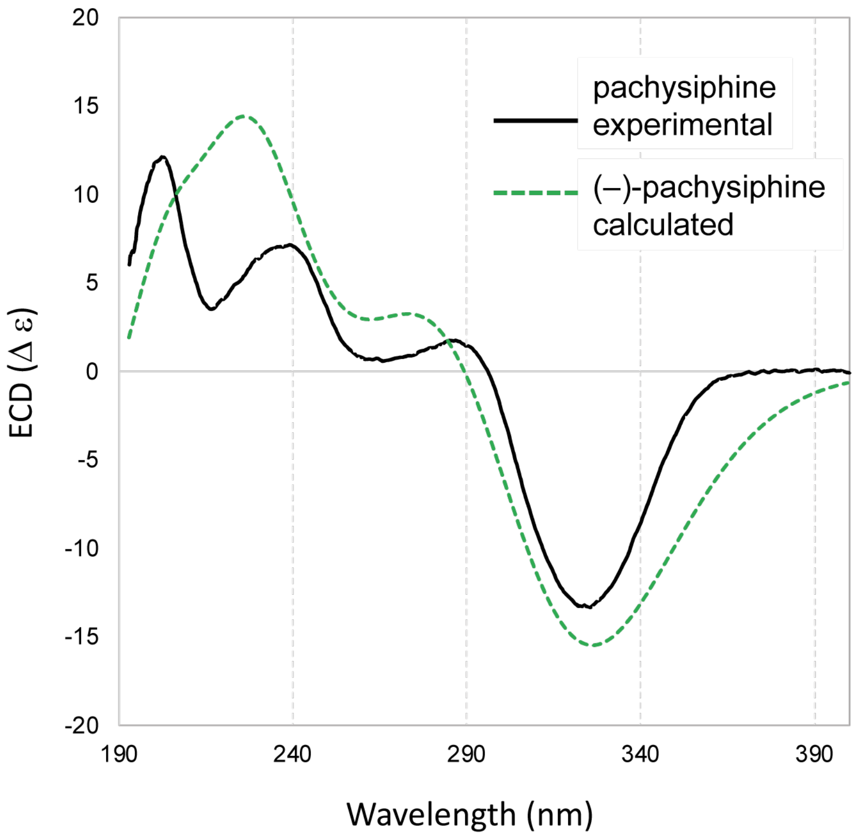


**Fig. S11** Electronic circular dichroism (ECD) spectra of (–)-pachysiphine. Comparison of simulated ECD spectrum (dashed green) and experimental ECD spectrum (solid black) of (–)-pachysiphine.

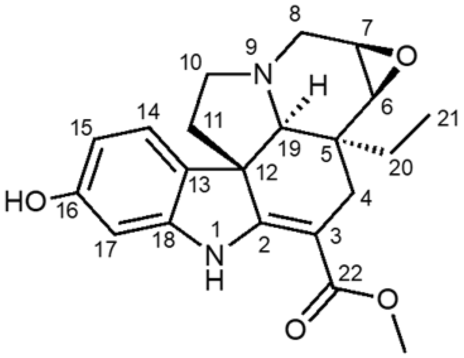


(–)-16-OH-pachysiphine
$^{1}$H-NMR, MeOH-$d_3$
700 MHz, 25 °C

| pos. | $\delta_H$ | mult. | $J_{HH}$ | $\delta_C$ |
|---|---|---|---|---|
| 1 | 9.19 | *bs* | - | - |
| 2 | - | - | - | 167.6 |
| 3 | - | - | - | 91.3 |
| 4α | 2.60 | *dd* | 15.4/1.6 | 24.6 |
| 4β | 2.46 | *d* | 15.4 | 24.6 |
| 5 | - | - | - | 38.5 |
| 6α | 3.06 | *d* | 3.7 | 57.6 |
| 7α | 3.28 | *bd* | 3.7 | 53.6 |
| 8α | 2.88 | *d* | 12.8 | 50.7 |
| 8β | 3.46 | *bd* | 12.8 | 50.7 |
| 10α | 2.68 | *ddd* | 11.4/8.4/4.3 | 51.8 |
| 10β | 2.88 | *dd* | 8.4/6.3 | 51.8 |
| 11α | 1.60 | *dd* | 11.4/4.3 | 45.8 |
| 11β | 1.91 | *ddd* | 11.4/11.4/6.3 | 45.8 |
| 12 | - | - | - | 55.7 |
| 13 | - | - | - | 130.0 |
| 14 | 6.99 | *d* | 8.1 | 122.8 |
| 15 | 6.28 | *dd* | 8.1/2.0 | 108.1 |
| 16 | - | - | - | 159.1 |
| 17 | 6.43 | *d* | 2.0 | 99.1 |
| 18 | - | - | - | 146.0 |
| 19α | 2.45 | *bd* | 1.6 | 72.5 |
| 20a | 0.96 | *q* | 7.3 | 27.3 |
| 20b | 0.96 | *q* | 7.3 | 27.3 |
| 21 | 0.72 | *t* | 7.3 | 7.5 |
| 22 | - | - | - | 170.1 |
| OMe | 3.77 | *s* | - | 51.5 |

**Fig. S12** Structure elucidation of (–)-16OH-pachysiphine by NMR. (Continued to next page)

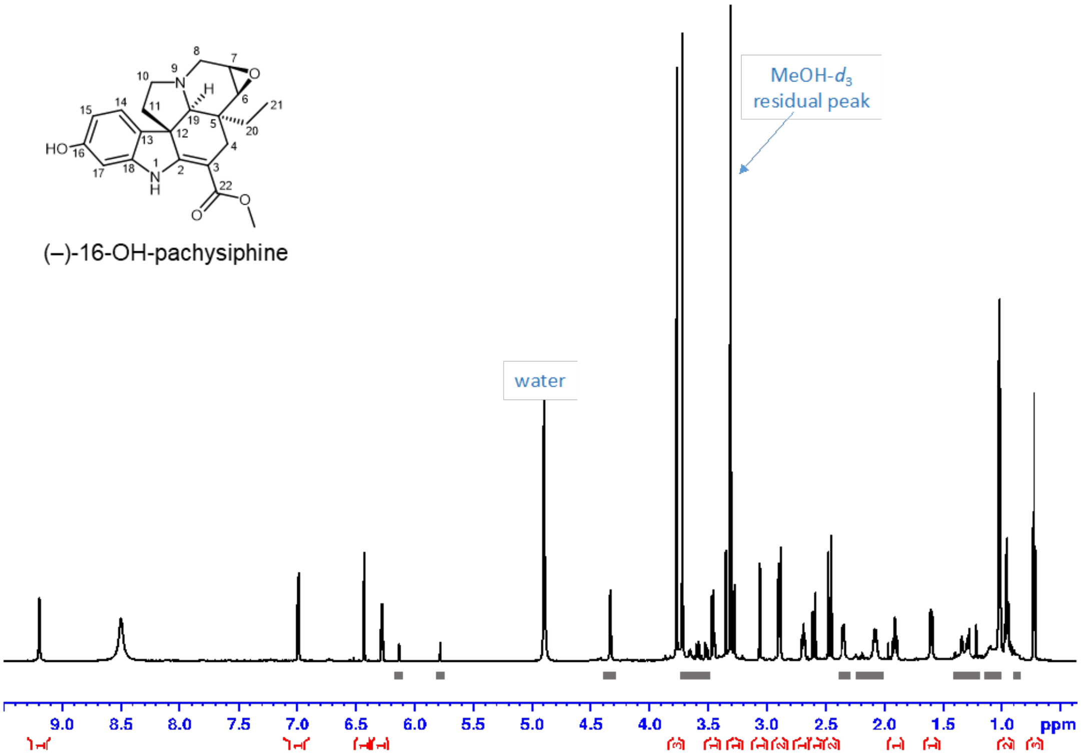


**$^1$H NMR with water suppression (zgpr)** full range in MeOH-$d_3$

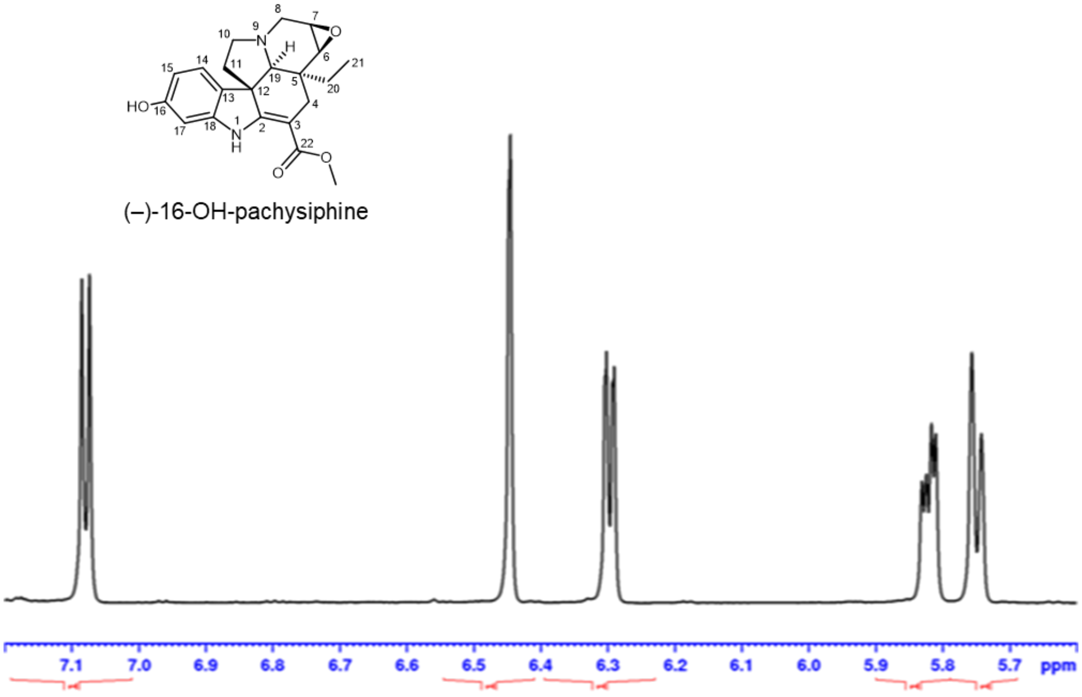


**$^1$H NMR with water suppression (zgpr)**
5.6-7.2 ppm in MeOH-$d_3$

**Fig. S12** Structure elucidation of (–)-16OH-pachysiphine by NMR. (Continued to next page).

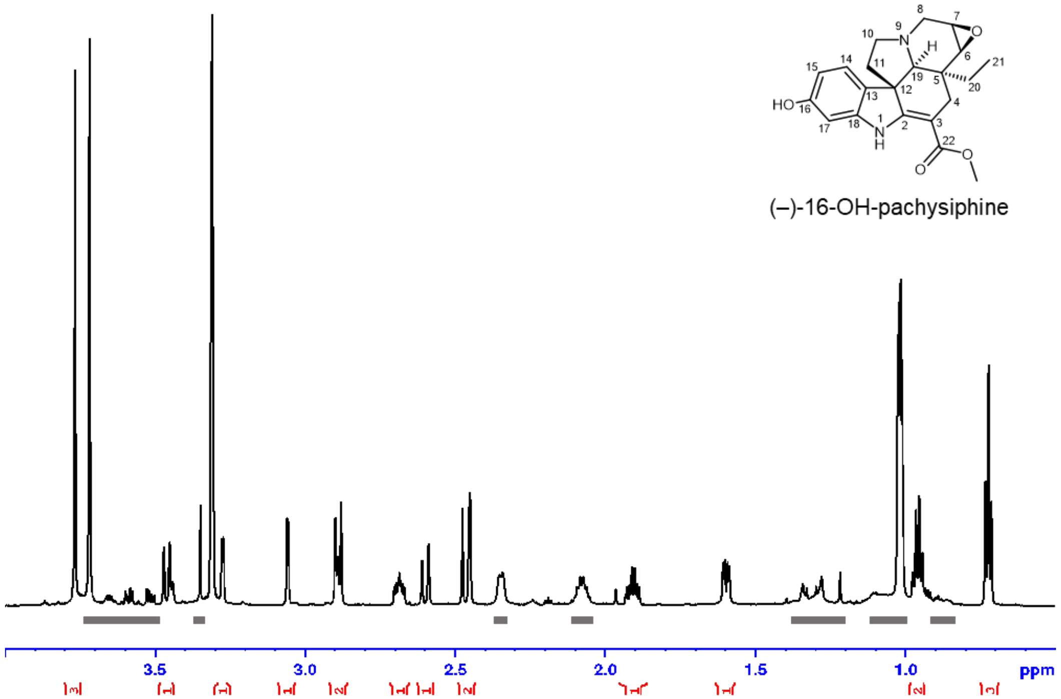


**$^1$H NMR with water suppression (zgpr)**
aliphatic range in MeOH-$d_3$. Shaded areas mark impurity

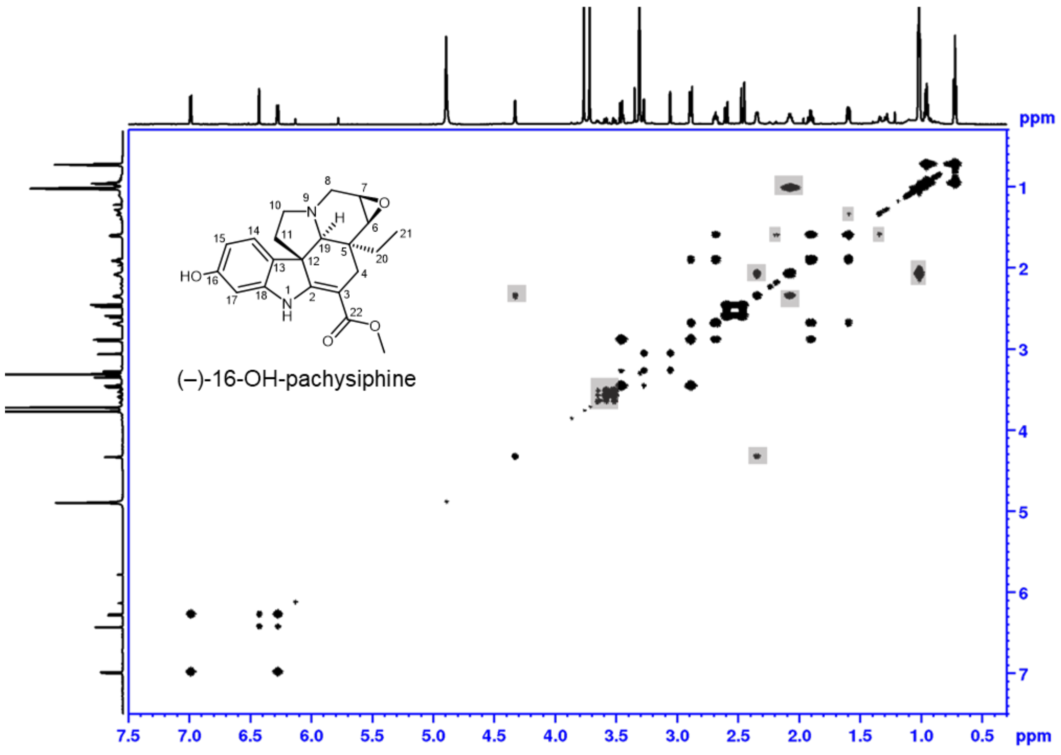


**$^1$H-$^1$H DQF COSY** with water suppression
full range in MeOH-$d_3$ .Shaded areas mark impurity

**Fig. S12** Structure elucidation of (–)-16OH-pachysiphine by NMR. (Continued to next page).

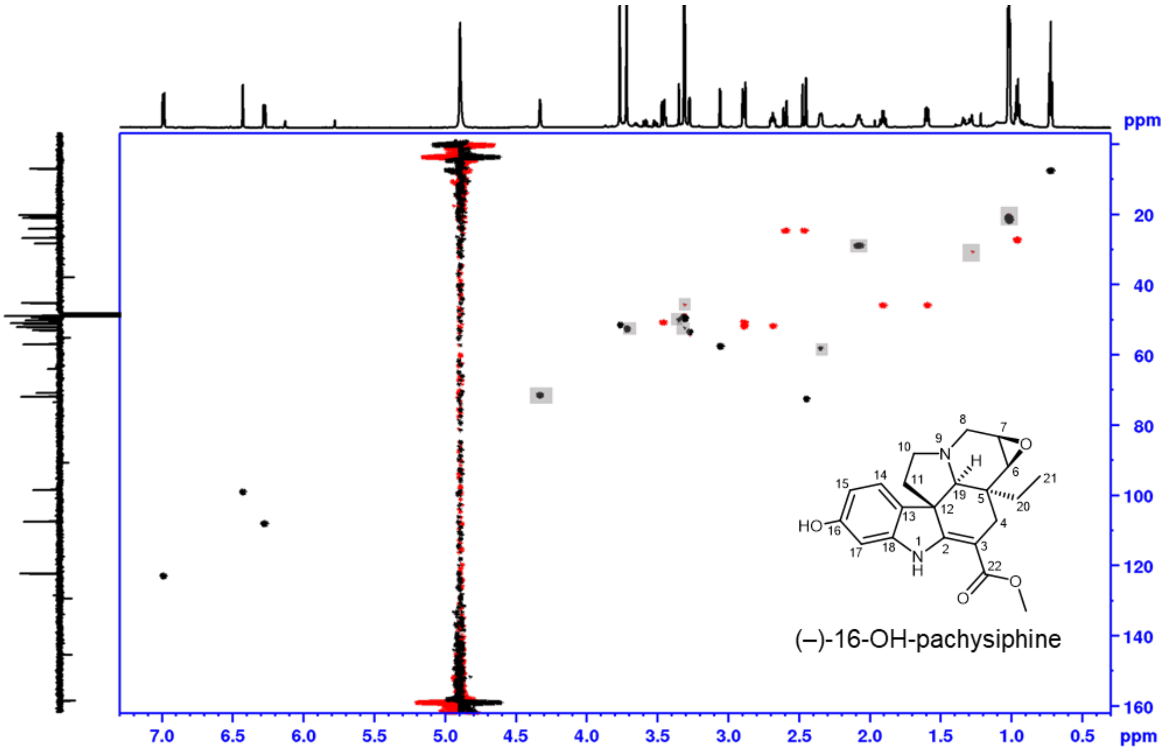


**phase sensitive HSQC**, full range in MeOH-$d_3$
red: $CH_2$, black: CH, $CH_3$. Shaded areas mark impurity

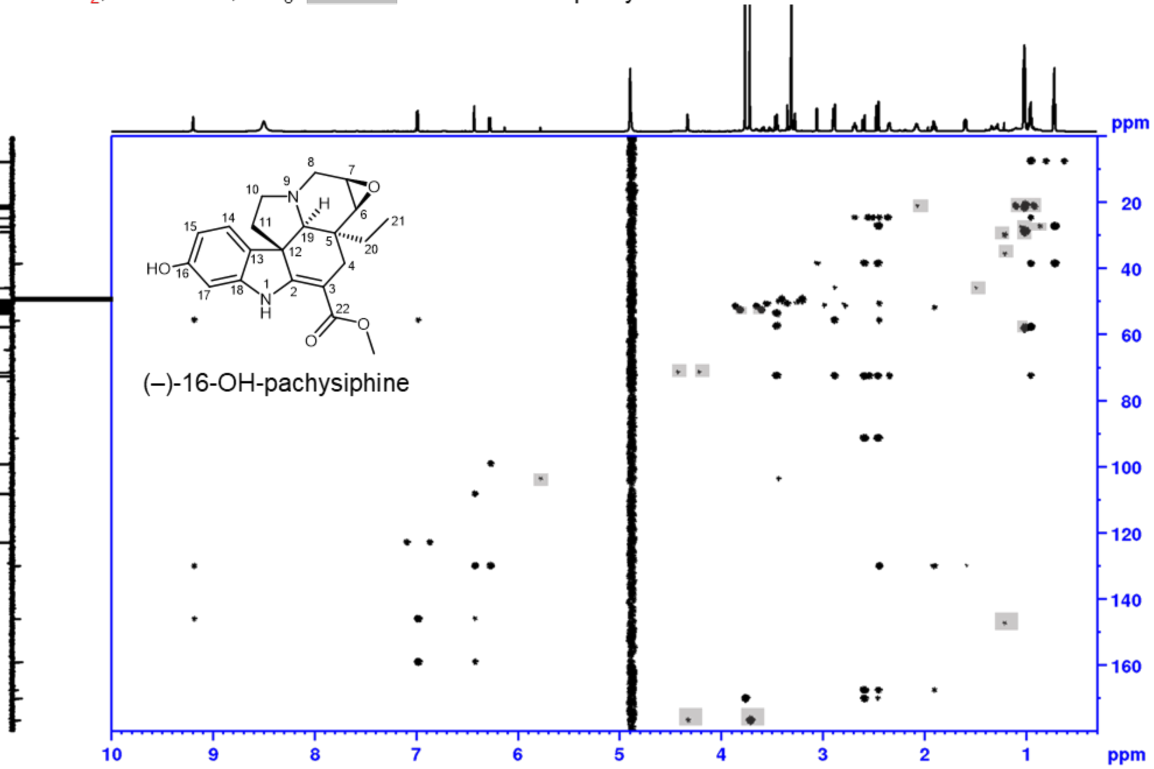


**HMBC**, full range in MeOH-$d_3$
Shaded areas mark impurity

**Fig. S12** Structure elucidation of (–)-16OH-pachysiphine by NMR. (Continued to next page).

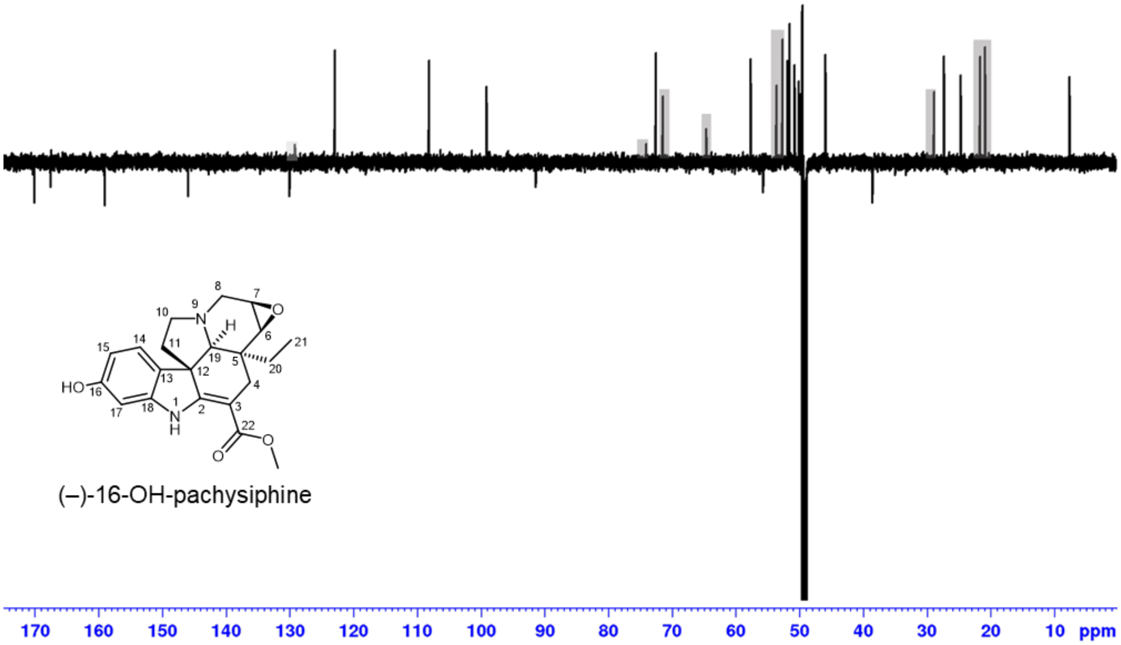


**DEPTQ**, full range in MeOH-$d_3$
Shaded areas mark impurity

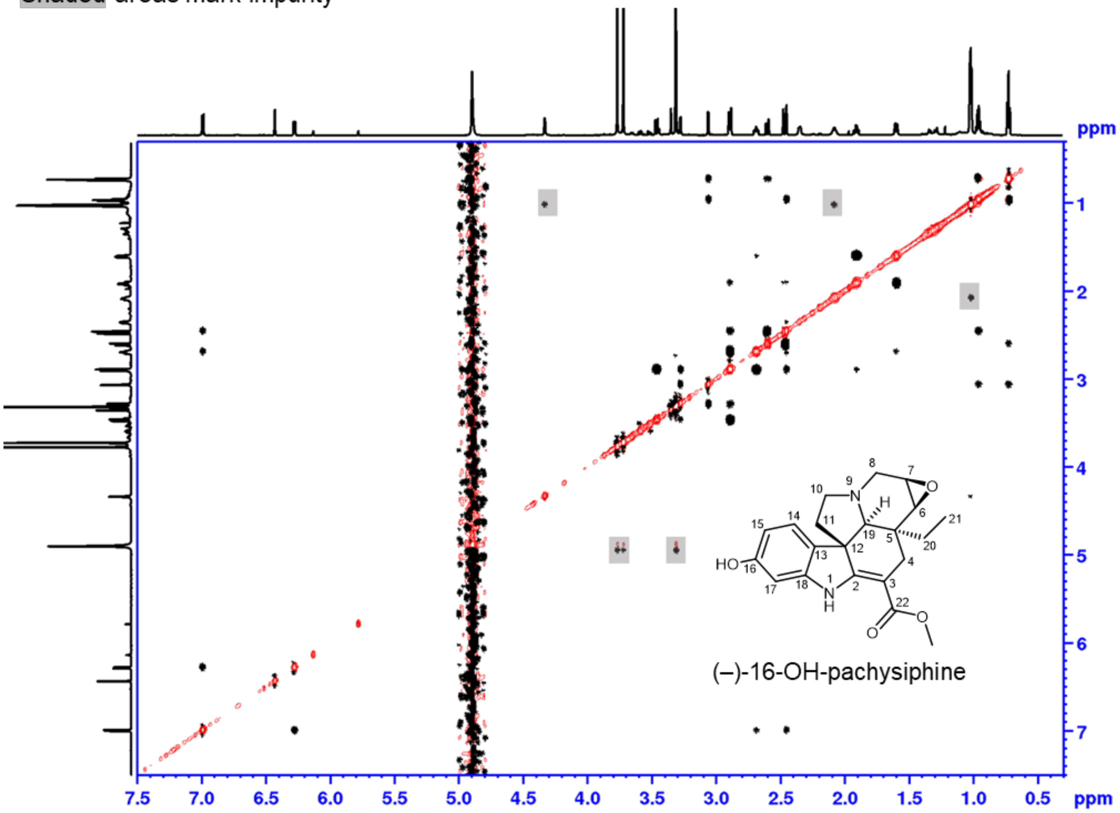


**$^{1}$H-$^{1}$H ROESY with water suppression**
full range in MeOH-$d_3$. Shaded areas mark impurity

**Fig. S12** Structure elucidation of (–)-16OH-pachysiphine by NMR. (Continued to next page).

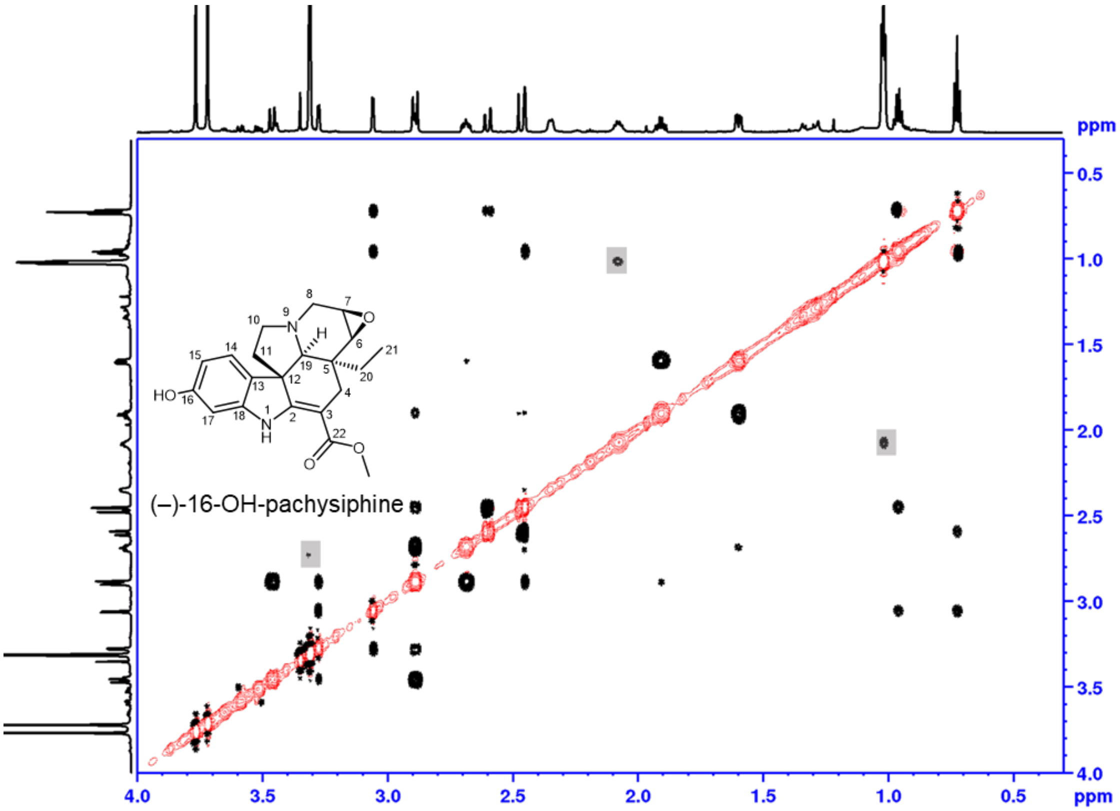


**$^1$H-$^1$H ROESY with water suppression.**
aliphatic range in MeOH-$d_3$. Shaded areas mark impurity

**Fig. S12** Structure elucidation of (–)-16OH-pachysiphine by NMR. (End).

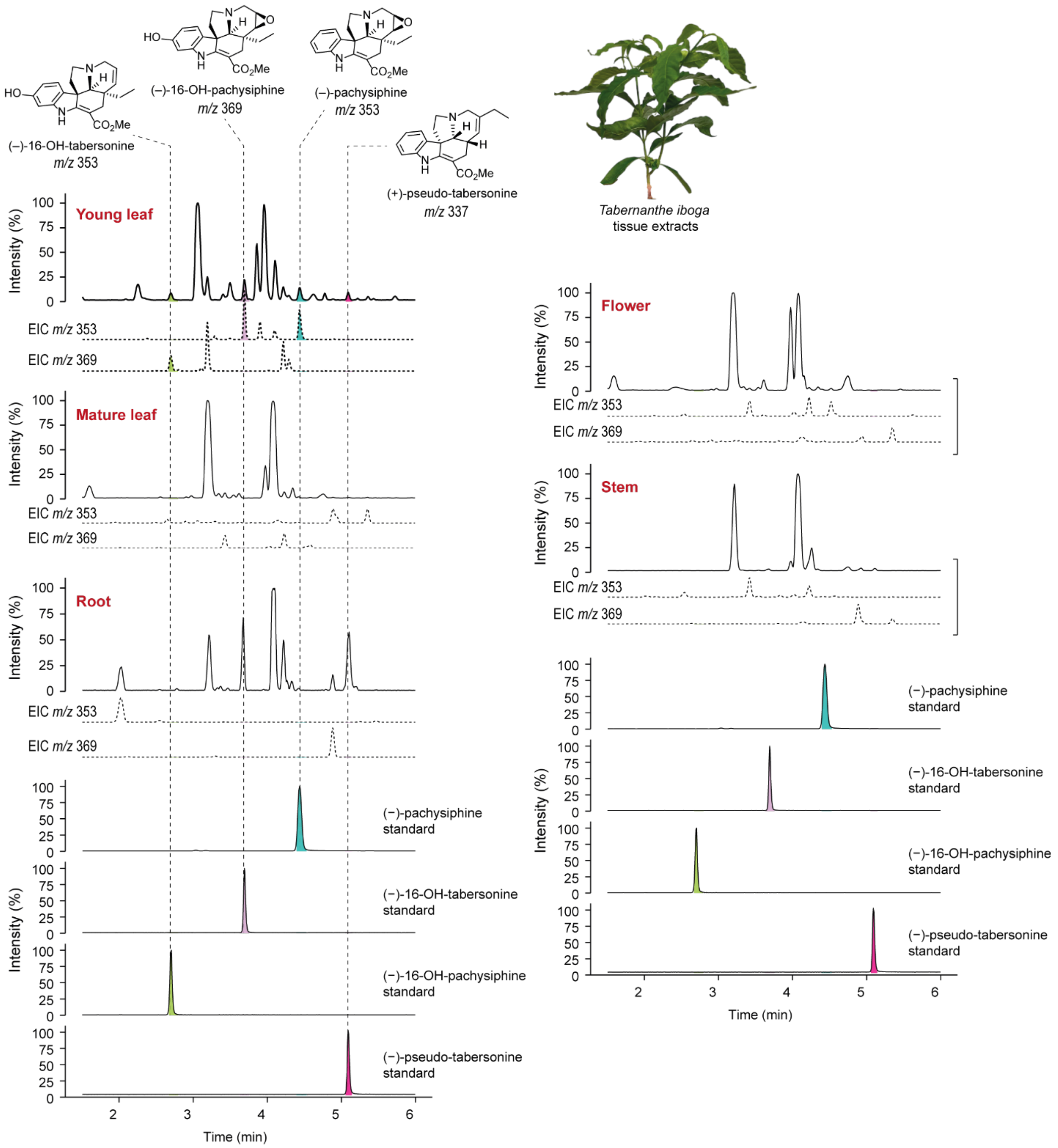


**Fig. S13** LC-MS metabolite profiles of *T. iboga* tissues. Untargeted LC-MS profiles of young leaves, mature leaves, stems, roots, and flowers of *T. iboga* methanolic extracts. The young leaf tissue shows an accumulation of pachysiphine, (–)-16-OH-tabersonine, and (–)-16-OH-pachysiphine as confirmed by authentic standards and $MS^2$ spectra. Total ion chromatograms are shown in solid lines, and extracted ion chromatograms are shown in dashed lines.

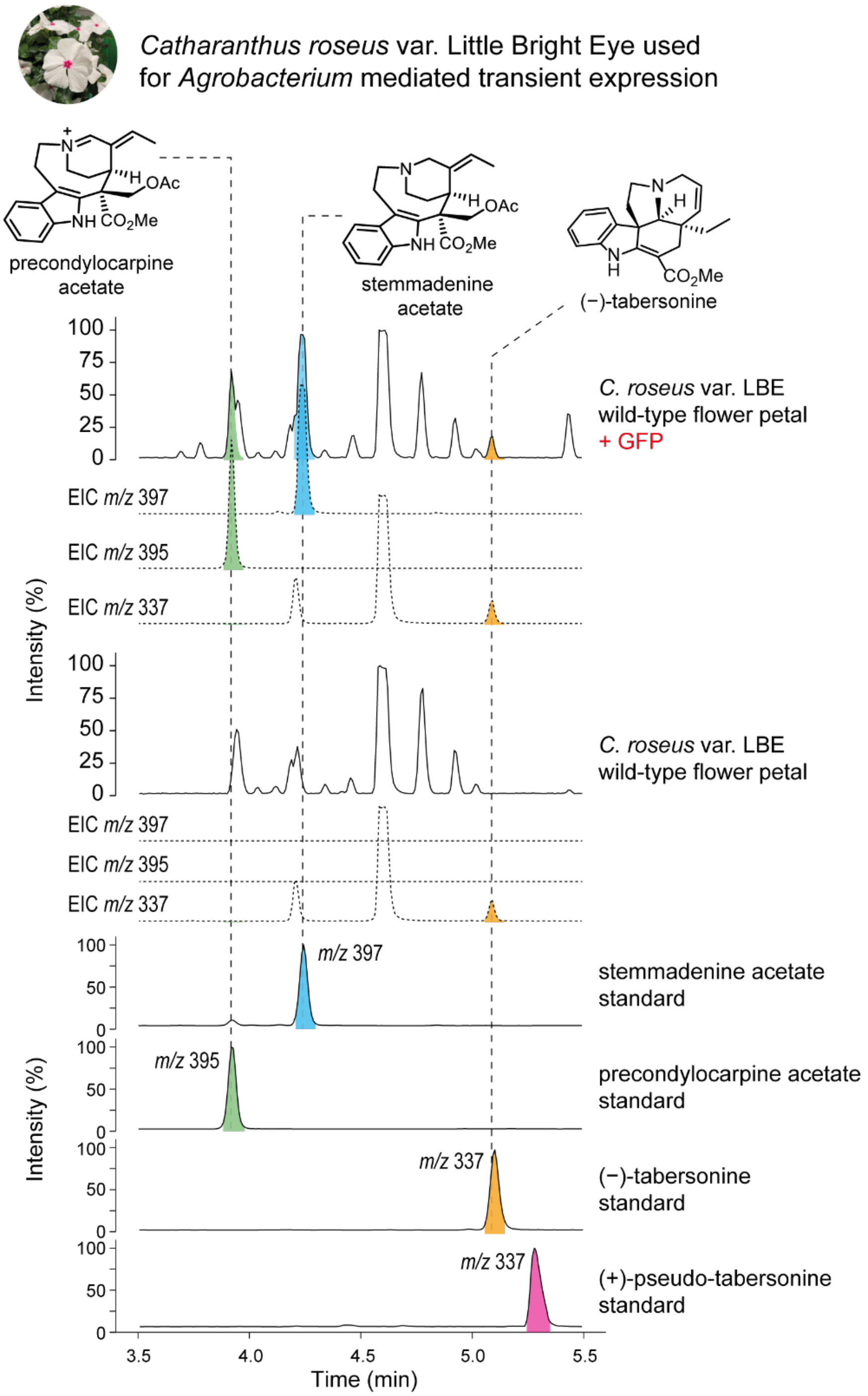


**Fig. S14** MIA precursors present in *C. roseus* var. LBE flower petals. Metabolic changes analyzed by LC-MS when *C. roseus* var. LBE flower petals are subjected to *Agrobacterium* infiltration (GFP control). Upon agroinfiltration, the flowers accumulate MIA alkaloids stemmadenine acetate, precondylocarpine acetate, and tabersonine as confirmed by authentic standards. These alkaloids are important precursors for MIA biosynthesis. LC-MS traces are presented as total ion chromatograms (solid lines) and extracted ion chromatograms are presented as dotted lines.

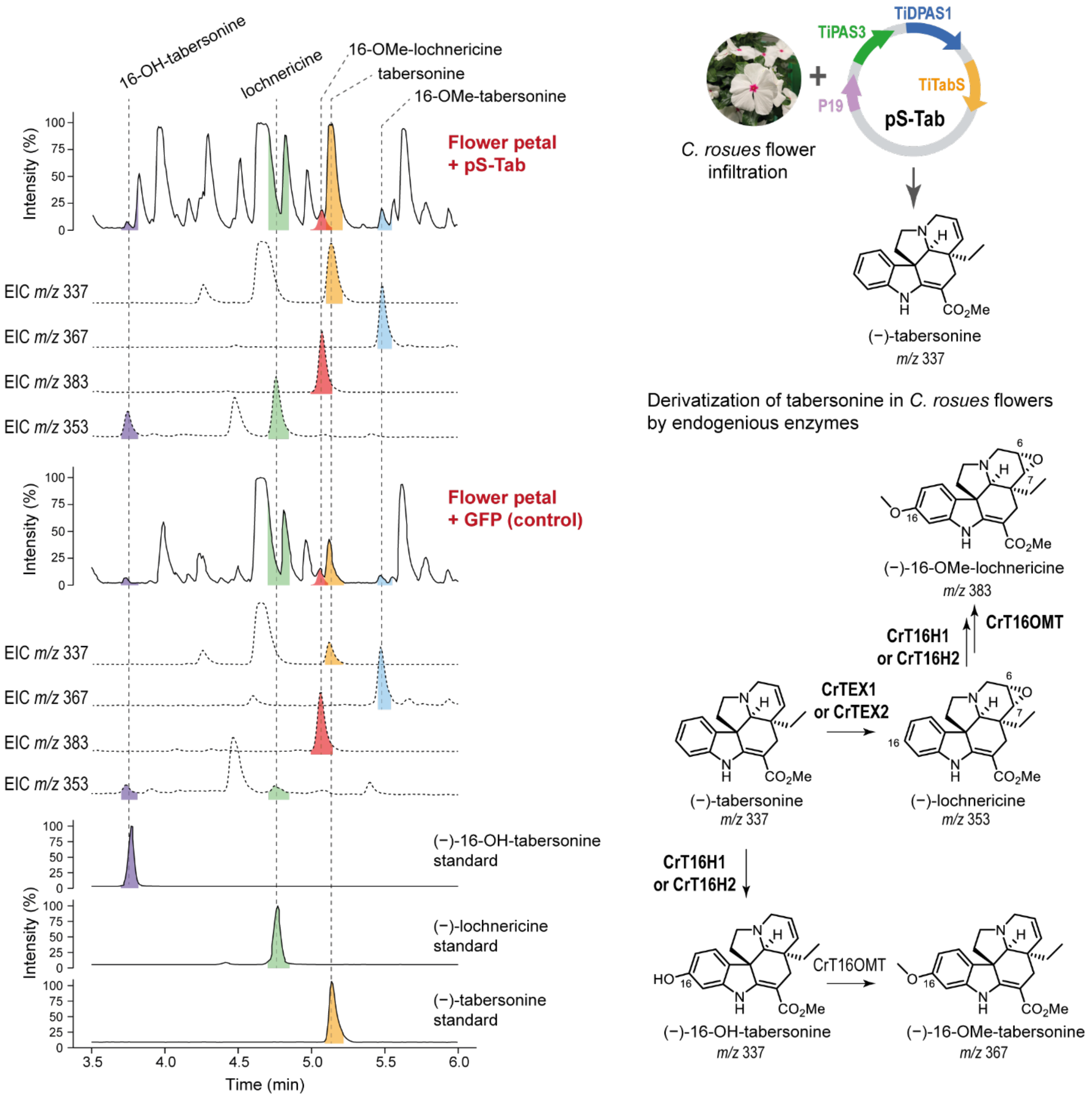


**Fig. S15** Derivatization of tabersonine by endogenous enzymes present in *C. roseus* flowers. LC-MS profiles of tabersonine derivatives observed when *C. roseus* flower petals are transfected with pS-Tab construct. In comparison to the GFP transfected control, the multigene construct pS-Tab substantially increases its product, tabersonine using endogenous substrate stemmadenine acetate. As confirmed by authentic standards, tabersonine is derivatized to lochnericine and 16-hydroxy-tabersonine by endogenous P450 enzymes CrTEX1/2 (Carqueijeiro *et al.*, 2018) and CrT16H1/2 (Besseau *et al.*, 2013) in the flowers. These compounds are further derivatized to 16-methoxy-tabersonine and 16-methoxy pachysiphine by CrT16OMTs in the flower (Sun *et al.*, 2018; Colinas *et al.*, 2021).

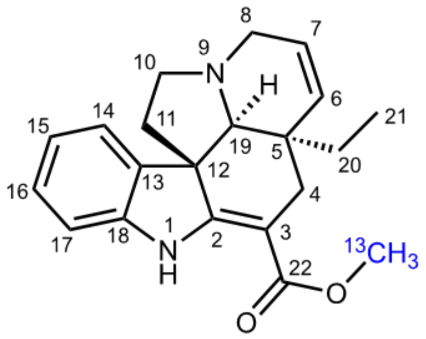


(–)-$^{13}$C-tabersonine
$^{1}$H-NMR, MeOH-$d_3$
500 MHz, 25 °C

| pos. | $\delta_H$ | mult. | $J_{HH}$ | $\delta_C$ |
|---|---|---|---|---|
| 1 | 9.29 | *bs* | - | - |
| 2 | - | - | - | 168.0 |
| 3 | - | - | - | 92.6 |
| 4a | 2.55 | *dd* | 15.2/1.7 | 29.9 |
| 4b | 2.44 | *d* | 15.2 | 29.9 |
| 5 | - | - | - | 42.5 |
| 6a | 5.71 | *ddd* | 10.0/1.8/1.8 | 134.2 |
| 7a | 5.81 | *ddd* | 10.0/4.8/1.5 | 126.8 |
| 8a | 3.22 | *ddd* | 15.9/1.8/1.5 | 51.6 |
| 8b | 3.45 | *ddd* | 15.9/4.8/1.8 | 51.6 |
| 10a | 2.76 | *ddd* | 11.2/8.5/4.7 | 51.8 |
| 10b | 3.03 | *dd* | 8.5/6.4 | 51.8 |
| 11a | 1.74 | *bdd* | 11.6/4.7 | 46.1 |
| 11b | 2.01 | *ddd* | 11.6/11.2/6.4 | 46.1 |
| 12 | - | - | - | 56.7 |
| 13 | - | - | - | 139.1 |
| 14 | 7.26 | *bd* | 7.5 | 122.4 |
| 15 | 6.86 | *ddd* | 7.5/7.5/0.7 | 121.8 |
| 16 | 7.13 | *ddd* | 7.9/7.5/1.1 | 129.0 |
| 17 | 6.94 | *bd* | 7.9 | 110.8 |
| 18 | - | - | - | 145.0 |
| 19a | 2.75 | *m*** | - | 71.4 |
| 20a | 0.87 | *dq* | 14.3/7.5 | 26.7 |
| 20b | 0.99 | *dq* | 14.3/7.5 | 26.7 |
| 21 | 0.63 | *t* | 7.5 | 7.8 |
| 22 | - | - | - | 170.1*** |
| OMe | 3.76 | *d** | 146.4* | 51.5 |

** overlapped signals J unresolved

*as $^{13}$C-H coupling

*** with $^{13}$C-$^{13}$C coupling (2.6 Hz)

**Fig. S16** Structure confirmation of (–)-[$^{13}$C]-tabersonine by NMR. (Continued to next page).

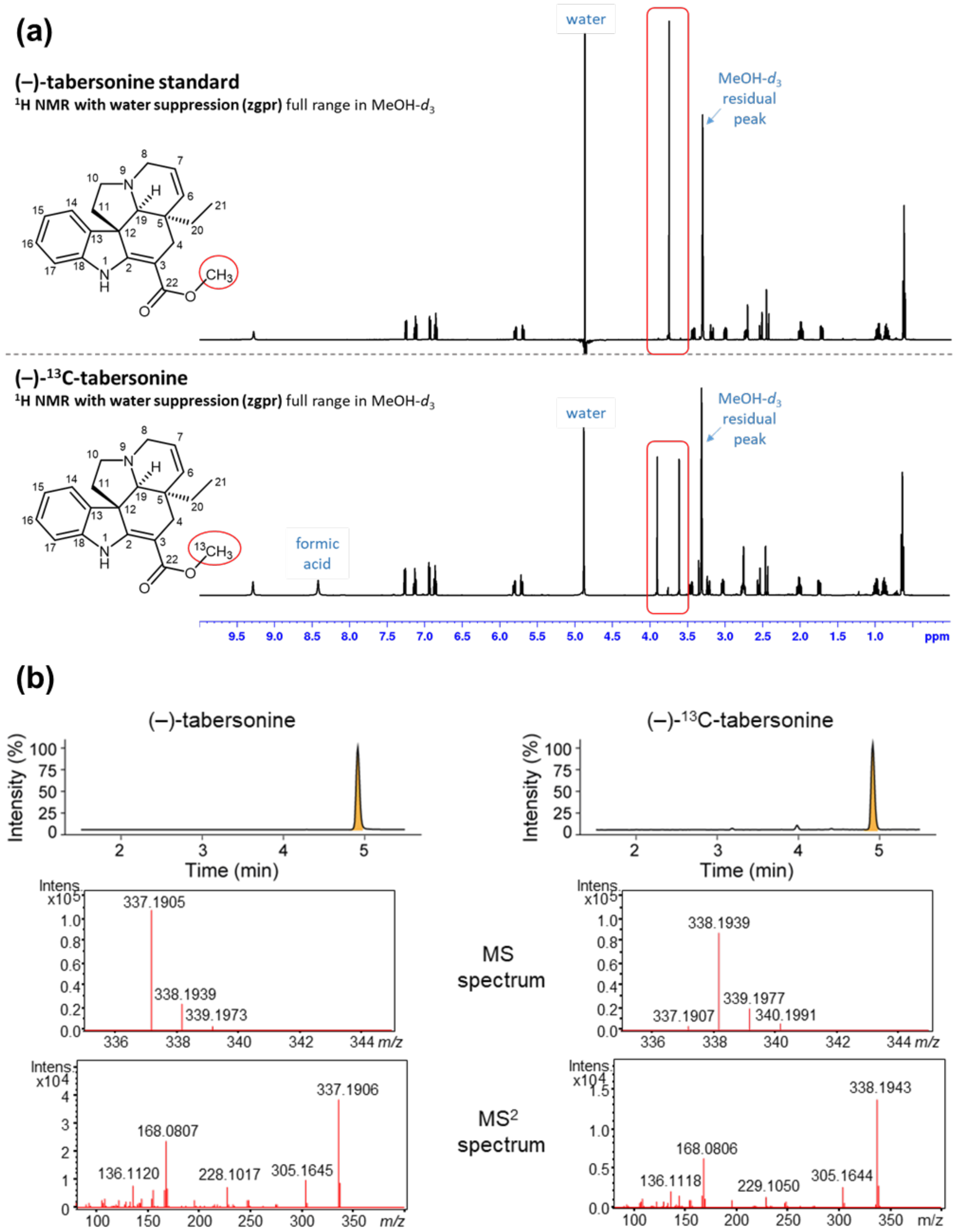


**Fig. S16** Structure confirmation of (–)-[$^{13}$C]-tabersonine. (a) NMR characterization of (–)-[$^{13}$C]-tabersonine. (b) LC-MS characterization of labeled, (–)-[13C]-tabersonine and un-labeled (–)-tabersonine. LC-MS traces are presented as total ion chromatograms.

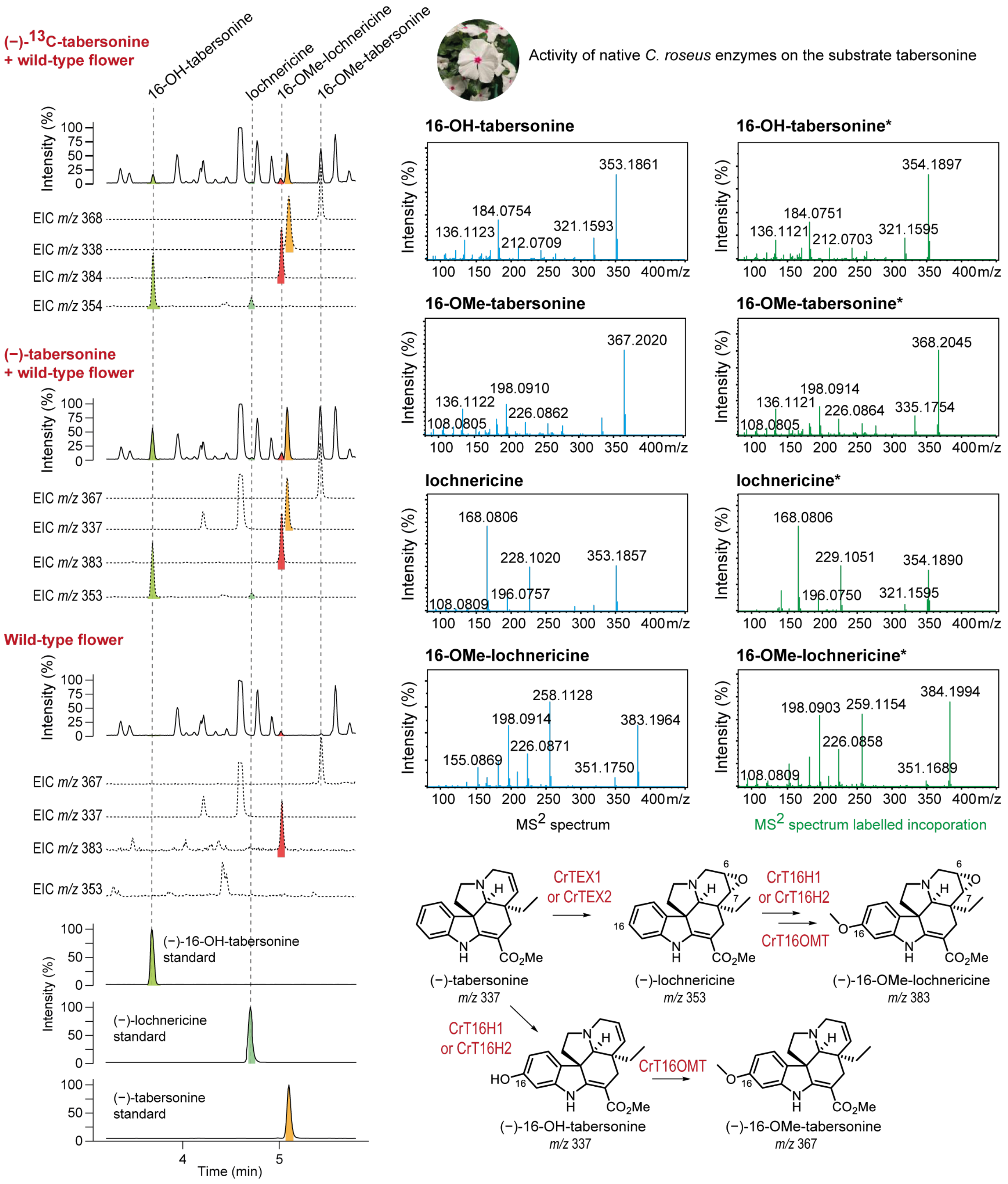


**Fig. S17** Incubation of *C. roseus* flowers with $^{13}$C-labelled tabersonine. LC-MS profiles of labeled tabersonine derivatives observed when *C. roseus* flowers were incubated with $^{13}$C-labeled tabersonine. $MS^2$ spectral comparison of standards and 13C-labeled tabersonine derivatives observed in the flowers reveals a biosynthetic network of tabersonine catalyzed by active endogenous enzymes present in the flowers. LC-MS traces with solid lines are presented as total ion chromatograms and dashed lines are extracted ion chromatograms.

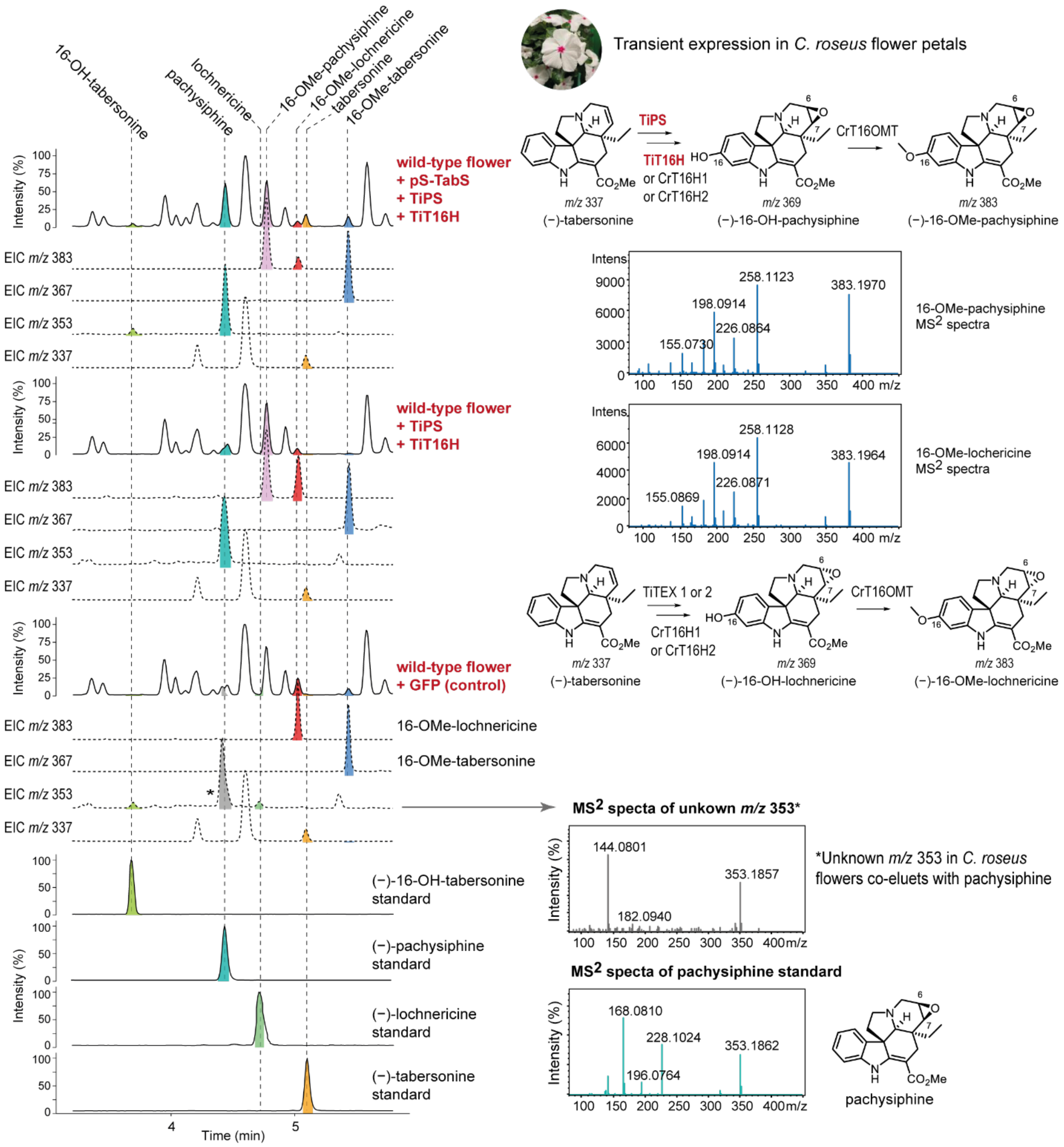


**Fig. S18** Overexpression of TiPS and TiT16H in *C. roseus* flowers. LC-MS analysis of *C. rosues* flowers simultaneously overexpressed with pS-Tab, TiPS and TiT16H indicate the formation of pachysiphine as confirmed by authentic standard and a derivative of pachysiphine with *m/z* 383 indicative of 16-methoxy-pachysiphine produced by endogenous flower enzymes. Both pachysiphine and 16-methoxy-pachysiphine are compounds not naturally present in *C. roseus.* Pachysiphine co-elutes with an unknown compound of *m/z* 353 (*) natively present in *C. roseus* flowers at low abundance but does not share $MS^2$ fragmentation pattern. LC-MS traces with solid lines are presented as total ion chromatograms and dashed lines are extracted ion chromatograms.

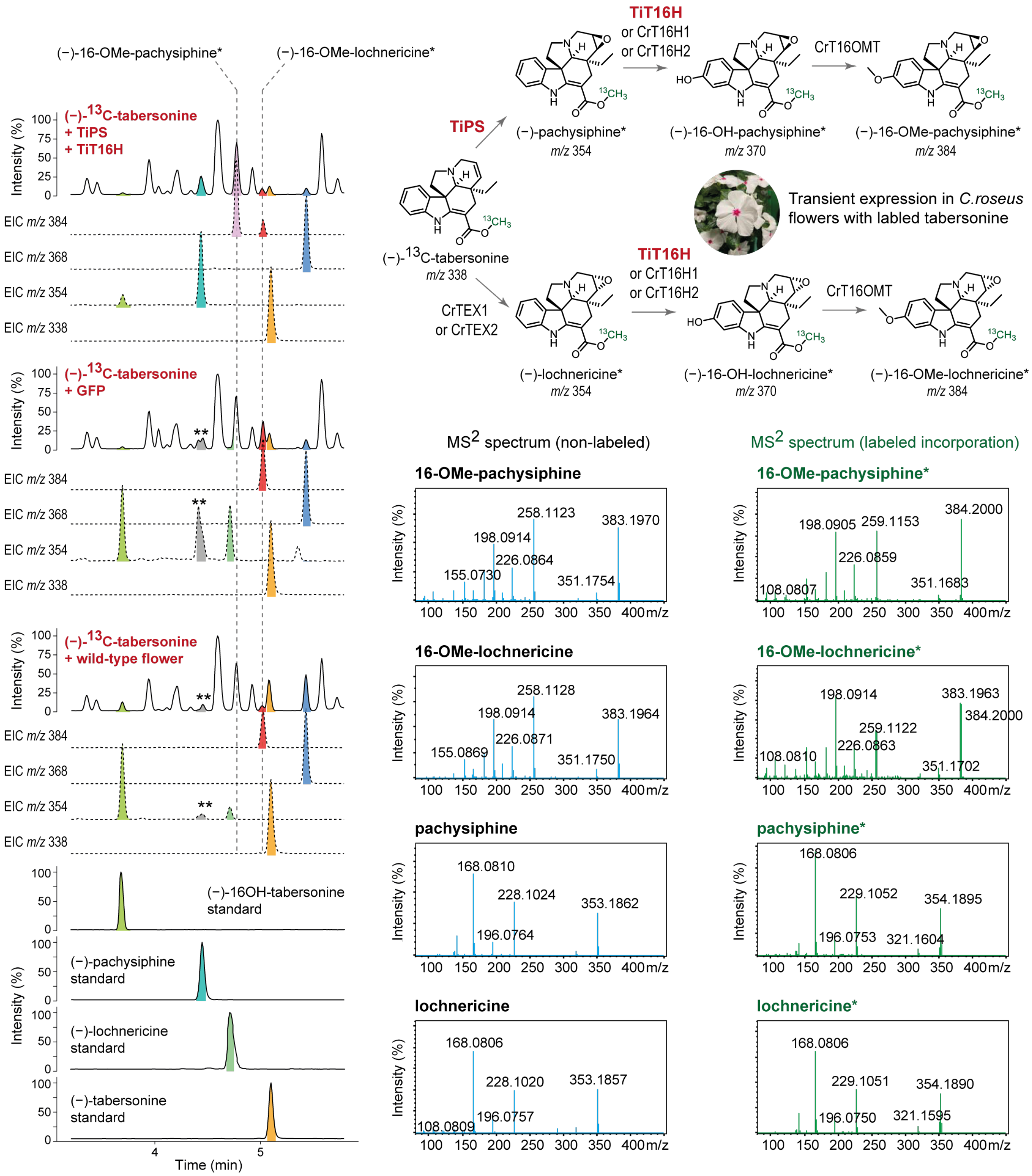


**Fig. S19** Overexpression of TiPS and TiT16H in *C. roseus* flowers with $^{13}$C-labeled tabersonine. LC-MS analysis and MS$^2$ spectral comparison confirm the formation of $^{13}$C-labeled pachysiphine catalyzed by TiPS and labeled pachysiphine derivative,16-metoxy-$^{13}$C-tabersonine catalyzed by TiT16H and flower endogenous enzymes. *MS$^2$ profiles of $^{13}$C-incorporated compounds. LC-MS traces with solid lines are presented as total ion chromatograms and dashed lines are extracted ion chromatograms. ** Unknown compound of *m/z* 353, which shares a similar retention time to pachysiphine (see Fig. S18).

| | TiPS | TiT16H | CrGO | CrT3O | CrT16H1 | CrT16H2 | CrT19H | CrTEX1 | CrTEX2 |
|---|---|---|---|---|---|---|---|---|---|
| TiPS | | 45.63 | 36.75 | 53.62 | 45.84 | 46.05 | 34.23 | 46.32 | 45.38 |
| TiT16H | 45.63 | | 32.94 | 42.35 | 56.34 | 56.89 | 35.4 | 53.42 | 53.1 |
| CrGO | 36.75 | 32.94 | | 36.4 | 33.53 | 32.5 | 33.33 | 36.72 | 36.03 |
| CrT3O | 53.62 | 42.35 | 36.4 | | 45.4 | 45.14 | 38.25 | 46.58 | 46.8 |
| CrT16H1 | 45.84 | 56.34 | 33.53 | 45.4 | | 80.85 | 35.27 | 57.62 | 56.65 |
| CrT16H2 | 46.05 | 56.89 | 32.5 | 45.14 | 80.85 | | 35.07 | 57.95 | 57.53 |
| CrT19H | 34.23 | 35.4 | 33.33 | 38.25 | 35.27 | 35.07 | | 36.17 | 36.21 |
| CrTEX1 | 46.32 | 53.42 | 36.72 | 46.58 | 57.62 | 57.95 | 36.17 | | 87.21 |
| CrTEX2 | 45.38 | 53.1 | 36.03 | 46.8 | 56.65 | 57.53 | 36.21 | 87.21 | |

**Fig. S20** Percentage of identity shared by TiPS and TiT16H with *C. roseus* P450s. Percentage identity of *T. iboga* pachysiphine synthase (TiPS) and tabersonine-16-hydroxylase (TiT16H) with known *C. roseus* P450s involved in aspidosperma (tabersonine) alkaloid hydroxylations.

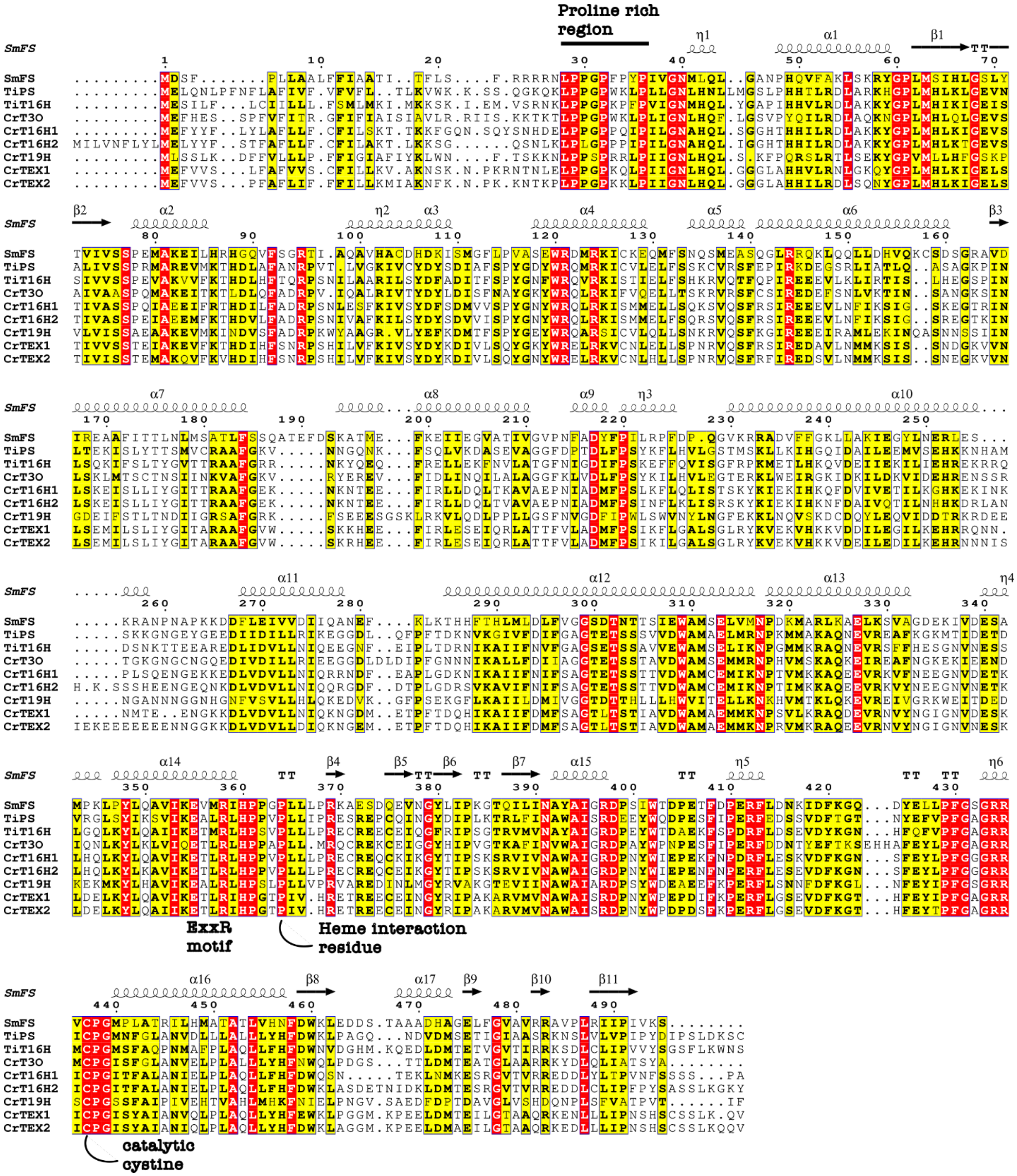


**Fig. S21** Sequence alignment of TiPS and TiT16H. Amino acid sequence alignment of *Tabernanthe iboga* pachysiphine synthase (TiPS) and tabersonine-16-hydroxylase (TiT16H) alongside known P450s from *Catharanthus roseus*. Structural features of the P450 enzyme are extracted from the crystal structure of *Salvia miltiorrhiza* Ferruginol synthase CYP76AH1 (PDB 7CB9).

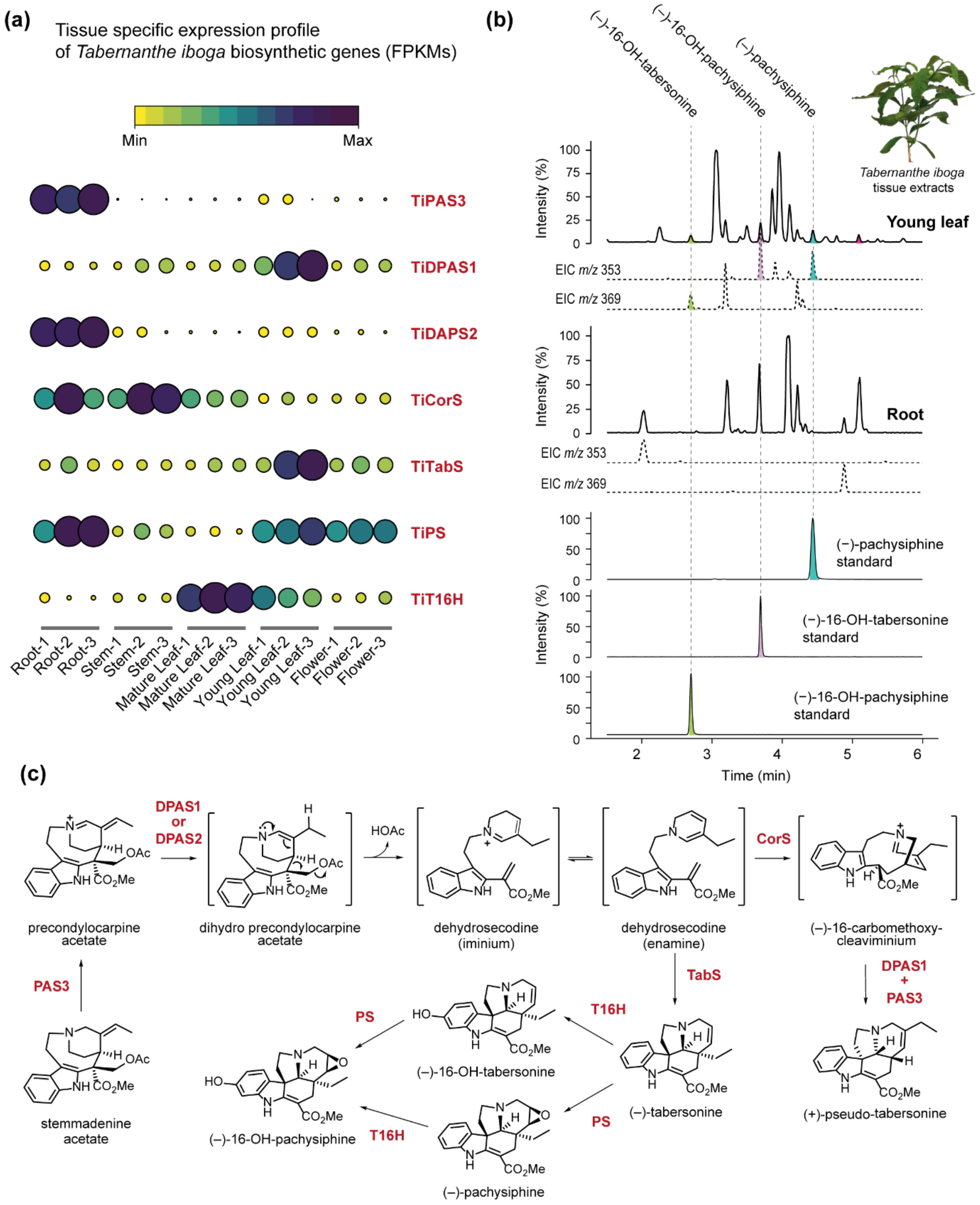


**Fig. S22** Transcriptomics and metabolomics related to (–)-16-OH-pachysiphine biosynthesis in *T. iboga.* (a) Tissue-specific transcriptomic profile of the genes involved in 16-OH-pachysiphine biosynthesis from stemmadenine acetate in *T. iboga*. (b) Representative metabolomic profile of *T. iboga* young leaf and root with authentic standards. (c) Biosynthesis of (–)-16OH-pachysiphine from stemmadenine acetate in *T. iboga*.

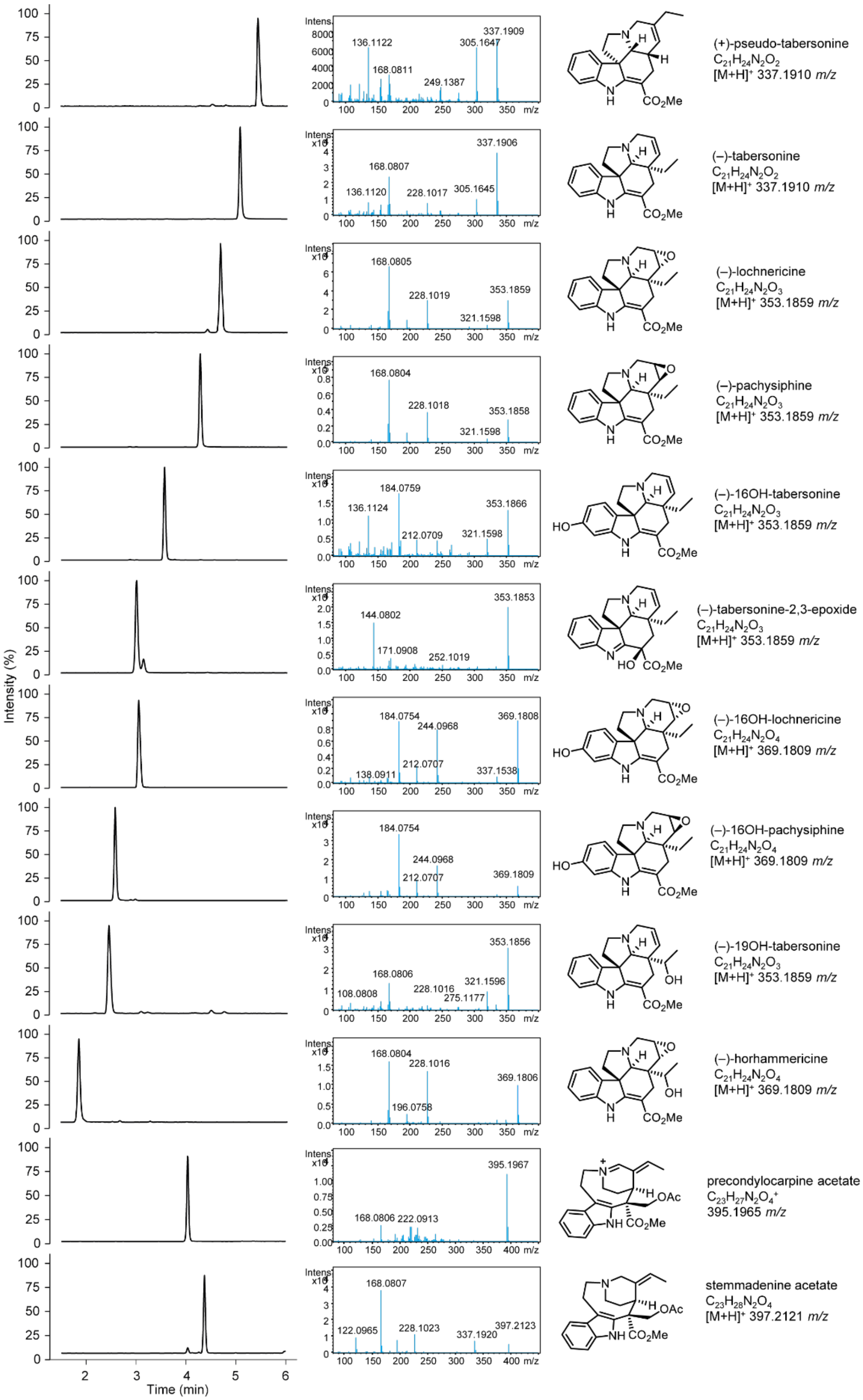


**Fig. S23** LC-MS profiles and $MS^2$ spectra of authentic standards based on $[M+H]^+$ parent ion.

**Table S1** Nucleotide sequences of genes transcriptional elements described in this study.

| **Gene Name (Genbank ID)** | **Nucleotide sequence** |
|---|---|
| P19<br>Goldenbraid 2.0 part ID<br>(GB0038) | ATGGAACGAGCTATACAAGGAAACGACGCTAGGGAACAAGCTAACAGTGAACGTTGGGA<br>TGGAGGATCAGGAGGTACCACTTCTCCCTTCAAACTTCCTGACGAAAGTCCGAGTTGGAC<br>TGAGTGGCGGCTACATAACGATGAGACTAATTCGAATCAAGATAATCCCCTTGGTTTCAA<br>GGAAAGCTGGGGTTTCGGGAAAGTTGTATTTAAGAGATATCTCAGATACGACAGGACGGA<br>AGCTTCACTGCACAGAGTCCTTGGATCTTGGACGGGAGATTCGGTTAACTATGCAGCATC<br>TCGATTTTTCGGTTTCGACCAGATCGGATGTACCTATAGTATTCGGTTTCGAGGAGTTAGT<br>ATCACCGTTTCTGGAGGCTCTCGAACTCTTCAGCATCTCTGTGAGATGGCAATTCGGTCT<br>AAGCAAGAACTGCTACAGCTTGCCCCAATCGAAGTGGAAAGTAATGTATCAAGAGGATGC<br>CCTGAAGGTACTGAAACCTTCGAAAAAGAAAGCGAGTGA |
| GFP | ATGGTGAGCAAGGGCGAGGAGCTGTTCACCGGGGTGGTGCCCATCCTGGTCGAGCTGG<br>ACGGCGACGTAAACGGCCACAAGTTCAGCGTGTCCGGCGAGGGCGAGGGCGACGCCAC<br>CTACGGCAAGCTGACCCTGAAGTTCATCTGCACCACCGGCAAGCTGCCCGTGCCCTGGC<br>CCACCCTCGTGACCACCTTCAGCTACGGCGTGCAGTGCTTCAGCCGCTACCCCGACCAC<br>ATGAAGCAGCACGACTTCTTCAAGTCCGCCATGCCCGAAGGCTACGTCCAGGAGCGCAC<br>CATCTTCTTCAAGGACGACGGCAACTACAAGACCCGCGCCGAGGTGAAGTTCGAGGGCG<br>ACACCCTGGTGAACCGCATCGAGCTGAAGGGCATCGACTTCAAGGAGGACGGCAACATC<br>CTGGGGCACAAGCTGGAGTACAACTACAACAGCCACAACGTCTATATCATGGCCGACAA<br>GCAGAAGAACGGCATCAAGGTGAACTTCAAGATCCGCCACAACATCGAGGACGGCAGCG<br>TGCAGCTCGCCGACCACTACCAGCAGAACACCCCCATCGGCGACGGCCCCGTGCTGCT<br>GCCCGACAACCACTACCTGAGCACCCAGTCCGCCCTGAGCAAAGACCCCAACGAGAAGC<br>GCGATCACATGGTCCTGCTGGAGTTCGTGACCGCCGCCGGGATCACTCACGGCATGGAC<br>GAGCTGTACAAGTGA |
| pUbq10<br>*Solanum lycopersicum*<br>Ubiquitin 10 promoter | GGAGGTCAACTACCCCAATTTAAATTTTATTTGATTAAGATATTTTTATGGACCTACTTTATA<br>ATTAAAAATATTTTCTATTTGAAAAGGAAGGACAAAAATCATACAATTTTGGTCCAACTACT<br>CCTCTCTTTTTTTTTTTGGCTTTATAAAAAAGGAAAGTGATTAGTAATAAATAATTAAATAAT<br>GAAAAAAGGAGGAAATAAAATTTTCGAATTAAAATGTAAAAGAGAAAAAGGAGAGGGAGT<br>AATCATTGTTTAACTTTATCTAAAGTACCCCAATTCGATTTTACATGTATATCAAATTATACA<br>AATATTTTATTAAAATATAGATATTGAATAATTTTATTATTCTTGAACATGTAAATAAAAATTA<br>TCTATTATTTCAATTTTTATATAAACTATTATTTGAAATCTCAATTATGATTTTTTAATATCAC<br>TTTCTATCCATGATAATTTCAGCTTAAAAAGTTTTGTCAATAATTACATTAATTTTGTTGATG<br>AGGATGACAAGATTTCGGTCATCAATTACATATACACAAATTGAAATAGTAAGCAACTTGA<br>TTTTTTTTCTCATAATGATAATGACAAAGACACGAAAAGACAATTCAATATTCACATTGATTT<br>ATTTTTATATGATAATAATTACAATAATAATATTCTTATAAAGAAAGAGATCAATTTTGACTG<br>ATCCAAAAATTTATTTATTTTTACTATACCAACGTCACTAATTATATCTAATAATGTAAAACA<br>ATTCAATCTTACTTAAATATTAATTTGAAATAAACTATTTTTATAACGAAATTACTAAATTTAT<br>CCAATAACAAAAAGGTCTTAAGAAGACATAAATTCTTTTTTTGTAATGCTCAAATAAATTTG<br>AGTAAAAAAGAATGAAATTGAGTGATTTTTTTTTAATCATAAGAAAATAAATAATTAATTTCA<br>ATATAATAAAACAGTAATATAATTTCATAAATGGAATTCAATACTTACCTCTTAGATATAAAA<br>AATAAATATAAAAATAAAGTGTTTCTAATAAACCCGCAATTTAAATAAAATATTTAATATTTT<br>CAATCAAATTTAAATAATTATATTAAAATATCGTAGAAAAAGAGCAATATATAATACAAGAAA<br>GAAGATTTAAGTACAATTATCAACTATTATTATACTCTAATTTTGTTATATTTAATTTCTTACG<br>GTTAAGGTCATGTTCACGATAAACTCAAAATACGCTGTATGAGGACATATTTTAAATTTTAA<br>CCAATAATAAAACTAAGTTATTTTTAGTATATTTTTTTGTTTAACGTGACTTAATTTTTCTTTT<br>CTAGAGGAGCGTGTAAGTGTCAACCTCATTCTCCTAATTTTCCCAACCACATAAAAAAAAA<br>ATAAAGGTAGCTTTTGCGTGTTGATTTGGTACACTACACGTCATTATTACACGTGTTTTCGT<br>ATGATTGGTTAATCCATGAGGCGGTTTCCTCTAGAGTCGGCCATACCATCTATAAAATAAA<br>GCTTTCTGCAGCTCATTTTTTCATCTTCTATCTGATTTCTATTATAATTTCTCTGAATTGCCT<br>TCAAATTTCTCTTTCAAGGTTAGAATTTTTCTCTATTTTTTGGTTTTTGTTTGTTTAGATTCT<br>GAGTTTAGTTAATCAGGTGCTGTTAAAGCCCTAAATTTTGAGTTTTTTTCGGTTGTTTTGAT<br>GGAAAATACCTAACAATTGAGTTTTTTCATGTTGTTTTGTCGGAGAATGCCTACAATTGGA<br>GTTCCTTTCGTTGTTTTGATGAGAAAGCCCCTAATTTGAGTGTTTTTCCGTCGATTTGATTT<br>TAAAGGTTTATATTCGAGTTTTTTTCGTCGGTTTAATGAGAAGGCCTAAAATAGGAGTTTTT<br>CTGGTTGATTTGACTAAAAAAGCCATGGAATTTTGTGTTTTTGATGTCGCTTTGGTTCTCAA<br>GGCCTAAGATCTGAGTTTCTCCGGTTGTTTTGATGAAAAAGCCCTAAAATTGGAGTTTTTA<br>TCTTGTGTTTTAGGTTGTTTTAATCCTTATAATTTGAGTTTTTTCGTTGTTCTGATTGTTGTT<br>TTTATGAATTTTGCAG |
| tUbq10<br>*Solanum* | GCTTGTTGTGGTTGTCTGGTTGCGTCTGTTGCCCGTTGTCTGTTGCCCATTGTGGTGGTT<br>GTGTTTGTATGATGGTCGTTAAGGATCATCAATGTGTTTTCGCTTTTTGTTCCATTCTGTTT |

| | |
|---|---|
| *lycopersicum* Ubiquitin 10 terminator | CTCATTTGTGAATAATAATGGTATCTTTATGAATATGCAGTTTGTGGTTTCTTTTCTGATTG<br>CAGTTCTGAGCATTTTGTTTTTGCTTCCGTTTACTATACCACTTACAGTTTGCACTAATTTA<br>GTTGATATGCGAGCCATCTGATGTTTGATGATTCAAATGGCGTTTATGTAACTCGTACCCG<br>AGTGGATGGAGAAGAGCTCCATTGCCGGTTTGTTTCATGGGTGGCGGAGGGCAACTCCT<br>GGGAAGGAACAAAAGAAAAACCGTGATACGAGTTCATGGGTGAGAGCTCCAGCTTGATC<br>CCTTCTCTGTCGATCAAATTTGAATTTTTGGATCACGGCAGGCTCACAAGATAATCCAAAG<br>TAAAACATAATGAATAGTACTTCTCAATGATCACTTATTTTTAGCAAATCAGCAATTGTGCA<br>TGTCAAATGATTTCGGTGTAAGAGAAAGAGTTGATGAATCAAAATATCTGTAGCTGGATCA<br>AGAATCTGAGGCAGTTGTATGTATCAATGATCTTTCCGCTACAATGATGTTAGCTATCCGA<br>GTCAAATTGTTGTAGAATTGCATACTTCGGCATCACATTCTGGATGACATAATAAATAGGA<br>AGTCTTCAGATCCCTAAAAAATTGAGAGCTAATAACATTAGTCCTAGATGTAACTGGGTGA<br>CAACCAAGAAAGAGACATGCAAATACTACTTTTGTTTGAAGGAGCATCCCTGGTTTGACAT<br>ATTTTTTCTGAATATCAAACTTTGAAACTCTACCTAGTCTAATGTCTAACGACAGATCTTAC<br>TGGTTTAACTGCAGTGATATCTACTATCTTTTGGAATGTTTTCTCCTTCAGTTATACATCAA<br>GTTCCAAGATGCAGGTGTGCTTGATTGATGTACATGGCTGTGAGAAGTGCATCCTGATGT<br>TCAGATGATGGTTCATTCTAATGTCTTTTCCTTCAATCAGTTTTCTCAGTCTGACTTAGCTT<br>GTTTCATCTGCATGTTTGAATGTTCGTTTACTCATAGTAATTGCATTTTTGTAGCAGAACAT<br>ATCATTGGTCATGGTTTCAACTGTGCGCGAGTCTTATGCTTATTCAAACTAGGAAAGCCTC<br>CGTCTAGAGGGTACACGAGTTGTTGCTCTGTGTGCGTCAGTCCATAGTATTAATCTTGCTA<br>GTTGTAGTATATTGTTTATGTGGACTCGGAATTCATCATATGCTCCTTCTTTGCATCAAGTA<br>AGGCAAGGTAATGTATAGAAGCTTTTTAACTCTTTCATGGAAGCTGGCCTTTGCCAGCATA<br>CCATCCAGAAGATATCAACCCTGCATCTTGGCTGCCG |
| TiPAS3 (BBL-like) (MK840852.1) | ATGATAAAAAAAGTCCCAATAGTTCTTTCAATTTTCTGCTTTCTTCTTCTACTCTCATCATCC<br>CATGGCTCAATTCCTGAAGCTTTTCTCAATTGTATTTCCAATAAATTTTCATTAGATGTATC<br>CATTTTAAACATTCTTCATGTTCCCAGCAATTCTTCCTATGATTCTGTTCTCAAATCTACTAT<br>CCAAAATCCAAGATTCCTCAAATCACCCAAGCCCTTAGCTATAATCACCCCAGTACTTCAC<br>TCCCATGTCCAATCTGCTGTTATCTGTACCAAACAAGCCGGTTTACAAATTAGAATCCGAA<br>GCGGAGGAGCTGATTACGAGGGCTTATCCTATCGTTCTGAGGTTCCCTTTATTCTGCTAG<br>ATCTCCAGAATCTTCGATCAATTTCCGTTGATATTGAAGACAACAGCGCTTGGGTCGAATC<br>AGGAGCAACAATTGGTGAATTCTATCATGAGATAGCTCAGAACAGCCCTGTTCATGCGTTT<br>CCAGCTGGGGTCTCTTCCTCTGTTGGAATTGGCGGCCATTTGAGTAGCGGCGGTTTTGGT<br>ACATTGCTTCGGAAATATGGATTAGCAGCCGATAATATAATCGATGCAAAAATTGTTGATG<br>CCAGAGGCAGAATTCTTGATAGGGAATCAATGGGAGAAGATCTATTTTGGGCTATTAGAG<br>GAGGAGGAGGAGCTAGTTTTGGTGTTATAGTTTCTTGGAAGGTTAAACTTGTAAAAGTCC<br>CTCCGATGGTAACTGTTTTCATCTTGTCCAAGACTTATGAAGAAGGAGGTTTAGATCTTCT<br>ACACAAATGGCAATATATAGAACACAAACTCCCTGAAGATTTATTCCTTGCTGTAAGCATC<br>ATGGATGATTCATCTAGTGGAAATAAAACACTTATGGCAGGTTTTATGTCTCTGTTTCTTG<br>GAAAAACAGAGGACCTTCTGAAAGTAATGGCGGAAAATTTCCCACAACTTGGATTGAAAA<br>AGGAAGATTGCTTAGAAATGAATTGGATTGATGCAGCAATGTATTTTTCAGGACACCCAAT<br>TGGAGAATCCCGATCTGTGCTTAAAAACCGAGAATCTCATCTTCCAAAGACATGCGTTTCG<br>ATCAAATCAGACTTTATTCAAGAACCACAATCCATGGATGCATTGGAAAAGTTATGGAAGT<br>TTTGTAGGGAAGAAGAAAATAGTCCCATAATACTGATGCTTCCACTGGGGGGAATGATGA<br>GTAAAATATCAGAATCAGAAATCCCATTTCCTTACAGAAAAGATGTGATTTACAGTATGATA<br>TACGAAATAGTTTGGAATTGTGAAGACGATGAATCATCGGAAGAATATATCGATGGATTGG<br>GAAGGCTTGAGGAATTAATGACTCCATATGTGAAACAACCAAGAGGTTCTTGGTTCAGCA<br>CCAGAAACCTTTATACCGGTAAAAATAAAGGTCCAGGAACAACTTATTCCAAAGCTAAAGA<br>ATGGGGATTTCGGTATTTTAATAATAATTTCAAAAAGTTGGCCCTTATCAAAGGACAAGTT<br>GATCCAGAAAACTTCTTCTACTATGAACAAAGCATTCCCCCTCTCCATTTACAAGTCGAAC<br>TTTAA |
| TiDPAS1 (ADH) (MK840855.1) | ATGGCTGTAAAATCACCTGAAGCAGAGCACCCAGTGAAGGCTGTAAAATCACCTGAAGAA<br>GAGCACCCAGTGAAGGCATACGGATGGGCTATCAAAGACAGAACATCTGGCATTCTTTCC<br>CCCTTCAAGTTTTCCAGAAGGGCAACAGGAGATGAAGACGTTCGAATAAAGATCCTCTGT<br>TGCGGAGTTTGTCACACGGATCTCACGTCTACCAAGAATGAATACGAGTTTCTTTCATATC<br>CGCTAGTGCCTGGGTTAGAGACTGTCGGAATAGCGACAGAGGTCGGAAGCAAAGTCACA<br>AAAGTAAAAGTTGGTGAAAAAGTAGCAGTGGCAGCCTATTTGGGCACTTGTGGAAAATGC<br>CACAATTGTCTAAATGACCAAGAGAATTACTGTCCCGAAGTGATCATTAGCTACGGCACAC<br>CATATCACGACGGAACAATCAACTACGGAGGCTTCTCGAATGAGACAGTCGTAAATGAGC<br>GCTTCGTTCTTCATTTTCCTGAAAAGCTTTCACTTTCTGGTGTTGCACCGCTAGTCAGCGC<br>AGGAAGCACCGCTTACAGTGCAATAAGAAATCAAGGCCTTGACAAACCCGGTATCCACTT<br>GGGAGTCGTCGGCCTTGGTGGACTTGGTCATCTGGCTGTGAAGTTTGCAAAGGCTTTTG<br>GTGTCAAGGTGACAGTGATTAGTTCCACTCCCAGCAAGAAGGATGAAGCCATCAAGAGCC<br>TTGGTGCCGATGCGTTCTTGTTCAGTCGTGATGATGAACAAATGAAGGCCGCTATTGGAA |

| | |
|---|---|
| | CTTTCGATGCAATCATAGATACTATTGCAGTCGCTCATCCTCTTGCGCCATTACTTGATCT<br>ACTAAGGAGTCATGGAAAAATTATTTTGGTCGGGGCACCGACCACCCCACTTGAGGTGCC<br>AGTTATTCCTTTAGTAGCAGGAGGGAAATCAATTACTGGATGCGTAGTTGGAAATTTGAAG<br>CAAACTCAAGAAATGCTTGAATTTGCTGCAGAACACAACATCACTGCAAACGTTGAGGTTA<br>TTTCAATGGATTACATAAACACTGCAATGGAACGTTTAGAAAAAGGTGATGTTAGATATAG<br>ATTTGTAATTGATATTGGAAACACACTAACTCCACCGGAATAG |
| TiTabS<br>(Hydrolase)<br>(MK840853.1) | ATGGCTTCTTCAACTGAAAGCTCTGATGAGATTATTTTTGATCTTCCTCCATACATTAGAGT<br>CTTTAAGGATGGAAGAGTAGAGAGACTCCACTCCTCACCATATGTTCCACCATCACTAGAT<br>GATCCCGCAACCGGCGTATCCTGGAAAGACGTCCCAATTTCATCAGAGGTTTCGGCTAGA<br>ATCTACCTCCCAAAGATAAGCCAAAAGGAAAAGGAAAAGCTTCCCATTGTGGTCTATTTCC<br>ATGGTGCAGGCTTCTGTCTGGAATCCGCCTACAAGTCATTTTTCCACACTTATGTCAAGCA<br>CTTTGCAGCCGAGGCCAAAGCAATTGCAGTTTCGGTTGAGTTCAGGCTCTCCCCAGAGCA<br>CCACCTGCCTGCAGCTTATGAAGATTGCTGGACTGCCCTTCAGTGGGTGGCTTCACATGT<br>AGATGTTGACAACTCCAGCCTCAAGAATGCTATAGATAAAGAGCCTTGGATAATCAACCAT<br>GGAGACTTTGACAAGGTCTACTTGTGGGGTGATAGTACGGGTGCCAATATTGTGCACAAC<br>GTACTCATCAGAGCTGGTAATGAGAGCTTGCATGGCGGAGTGAAAATCGTGGGTGCAATT<br>CTTTATTACCCATATTTCTTGATCAGGACAAGCTCCAGACAGAGCGATTATATGGAGAACG<br>AGTACAGAGCATACTGGAAGCTGGCTTATCCATCTGCTCCAGGTGGGAACGACAACCCG<br>ATGATAAACCCCGTGGCTGAGAACGCTCCTGATTTGGCCGGATATGGATGTTCGAGGCT<br>GCTGGTATCCATGGTGGCAGACGAGGCCAGAGACATAACCCTTCTCTACATCGAGGCGG<br>TGAAGAAGAGTGGGTGGAAAGGTGAATTGGAGGTGGCTGATTTCGAAGGAGATTACTTTG<br>AAATATTCAGCCCAGAAACTGAGACAGGCAAGAACAAGGTCAAACGTTTAACGTCTTTCAT<br>CAACAAGGACTAA |
| TiCorS<br>(Hydrolase)<br>(MK840854.1) | ATGGCTAATTCAACTGCGAACTCTGATGAGATTGTTTTCGATCTTCATCCATACATCAGAG<br>TCTTTAAAAACGGCAAGGTAGAAAGACTTCACGACACCCCATATGTTCCGCCATCACTTGA<br>AGATCCAGCTACCGGTGTATCCTGGAAAGACGTCCCAATTTCATCCGACGTTTCAGCTAG<br>AGTCTACCTCCCGAAGATCAGCGAAGCGGAAAAGAAAAAGCTCCCCATTTTCGTCTATTT<br>CCATGGTGCAGGCTTCTGTCTGGAATCAGCCTTCAAATCATTTTTCCATACTTATGTTAAG<br>CACGTTGTTGCCGAAACCAAAGCTGTCGGAGTTTCGGTTGAGTACAGACTCGCCCCCGA<br>GCACCCTTTACCTGCGGCTTATGAAGATTGCTGGACTGCCCTTCAGTGGGTGGCTTCCCA<br>TGTTGGTCTTGACAACTCCAGCCTCAAGAATGCTATTGATAAAGAGCCTTGGATAATCAAC<br>CATGGCGACCTCAATAAGCTTTACTTGGGGGGTGATAGTCCTGGTGGAAATATTGTGCAC<br>AACGTACTCATTAGGGCTGGTAAGGAGAGCTTGCATAACGGAGTGAAAATCCGGGGTGC<br>AATTCTTTATTACCCATATTTCTTGATCAGGACAAGCAACAGACAGAGTGATTATATGGAG<br>ATTGACTATAGAGGCTACTGGAAGTTGGCTTATCCATCTGCTCCTGGCGGCACTGACAAC<br>CCAATGATAAACCCTGTAGCTAAGAATGCTCCTGATTTGGCCGGATATGGATGTTCGAGG<br>CTGCTTGTTTCCATGGTTTCGGACGAGACCAGAGATATAACCCTTCTCTACCTTGAGGCAT<br>TGAAGAAGAGTGGGTGGAAAGGTGAATTGGAAGTGGGTGACTACGAAGCACATTTCTTTG<br>ACTTGTTCAGCCCTGAAAATGAAGTTGGCAAGACTTGGATCAAACGTTCAAGCGATTTCAT<br>CAACAAGGAGTAA |
| TiPS (P450)<br>(This study) | ATGGAGCTTCAGAACTTACCCTTTAACTTCTTAGCTTTCATCGTCTTCGTATTCGTTTTCCT<br>AACTCTTAAAGTCTGGAAAAAGTCCAGCCAGGGAAAGCAAAAGCTCCCACCAGGGCCAT<br>GGAAGCTACCTTTGTTAGGAAACCTTCATAATTTGCTGATGGGTTCTCTTCCACATCATAC<br>ACTCAGGGATCTAGCTCGAAAACATGGACCTCTGATGCACCTAAAGCTCGGAGAAGTTAA<br>CGCTCTCATCGTATCATCACCTCGTATGGCTAGGGAGGTGATGAAAACCCATGATCTTGC<br>ATTTGCAAACAGGCCTGTTACTCTGGTTGGGAAGATAGTATGTTATGACTATTCCGATATT<br>GCTTTTAGTCCTTACGGTGATTACTGGCGACAAATGCGCAAAATATGTGTATTGGAGCTTT<br>TTAGCTCCAAGTGTGTTCGATCGTTTGAGCCTATCAGGAAGGATGAAGGCTCTCGTCTGA<br>TTGCCACTCTTCAAGCGTCAGCTGGGAAACCGATAAATCTGACGGAGAAAATCAGCTTGT<br>ATACAACTTCCATGGTCTGCAGAGCAGCATTTGGTAAGGTAAACAACGGGCAAAACAAAT<br>TCTCCCAGTTAGTGAAGGATGCATCAGAGGTAGCAGGAGGCTTTGACCCTACTGATCTAT<br>TTCCATCCTACAAATTTCTTCACGTTCTTGGCAGTACGATGTCTAAATTGCTGAAGATTCAT<br>GGCCAAATAGACGCCATTTTGGAGGAAATGGTTAGTGAGCACAAAAAGAATCATGCCATG<br>TCAAAGAAAGGCAACGGTGAGTATGGTGAAGAAGACATTATTGATATCCTACTCAGAATCA<br>AAGAAGGTGGTGACTTGCAATTTCCATTCACGGACAAGAATGTCAAAGGCATCGTATTTG<br>ATATTTTTGGTGCTGGAACCGAAACTTCATCCTCAGTTGTTGACTGGGCTATGGCAGAATT<br>GATGAGAAATCCAAAAATGATGGCCAAGGCACAAAACGAAGTGCGAGAAGCATTTAAGG<br>GAAAGATGACAATTGATGAAACTGATGTTCGAGGATTGAGTTACATCAAGTCAGTAATCAA<br>AGAAGCTCTGAGGCTACATCCTCCTGTCCCTTTGTTGATTCCCAGAGAATCCAGAGAGCC<br>TTGTCAAATTAATGGCTACGATATACCCCTCAAGACAAGGCTTTTCATCAATGCGTGGGCG<br>ATTTCAAGAGACGAAGAATACTGGCAGGATCCAGAGAGTTTTATTCCTGAGAGATTTGAA<br>GACAGCTCCGTTGATTTTACAGGAACTAACTATGAGTTTGTACCCTTTGGTGCCGGAAGG |

| | |
|---|---|
| | AGGATATGCCCAGGAATGAATTTTGGTCTGGCAAATGTTGATCTTCTTCTAGCTCTGCTAC TCTACCATTTTGACTGGAAACTCCCAGCTGGTCAGAATGATGTAGACATGTCGGAGACCA TTGGTATAGCAGCCAGCAGAAAAAATAGCCTCGTTTTGGTTCCCATTCCTTACGACATTCC ATCTCTTGATAAATCATGCTGA |
| TiT16H (P450) (This study) | ATGGAGTCCATTCTTTTCCTTTGCATCATCCTCCTCCTTTTCTCTATGTTAATGAAAATTAT GAAGAAATCAAAGATCGAGATGGTCTCAAGAAACAAATTGCCCCCAGGGCCGAAACCATT TCCAGTTATAGGAAACATGCATCAGCTCTATGGAGCTCCAATCCACCATGTCCTCAGAGA TTTAGCCAAGAAATATGGTCCTCTTATGCACATCAAGATTGGTGAAATTTCTAGCATTGTT GTCTCCTCGCCAGAAGTTGCCAAAGTGGTTTTCAAGACACATGATCTCCACTTCACTCAAA GACCCTCCAACATTCTAGCAGCGAGGATACTCTCATATGATTTCGCTGACATTACCTTCTC TCCTTATGGAAATTTCTGGAGACAAGTACGGAAAATCAGCACGATTGAACTTTTTAGTCAC AAACGTGTTCAGACATTCCAACCAATTAGGGAAGAGGAGGTGCTGAAACTGATTAGAACC ATTTCGTTGCATGAGGGATCACCGATCAATCTCAGCCAAAAAATTTTTTCATTGACTTATG GAGTCACTACTCGTGCTGCCTTTGGTAGAAGAAACAAATATCAAGAACAATTCCGGGAAC TTTTGGAAAAAATTCAATGTGCTAGCAACAGGATTTAACATAGGAGACATCTTCCCTTCAAA GGAATTCTTTCAAGTGATCAGTGGATTTCGTCCAAAGATGGAAACCCTACACAAGCAGGT TGATGAAATAATTGAAAAAATCCTGATTGAGCACAGGGAGAAACGCAGACAAGACTCCAA TAAGACAACAGAAGAAGCAAGGGAGGATTTAATTGACGTTCTTCTGAATATTCAAGAAGAA GGGAACTTTGAAATTCCTCTTACTGACCGCAACATCAAAGCAATCATCTTTAATGTATTCG GTGCAGGAAGTGAGACATCTTCAGCAGTTGTAGAATGGGCGATGTCGGAACTAATAAAAA ATCCAGGTATGATGAAAAGGGCACAAAACGAAGTTAGAAGTTTCTTCCATGAAAGTGGAA ATGTTAATGAATCGAGCCTTGGACAACTTAAATACTTGCAAGCAATCATTAAAGAAACCAT GAGATTACACCCTAGTGTTCCATTGCTACTCCCCAGGGAATGCAGGGAAGAATGTGAAAT CCAAGGATTTCGGATACCTTCCGGGACCAGAGTCATGGTAAATGCATATGCAATTGGCAG AGATCCTGAATACTGGACTGATGCTGAGAAATTCAGCCCCGACAGATTTCTTGATTCTGAA GTTGATTATAAGGGAAATCATTTCCAGTTTGTACCATTTGGTGCTGGAAGAAGGATGTGTC CAGGGATGTCGTTTGCTCAACCGAATATGGCATTCCCTCTCGCACAATTATTGTTTCATTT TGACTGGAACGTTGATGGCCATATGAAGCAAGAAGATCTAGACATGACCGAGACAGTTGG GGTGACCATTAGAAGGAAAAGTGACTTGTGCCTTATTCCAGTTGTCTATTCAGGTTCTTTT CTTAAATGGAATTCCTAG |
| CrT16H1 (P450) (FJ647194.1) | ATGGAATTCTATTATTTTCTCTACTTGGCCTTCCTTCTTTTCTGCTTCATTTTATCAAAAACC ACAAAGAAATTTGGCCAAAACAGCCAATATTCAAACCATGATGAGCTACCTCCGGGGCCT CCCCAAATTCCTATATTAGGAAATGCCCATCAACTTAGCGGTGGCCATACTCATCACATTC TAAGAGATTTGGCCAAAAAATATGGGCCGTTGATGCACTTAAAGATTGGTGAAGTTTCAAC TATTGTTGCATCTTCACCACAAATTGCTGAAGAGATTTTTAGAACGCATGATATTCTTTTTG CCGATAGACCCTCAAATCTTGAGTCTTTTAAAATCGTGTCTTATGATTTTTCAGATATGGTT GTTAGTCCATATGGTAATTATTGGAGACAACTTCGTAAAATTAGCATGATGGAACTTCTAA GCCAAAAGAGTGTCCAATCTTTTAGATCAATTAGAGAAGAGGAAGTATTAAATTTTATTAAA TCAATTGGTTCCAAAGAGGGTACAAGAATTAATCTCAGCAAAGAGATATCGTTACTTATTT ATGGAATTACTACGCGTGCTGCTTTTGGAGAGAAAAATAAGAATACAGAAGAATTTATTCG TCTACTTGATCAACTTACAAAGGCAGTAGCGGAACCTAACATTGCAGATATGTTCCCTTCT CTCAAATTTCTTCAATTGATTAGTACATCAAAATATAAGATTGAGAAAATACACAAACAATTT GATGTTATAGTTGAAACTATTCTCAAAGGGCATAAGGAGAAAATCAACAAGCCCTTAAGTC AAGAGAATGGAGAAAAAAAGGAGGACCTTGTTGATGTGCTACTCAATATTCAACGACGTA ATGACTTTGAAGCCCCACTGGGGGATAAAAACATCAAAGCCATAATCTTCAACATATTCAG TGCCGGCACTGAGACATCGTCAACAACAGTCGATTGGGCAATGTGCGAAATGATAAAAAA TCCAACGGTAATGAAAAAGGCACAAGAAGAGGTAAGAAAGGTATTTAATGAAGAAGGAAA TGTTGATGAAACAAAACTTCATCAACTAAAATATTTACAAGCAGTGATTAAAGAAACATTAA GGCTTCATCCACCAGTTCCATTACTACTTCCAAGAGAATGTCGAGAACAATGTAAGATTAA AGGGTACACAATACCATCCAAATCTAGAGTTATAGTCAATGCATGGGCTATCGGAAGGGA TCCAAATTACTGGATTGAACCTGAGAAATTTAACCCGGATAGATTTCTTGAATCAAAAGTT GATTTTAAAGGAAATTCATTCGAGTATCTACCATTTGGTGGTGGAAGAAGGATATGTCCGG GCATAACATTTGCTTTGGCTAATATAGAATTGCCATTAGCACAACTTTTGTTCCATTTTGAT TGGCAATCAAATACTGAGAAATTAAATATGAAAGAGAGTAGAGGGGTAACAGTTAGGCGA GAAGATGATTTGTATTTGACTCCAGTTAATTTTTCTTCCTCTTCTCCTGCTTGA |
| CrT16H2 (P450) (JF742645.1) | ATGGAGTTGTATTATTTTTCCACCTTTGCCTTCCTTGTTTTCTGCTTCATTTTAGCCAAAAC TCTAAAGAAATCTGGCCAATCAAATCATAAGCTGCCTCCGGGGCCTCCCCCAATTCCTAT ATTAGGAAATGTCCATCAACTTATCGGTGGCCATACTCATCACGTTCTAAGAGATTTGGCC AAAAAATATGGACCGTTGATGCACTTAAAGACTGGTGAAGTTTCAATCATTGTTGCATCCT CACCAGAAATTGCTGAAGAGATGTTTAAAACACATGATGTTCTCTTTGCGGACAGACCCTC AAATATTGTTGCCTTCAAAATCTTGTCTTATGATTATTCGGATATTGTCATTAGTCCATATG GCAATTATTGGAGACAACTTCGTAAAATTAGCATGATGGAGCTTTTTAGCCAAAGGAGTGT |

| | |
|---|---|
| | CCAATCTTTCAGATCACTTAGAGAAGAGGAAGTCTTGAATTTTATTAAATCAATTGGTTCGA<br>AAGAGGGTACAAAAATTAATCTTAGCAAGGAAATATCGTTACTTATTTATGGAATTACTACG<br>CGTGCTGCTTTTGGAGAAAAAAATAAGAATACAGAAGAATTTATTCGTCTTCTTGATCAACT<br>CACAGTGGCAGTAGCGGAACCTAACATTGCAGATATGTTCCCTTCTATCAATTTTCTTAAA<br>TTAATTAGTAGATCGAAATATAAGATTGAGAAAATACACAAAAATTTTGATGCCATAGTTCA<br>AACTATTCTCAACCATCATAAGGATAGATTAGCCAATTACAAGTCCTCAAGTCCTGAAGAG<br>AATGGGGAACAAAACAAGGATCTTGTTGATGTGCTACTCAATATTCAACAACGTGGTGATT<br>TTGATACACCACTAGGTGATCGCAACGTCAAAGCAGTAATTTTTAACATATTCAGTGCCGG<br>AACTGAGACATCGTCAACGACAGTGGATTGGGCGATGTGTGAAATGATAGAAAATCCAAC<br>GATGATGAAAAAGGCACAAGAAGAGGTAAGAAAGGTATATAACGAAGAAGGAAATGTTAA<br>TGAAACAAAACTTCATCAGCTACAATATTTAAAAGCAGTGATTAAAGAAACATTAAGGCTTC<br>ATCCACCAGTTCCATTACTACTTCCAAGAGAATGTCGAGAACAATGTGAGATTAAAGGGTA<br>CACAATACCATCCAAATCTAGAGTTATAGTCAATGCATGGGCTATCGGAAGGGATTCGAAT<br>TACTGGATTGAACCTGAAAATTTTAACCCGGAGAGATTTCTTGAATCAGAAGTTGATTTTAA<br>AGGAAACTCATTCGAGTATCTACCGTTTGGTGGTGGAAGAAGGATATGTCCGGGCATAAT<br>ATTTGCTTTGGCTAATATAGAACTGCCATTGGCACAACTTTTGTTCCATTTTGATTGGAAAC<br>TTGCCAGTGATGAAACAAATATTGATAAATTAGACATGACGGAGAGTAGAGGAGTAACAG<br>TTAGAAGAGAAGATGATTTGTGTCTGATTCCATTTCCTTATTCTGCTTCTTCTCTCAAAGGT<br>AAATATTGA |
| CrTEX1 (P450)<br>(A0A343URW6.1) | ATGGAGTTTGTGGTTTCCCTCTTTGCCTTCGTCGTTTCCTGCTTCATTTTACTTAAAGTAGC<br>AAAGAATTCCAAGAACCCAAAGAGAAACACAAATCTTGAGCTTCCCCCGGGGCCAAAACA<br>ACTTCCTATAATTGGAAACCTTCATCAGCTTGGCGGCGGCTTAGCTCATCATGTACTTAGA<br>AATTTGGGCAAACAATATGGACCTTTGATGCACTTGAAAATTGGTGAACTTTCAACAATAG<br>TAGTTTCCTCAACAGAAATAGCAAAAGAAGTTTTCAAGACTCATGATATCCACTTTTCTAAT<br>AGACCTTCTCATATTCTTGTTTTTAAAATCGTTAGTTACGATTATAAAGACATTGTTCTTTCT<br>CAATATGGAAAGTATTGGAGAGAGCTTCGAAAAGTTTGTAATCTCGAACTTCTCAGTCCAA<br>ATCGTGTCCAATCCTTCAGATCCATAAGGGAAGATGCTGTCCTTAATATGATGAAATCAAT<br>TTCTTCAAATGATGGCAAAGTTGTCAATTTAAGTGAAATGATCCTCTCTCTTATTTATGGAA<br>TCACTGCACGTGCAGCTTTTGGTGTTTGGAGTAAAAAGCATGAAGAGTTCATAAGACTTGA<br>GAGTGAAATTCAGAGGTTGGCAACAACGTTTGTTTTAGCAGATATGTTTCCTTCGATTAAA<br>TTTCTTGGAGCTCTAAGTGGATTGAGATATAAGGTTGAGAAAGTACATAAAAAGGTTGATG<br>ACATACTTGAAGGCATTCTGAAGGAGCATAGACGACAGAATAATAATATGACAGAAGAAA<br>ATGGAAAAAAGGATCTTGTTGATGTCCTTCTCAACATCCAAAAGAATGGAGACATGGAAAC<br>CCCTTTTACTGATCAACACATCAAAGCCATAATTTTTGATATGTTCAGTGCCGGAACTCTAA<br>CATCAACAATAGCAGTGGATTGGGCAATGGCGGAAATGATGAAGAATCCGAGCGTGCTG<br>AAAAGGGCTCAAGATGAAGTTAGAAATGTGTATAATGGAATTGGAAATGTTGATGAGTCTA<br>AGCTTGATGAATTAAAATATTTACAAGCGGTTATAAAAGAGACATTGAGAATACATCCTGG<br>CACTCCTATTGTTCACAGAGAAACTCGTGAAGAATGTGAAATTAATGGGTATAGGATTCCT<br>GCAAAAGCTAGAGTTATGGTCAATGCATGGGCTATCAGTAGAGATCCGAATTACTGGCCG<br>GAGCCGGATATTTTTAAACCTGAAAGATTTCTAGGGTCGGAAGTTGATTTTAAGGGAACCC<br>ATTTTGAATACATTCCATTTGGAGCAGGAAGGCGTATTTGTCCGGGAATATCATATGCTAT<br>TGCTAATGTTCAACTACCGTTGGCACAGCTTTTGTACCATTTTGAGTGGAAACTTCCAGGC<br>GGAATGAAGCCAGAAGAACTGGACATGACCGAGATTTTGGGTACTGCCGCCCAGCGCAA<br>AGAAAATTTGCTTCTGATACCAAATTCGCATTCTTGTTCTTCTTTGAAACAAGTGTGA |
| CrTEX2 (P450)<br>(A0A343URW7.1) | ATGGAGTTTGTGGTTTCCCCCTTTGCTTTCCTCATTTTCTTCTTCATTTTACTCAAAATGAT<br>AGCAAAGAATTTCAAGAACCCCAAGAAAAACACAAAGCCTCTTCCTCCAGGGCCAAAAAA<br>GCTTCCTATAATAGGAAACCTTCATCAGCTAGGTGGTGGTTTAGCTCACCATATCCTTAGA<br>GATTTATCCCAAAACTATGGACCTTTGATGCATTTGAAAATCGGTGAACTTTCAACAATAGT<br>CATTTCCTCAACAGAAATGGCTAAACAAGTTTTCAAGGTTCATGATATCCATTTCTCAAACA<br>GACCTTCTCATATTCTTGTTTTTAAAATCGTTAGCTATGATTATAAAGACATCGTACTTTCC<br>CAATATGGAAACTACTGGCGAGAGCTTCGTAAAGTCTGTAATCTCCATCTTTTAAGTCCAA<br>ATCGTGTCCAATCCTTTAGATTCATAAGGGAAGATTCTGTTCTTAATATGATGAAATCAATT<br>TCTTCCAACGAGGGTAAAGTTGTCAATTTAAGTGAAATGATTTTATCTCTTATTTATGGAAT<br>TACTGCTCGTGCTGCATTTGGTGTTTGGAGTAAAAGACATGAAGAATTCATAAGACTTGAG<br>AGTGAAATTCAAAGGTTGGCAACAACGTTTGTTTTAGCAGATATGTTTCCTTCGATTAAAAT<br>TCTTGGAGCTTTAAGTGGATTGAGATATAAGGTTGAGAAAGTACATAAAAAAGTTGATGAA<br>ATTCTTGAAGATATTCTCAAAGAGCATAGAAATAATAATATTAGTATTGAGAAAGAAGAAGA<br>AGAAGAAGAAGAAAATAATGGTGGAAAAAAGGATCTTGTTGATGTCCTTCTTGATATCCAA<br>AAGAATGGAGAAATGGAAACCCCATTTACAGATCAACATATCAAAGCCATCATTTTTGATA<br>TGTTCAGTGCCGGAACCCTAACGTCGACGATAGCAGTGGATTGGGCAATGGCCGAAATG<br>ATGAAGAATCCAAGAGTGATGAAAAGGGCTCAAGAAGAAGTTAGGAATGTATACAACGGA<br>ATAGGAAACGTAAACGAATCAAAACTCGATGAATTAAAATATTTACAAGCAATTATAAAAGA |

| | |
|---|---|
| | GACACTAAGAATACATCCAGGAACTCCCATAGTTCATAGAGAAACTCGTGAAGAATGCGA<br>GATTAATGGGTATAGAATACCTGCAAAAGCTAGAGTTATGGTTAATGCATGGGCAATCAGT<br>AGAGATCCAAATTACTGGCCAGATCCGGATAGCTTTAAACCTGAGAGATTTTTGGGATCA<br>GAAGTCGATTTTAAAGGTACCCATTTTGAATATACTCCATTTGGAGCAGGTAGGCGTATTT<br>GTCCGGGGATATCATATGCCATTGCCAATATTCAGCTGCCGTTGGCGCAACTTTTGTACC<br>ATTTTGATTGGAAACTTGCCGGTGGAATGAAGCCTGAAGAATTGGACATGGCGGAGATTT<br>TGGGTACTGCTGCCCAACGCAAAGAAGATTTGCTTCTCATACCAAATTCCCATTCTTGTTC<br>TTCTTTGAAACAACAAGTGTGA |
| CrT19H (P450)<br>(HQ901597.1) | ATGTTGTCTTCATTGAAAGATTTCTTCGTTCTACTTCTGCCTTTCTTCATTGGCATTGCATT<br>TATATATAAATTATGGAATTTTACATCCAAGAAAAATCTTCCGCCATCGCCAAGAAGGCTA<br>CCGATTATCGGAAACCTTCATCAGCTCAGTAAATTTCCTCAACGTTCACTGAGAACATTAT<br>CAGAAAAATATGGGCCTGTCATGTTACTTCATTTTGGTAGCAAGCCAGTTTTGGTTATATC<br>CTCAGCAGAAGCAGCTAAAGAAGTAATGAAAATCAATGATGTTTCTTTCGCTGATAGGCCT<br>AAATGGTATGCTGCTGGTAGGGTTCTTTATGAATTTAAGGATATGACATTTTCACCCTATG<br>GTGAGTATTGGAGACAAGCTAGAAGTATATGTGTTCTTCAGCTTCTAAGTAACAAAAGGGT<br>TCAATCTTTTAAAGGTATTAGAGAAGAAGAGATAAGGGCAATGTTGGAAAAAATAAATCAA<br>GCTAGTAATAATTCAAGTATCATTAATGGTGATGAGATTTTCTCGACACTTACAAATGACAT<br>AATTGGAAGATCTGCTTTTGGGAGGAAATTTTCTGAAGAAGAAAGTGGAAGTAAATTAAGA<br>AAAGTTCTTCAAGATTTGCCACCTTTGTTAGGTTCTTTCAATGTTGGTGATTTTATTCCATG<br>GCTTTCTTGGGTTAATTATTTAAATGGTTTTGAAAAAAAATTGAACCAAGTTTCAAAAGATT<br>GTGATCAATATCTTGAGCAAGTTATTGATGATACGAGGAAGAGAGATGAAGAAAATGGAG<br>CTAATAACAATGGCGGAAATCATGGAAATTTTGTTAGTGTTCTACTTCATCTCCAGAAAGA<br>AGATGTCAAGGGTTTTCCTTCAGAAAAAGATTTTCTTAAAGCTATCATTTTGGACATGATTG<br>TTGGGGGAACAGACACAACACATTTGCTTTTGCATTGGGTAATCACAGAGCTTCTCAAAAA<br>CAAACATGTAATGACTAAATTACAAAAAGAGGTGAGAGAAATAGTCGGAAGAAAATGGGA<br>GATAACTGATGAAGACAAAGAAAAAATGAAATATTTACACGCAGTTATAAAAGAGGCACTC<br>CGGTTACATCCTTCTCTTCCTTTACTCGTTCCCAGAGTAGCAAGAGAGGATATAAACCTAA<br>TGGGTTATCGCGTAGCGAAAGGCACAGAAGTGATTATTAATGCTTGGGCAATAGCGAGAG<br>ATCCATCGTACTGGGATGAAGCCGAGGAGTTTAAGCCGGAGAGATTCTTGAGTAATAATT<br>TTGATTTTAAAGGACTTAATTTTGAGTATATTCCATTTGGTTCTGGAAGAAGAAGTTGCCCT<br>GGATCTTCATTTGCAATCCCCATTGTAGAACACACAGTGGCACATTTGATGCATAAATTCA<br>ATATTGAGTTGCCAAATGGAGTCAGTGCTGAAGATTTTGATCCCACAGATGCCGTTGGAC<br>TCGTATCTCATGACCAAAACCCTCTCTCGTTTGTGGCAACTCCGGTTACCATTTTTGA |
| CrT3O (P450)<br>(JN613015.1) | ATGGAGTTTCATGAATCTTCTCCCTTCGTGTTCATCACTCGTGGCTTTATATTCATAGCAAT<br>TTCAATAGCCGTACTGAGAAGAATAATATCAAAGAAAACTAAAACATTGCCTCCAGGACCA<br>TGGAAGCTACCTTTGATTGGAAATCTTCATCAATTTCTGGGTTCTGTTCCTTATCAAATTCT<br>CAGAGATTTAGCTCAAAAAAATGGGCCTTTGATGCACCTTCAATTAGGTGAAGTTTCTGCC<br>ATCGTGGCAGCATCTCCTCAAATGGCTAAGGAGATTACAAAAACTTTGGATCTTCAATTTG<br>CAGACAGGCCAGTTATTCAAGCATTAAGGATTGTGACCTATGATTATTTAGATATATCCTTT<br>AATGCATACGGAAAATATTGGAGACAATTGCGTAAAATTTTTGTCCAAGAACTATTAACTTC<br>AAAGAGAGTTCGATCATTTTGCTCTATAAGAGAAGATGAATTTTCCAATCTGGTAAAAACA<br>ATCAATTCTGCGAATGGAAAATCAATCAATTTGAGCAAATTGATGACGTCATGCACAAATT<br>CAATTATTAATAAAGTAGCTTTTGGTAAAGTACGTTATGAACGGGAGGTGTTTATTGATCTA<br>ATTAATCAAATATTAGCATTAGCAGGCGGTTTTAAGCTGGTTGATCTGTTTCCGTCCTACA<br>AGATACTTCATGTTCTTGAAGGTACAGAACGTAAGCTGTGGGAAATCCGCGGTAAGATTG<br>ACAAGATTTTGGATAAAGTCATAGACGAGCACAGAGAAAATTCGTCAAGAACTGGAAAGG<br>GCAACGGTTGTAATGGCCAGGAAGATATAGTTGATATTTTACTTAGGATTGAAGAGGGTG<br>GTGATCTTGACCTTGATATTCCCTTTGGCAACAACAATATCAAAGCTCTTTTATTCGATATA<br>ATTGCAGGCGGAACTGAAACCTCATCAACAGCAGTTGACTGGGCAATGTCAGAGATGATG<br>AGAAATCCCCATGTGATGAGCAAAGCGCAAAAGGAAATTAGGGAAGCGTTCAATGGAAAG<br>GAGAAGATTGAGGAGAATGATATTCAAAATTTGAAGTACCTAAAGTTAGTGATCCAAGAAA<br>CCTTAAGGTTACACCCTCCTGCTCCATTGTTGATGAGACAATGCCGAGAGAAATGTGAAA<br>TTGGCGGATATCATATACCTGTTGGAACAAAAGCGTTCATCAATGTCTGGGCAATCGGAA<br>GGGATCCTGCGTATTGGCCTAATCCAGAGAGTTTTATTCCGGAAAGATTTGACGATAATAC<br>TTATGAATTTACAAAATCTGAACATCATGCGTTTGAATATTTGCCATTTGGTGCCGGAAGA<br>AGGATGTGTCCGGGCATTTCATTTGGTTTGGCCAACGTGGAGCTTCCTTTAGCTCTACTT<br>CTTTACCATTTCAACTGGCAACTCCCAGATGGTTCTACTACTCTGGATATGACAGAGGCTA<br>CTGGATTAGCAGCAAGAAGAAAATATGATCTTCAATTAATCGCTACGTCCTATGCATGA |

**Table S2** Primers used in this study.

| **Gene** | **Plasmid** | **Primer direction** | **Sequence (5'-3')** |
|---|---|---|---|
| Yeast expression constructs (In-Fusion cloning) | | | |
| TiPS | pESC-HIS | Forward | **GAGAAAAAACCCCGGATCC**ATGGAGCTTCAGAACTTACC |
| | | Reverse | **ACTTCTGTTCCATGTCGAC**GCATGATTTATCAAGAGATGGAA |
| TiT16H | pESC-HIS | Forward | **GAGAAAAAACCCCGGATCC**ATGGAGTCCATTCTTTTCCTTT |
| | | Reverse | **ACTTCTGTTCCATGTCGAC**CTAGGAATTCCATTTAAGAAAAGAA |
| CrT16H1 | pESC-HIS | Forward | **GAGAAAAAACCCCGGATCC**ATGGAATTCTATTATTTTCTCTACTTG |
| | | Reverse | **ACTTCTGTTCCATGTCGAC**AGCAGGAGAAGAGGAAGAAA |
| CrT16H2 | pESC-HIS | Forward | **GAGAAAAAACCCCGGATCC**ATGGAGTTGTATTATTTTTCCAC |
| | | Reverse | **ACTTCTGTTCCATGTCGAC**ATATTTACCTTTGAGAGAAGAAG |
| CrTEX1 | pESC-HIS | Forward | **GAGAAAAAACCCCGGATCC**ATGGAGTTTGTGGTTTCCCT |
| | | Reverse | **ACTTCTGTTCCATGTCGAC**CACTTGTTTCAAAGAAGAACAAGAA |
| CrTEX2 | pESC-HIS | Forward | **GAGAAAAAACCCCGGATCC**ATGGAGTTTGTGGTTTCCC |
| | | Reverse | ACTTCTGTTCCATGTCGACCACTTGTTGTTTCAAAGAAGAAC |
| CrT19H | pESC-HIS | Forward | **GAGAAAAAACCCCGGATCC**ATGTTGTCTTCATTGAAAGATTTCTT |
| | | Reverse | **ACTTCTGTTCCATGTCGAC**AAAAATGGTAACCGGAGTTGC |
| CrT3O | pESC-HIS | Forward | **GAGAAAAAACCCCGGATCC**ATGGAGTTTCATGAATCTTC |
| | | Reverse | **ACTTCTGTTCCATGTCGAC**TGCATAGGACGTAGCGATTA |
| Sequence verification primers pESC-HIS | | Forward | ATGATTTTTGATCTATTAACAGATA |
| | | Reverse | GTATAATGTTACATGCGTACAC |
| **Gene** | **Plasmid** | **Primer direction** | **Sequence (5'-3')** |
| *Nicotiana benthamiana* single transcriptional unit constructs (In-Fusion cloning) | | | |
| P19 (TBSV) | 3Ω1 | Forward | **TTTATGAATTTTGCAGCTCG**ATGGAACGAGCTATACAAGGAAA |
| | | Reverse | **GACAACCACAACAAGCACCG**TCACTCGCTTTCTTTTTCGAAGG |
| GFP | 3Ω1 | Forward | **TTTATGAATTTTGCAGCTCG** ATGGTGAGCAAGGGCGAG |
| | | Reverse | **GACAACCACAACAAGCACCG**TCACTTGTACAGCTCGTCCATGC |
| TiPAS3 | 3Ω1 | Forward | **TTTATGAATTTTGCAGCTCG**ATGTTAGCAGAAGTCTCCAAAGTTC |
| | | Reverse | **GACAACCACAACAAGCACCG**TTACAATTCATCATGTAAAGTTAGAG |
| TiDPAS1 | 3Ω1 | Forward | **TTTATGAATTTTGCAGCTCG**ATGGCTGTAAAATCACCTGAAGC |
| | | Reverse | **GACAACCACAACAAGCACCG**CTATTCCGGTGGAGTTAGTGTG |
| TiCorS | 3Ω1 | Forward | **TTTATGAATTTTGCAGCTCG**ATGGCTAATTCAACTGCGAACTCT |
| | | Reverse | **GACAACCACAACAAGCACCG**TTACTCCTTGTTGATGAAATCGCTTG |
| TiTabS | 3Ω1 | Forward | **TTTATGAATTTTGCAGCTCG**ATGGCTTCTTCAACTGAAAGCTCTG |
| | | Reverse | **GACAACCACAACAAGCACCG**TTAGTCCTTGTTGATGAAAGACGTTAG |
| TiPS | 3Ω1 | Forward | **TTTATGAATTTTGCAGCTCG**ATGGAGCTTCAGAACTTACC |
| | | Reverse | **GACAACCACAACAAGCACCG**TCAGCATGATTTATCAAGAGATGGAA |
| TiT16H | 3Ω1 | Forward | **TTTATGAATTTTGCAGCTCG**ATGGAGTCCATTCTTTTCCTTT |
| | | Reverse | **GACAACCACAACAAGCACCG**CTAGGAATTCCATTTAAGAAAAGAA |
| CrT16H1 | 3Ω1 | Forward | **TTTATGAATTTTGCAGCTCG**ATGGAATTCTATTATTTTCTCTACTTG |
| | | Reverse | **GACAACCACAACAAGCACCG**TCAAGCAGGAGAAGAGGAAGAAA |
| CrT16H2 | 3Ω1 | Forward | **TTTATGAATTTTGCAGCTCG**ATGGAGTTGTATTATTTTTCCAC |
| | | Reverse | **GACAACCACAACAAGCACCG**TCAATATTTACCTTTGAGAGAAGAAG |
| CrTEX1 | 3Ω1 | Forward | **TTTATGAATTTTGCAGCTCG**ATGGAGTTTGTGGTTTCCCT |
| | | Reverse | **GACAACCACAACAAGCACCG**TCACACTTGTTTCAAAGAAGAACAAGAA |
| CrTEX2 | 3Ω1 | Forward | **TTTATGAATTTTGCAGCTCG**ATGGAGTTTGTGGTTTCCC |
| | | Reverse | **GACAACCACAACAAGCACCG**TCACACTTGTTGTTTCAAAGAAGAAC |
| CrT19H | 3Ω1 | Forward | **TTTATGAATTTTGCAGCTCG**ATGTTGTCTTCATTGAAAGATTTCTT |
| | | Reverse | **GACAACCACAACAAGCACCG**TCAAAAAATGGTAACCGGAGTTGC |
| CrT3O | 3Ω1 | Forward | **TTTATGAATTTTGCAGCTCG**ATGGAGTTTCATGAATCTTC |
| | | Reverse | **GACAACCACAACAAGCACCG**TCATGCATAGGACGTAGCGATTA |

| Gene | Level 1 CDS | Primer direction | Sequence (5'-3') |
|---|---|---|---|
| Sequence verification primers 3Ω1 InFusion | | Forward | GATGAAAAAGCCCTAAAATTGGAG |
| | | Reverse | ATTATTCACAAATGAGAAACAGAATGG |
| **Gene** | **Level 1 CDS** | **Primer direction** | **Sequence (5'-3')** |
| *Nicotiana benthamiana* multi-transcriptional unit constructs | | | |
| Level 1 CDS fragment for 3α1 or 3α2 golden-braid assembly | | | |
| Domestication of TiDPAS1 | Frag. A | Forward | GCGCCGTCTCGCTCGAATGGCTGTAAAATCACCTGAAG |
| | | Reverse | GCGCCGTCTCGCAGTCTCATTCGAGAAGCCT |
| | Frag. B | Forward | GCGCCGTCTCGACTGTCGTAAATGAGCGCTTC |
| | | Reverse | GCGCCGTCTCGCTCGAAGCCTATTCCGGTGGAGTTAGTG |
| | Full length | Forward | GCGCCGTCTCGCTCGAATGGCTGTAAAATCACCTGAAG |
| | | Reverse | GCGCCGTCTCGCTCGAAGCCTATTCCGGTGGAGTTAGTG |
| Domestication of TiCorS | Frag. A | Forward | GCGCCGTCTCGCTCGAATGGCTAATTCAACTGCGAAC |
| | | Reverse | GCGCCGTCTCGTAGTCTCGTCCGAAACCATG |
| | Frag. B | Forward | GCGCCGTCTCGACTAGAGATATAACCCTTCTCTAC |
| | | Reverse | GCGCCGTCTCGCTCGAAGCTTACTCCTTGTTGATGAAATCGC |
| | Full length | Forward | GCGCCGTCTCGCTCGAATGGCTAATTCAACTGCGAAC |
| | | Reverse | GCGCCGTCTCGCTCGAAGCTTACTCCTTGTTGATGAAATCGC |
| TiPAS3 | Full length | Forward | **TTCAGAGGTCTCTA**ATGTTAGCAGAAGTCTCCAAAG |
| | | Reverse | **AGCGTGGGTCTCGAAGC**TTACAATTCATCATGTAAAGTTAGAG |
| TiTabS | Full length | Forward | **TTCAGAGGTCTCTA**ATGGCTTCTTCAACTGAAAGCT |
| | | Reverse | **AGCGTGGGTCTCGAAGC**TTAGTCCTTGTTGATGAAAGACG |
| P19 | Full length | Forward | **TTCAGAGGTCTCTA**ATGGAACGAGCTATACAAGGAAA |
| | | Reverse | **AGCGTGGGTCTCGAAGC**TCACTCGCTTTCTTTTTCGAAGG |
| Plasmid validation | P19 | Forward | TATTTAAGAGATATCTCAGATACG |
| | | Reverse | CTGGTCGAAACCGAAAAATC |
| | TiPAS3 | Forward | GCACTTCAGTTGTGGTGG |
| | | Reverse | TCTTCGTCCGATGAGAACT |
| | TiDPAS1 | Forward | CGTAAATGAGCGCTTCGTT |
| | | Reverse | TGGGAGTGGAACTAATCACT |
| | TiTabS | Forward | CCACCTGCCTGCAGCTTAT |
| | | Reverse | GGGTAATAAAGAATTGCACCC |
| | TiCorS | Forward | CCTTCAGTGGGTGGCT |
| | | Reverse | CAATCTCCATATAATCACTCTGT |
| Overhangs designed for non-domesticated sequences 3α1 or 3α2 golden-braid assembly | | | |

**Methods S1** Detailed Materials and Methods

**Candidate selection and phylogenetic analysis**

The candidate genes were selected from a previously generated *T. iboga* transcriptome (Farrow *et al.*, 2019). Full-length cytochromes P450 gene sequences were extracted from the transcriptome using annotations corresponding to cytochrome P450, CYP, and EC:1.14. Transcripts were filtered to remove duplicates and include CDS (coding sequence) between 400 and 550 amino acids. Multiple sequence alignment (amino acid) was constructed using MUSCLE (Edgar, 2004) using default parameters, including 245 *Arabidopsis thaliana* cytochromes P450s (Schuler *et al.*, 2006) and functionally characterized cytochromes P450 involved in monoterpene indole alkaloid biosynthesis (Nguyen & Dang, 2021; Williams & Luca, 2023). The alignment was used to construct a maximum-likelihood phylogenetic tree using the iQtree web server (Trifinopoulos *et al.*, 2016) under default parameters. Phylogenetic trees were visualized using iToL (Letunic & Bork, 2024) and Geneious Prime ver. 2023.1.2. Major clades in the tree were identified and annotated based on *Arabidopsis thaliana* sequences, including the CYP71 family and Clan 86, the former used to identify candidates and the latter used to root the tree (Hansen *et al.*, 2021).

**Cloning methods**

Cytochrome P450 enzymes characterized in this study were amplified by PCR from a cDNA library generated from *T. iboga* as previously described (Farrow *et al.*, 2019). Known *C. roseus* cytochromes P450 genes (Table S1) were amplified from a cDNA library generated from *C. roseus* var. LBE. RNA was isolated from *C. roseus* var. LBE using the RNeasy Plant Mini Kit (Qiagen) and gDNA digestion and cDNA synthesis were performed using SuperScript IV-VILO Master-Mix (ThermoFisher). Overhangs compatible with the appropriate vector for yeast expression or plant transient expression (*N. benthamiana* or *C. roseus*) were included on the 5' and 3' end of the primers used for PCR amplification (Table S2). HiFi Q5 Hot-Start 2X Master Mix (New England Biolabs) was used for gene amplification. PCR products were analyzed on 1% agarose gel and purified using the Zymoclean Gel DNA Recovery Kit (Zymo Research). In-Fusion HD cloning kit (Takara Bioscience) was used to assemble the amplified genes into target vectors.

For yeast expression, amplicons were cloned into the MCS2 of pESC-HIS ($Kan^R$) plasmid (GenBank: AF063850.1, Agilent). The plasmid was linearized using *SalI* and *BamHI* restriction enzymes (New England Biolabs). The In-Fusion assembly products were transformed into *E. coli* TOP10 cells (ThermoFisher) by heat-shock. Positive transformants were selected on LB agar (50 µg $ml^{-1}$ kanamycin), and colonies were screened by colony PCR using Phire Hot Start II 2X Master Mix (ThermoFisher). Plasmid DNA was isolated from positive colonies using Wizard Plus SV Minipreps DNA Purification Kit (Promega) and verified by Sanger sequencing (pESC-HIS sequencing primers, Table S2). Sequence-verified constructs were then transformed into yeast strain *Saccharomyces cerevisiae* WAT11 (*ade2*; contains the *Arabidopsis thaliana* cytochrome P450 reductase I gene, *ATR1*) (Urban *et al.*, 1997) using the Frozen-EZ Yeast Transformation II Kit (Zymo Research). Transformed yeast cells were plated on SD-His medium (6.7 g $l^{-1}$ yeast nitrogen base without amino acids, 2 g $l^{-1}$ drop-out mix without histidine, 74 mg $l^{-1}$ adenine hemisulfate) containing 2% glucose (w/v) and grown at 30 °C for 48 hours. Positive transformants were confirmed by colony PCR as described above and used to inoculate 10 ml of SD-His medium + 2% glucose (w/v), which were grown at 30 °C, 220 rpm for 48 hours. These cultures were then used to store 25% glycerol stocks at -80 °C for further use.

For transient expression in *N. benthamiana* or *C. roseus*, all single transcriptional unit plasmids were generated by cloning the amplicons (Table S2) directly into a modified 3Ω1 (Sarrion-Perdigones *et al.*, 2013) vector. The 3Ω1 binary vector modification includes a *CCDB* gene cassette between ubiquitin-10 promoter (pUbq10) and ubiquitin-10 terminator (tUbq10) instead of *LacZ* (present in classical 3Ω1 plasmid). The vector was linearized using *BsaI* restriction enzyme (New England Biolabs), and the amplicon assembly was carried out using an In-Fusion HD cloning kit (Takara Bioscience). The assembled product of In-Fusion was transformed into *E. coli* TOP10 (ThermoFisher) cells. Positive transformants were identified by colony PCR as described above (3Ω1 sequencing primers, Table S2) and then grown overnight in liquid LB medium (200 µg $ml^{-1}$ spectinomycin) at 37 °C, 200 rpm. Plasmid DNA was isolated, and the sequence was verified as described above. Sequence-verified constructs were used to transform electrocompetent *Agrobacterium tumefaciens* GV3101 cells (Goldbio) by electroporation. Cells were plated on LB medium (20 µg $ml^{-1}$ rifampicin, 30 µg $ml^{-1}$ gentamicin, 200 µg $ml^{-1}$ spectinomycin) containing agar (20 g $l^{-1}$) and grown at 28 °C for 48 hours. Positive colonies were identified by colony PCR, and a single transformant was grown in 10 ml LB medium (20 µg $ml^{-1}$ rifampicin, 30 µg $ml^{-1}$ gentamicin, 200 µg $ml^{-1}$ spectinomycin) at 28 °C, 250 pm overnight, after which 25% glycerol stocks were prepared and stored at -80 °C for further use.

Multi-transcriptional unit constructs for plant transient expression in *N. benthamiana* or *C. roseus* were assembled using the Goldenbraid 2.0 toolkit (Sarrion-Perdigones *et al.*, 2013). Geneious Prime ver. 2023.1.2 was used to construct in-silico assemblies before cloning. Domestication (removal of *BsaI* and *BsmbI* cut-sites) of TiDPAS1 and TiCorS was performed by PCR according to primers generated by the Goldenbraid domesticator web tool (Sarrion-Perdigones *et al.*, 2013) (Table S2). All CDS elements used for multi-transcriptional unit assemblies were cloned into a Level 1 transcriptional unit in 3α1 or 3α2 plasmid (kanamycin) using *BsaI* (New England Bioscience). Level 2 multi-gene constructs were assembled into 3Ω1 or 3Ω2 (spectinomycin) plasmid with *Esp3I* (New England Bioscience) using Level 1 plasmids. Level 3 multi-gene constructs were assembled into 3α1 plasmid with *BsaI* (New England Bioscience) using Level 2 plasmids (Fig. S1). All transcriptional units consisted of a *Solanum lycopersicum* Ubiquitin10 promoter (pUbq10) and Ubiquitin 10 terminator (tUbq10, Table S1). The assembly was carried out by incubating the reaction mix at 37 °C for 5 min, followed by 5 min at 16 °C for 50 cycles, and terminated at 65 °C for 10 min. Plasmids were transformed into *E. coli* TOP10 (ThermoFisher) cells by heat shock. Blue-white screening was used to identify positive transformants. Sequence verification was performed at each stage of the assembly by Sanger sequencing using gene-specific primers (Table S2), and the final multi-transcriptional unit (Level 3) construct was verified using whole plasmid sequencing (Plasmidsaurus). Sequence-verified constructs were transformed into *A. tumefaciens* GV3101 cells (Goldbio) by electroporation. Transformed cells were grown, verified, and maintained in glycerol stocks described above in LB medium (20 µg $ml^{-1}$ rifampicin, 30 µg $ml^{-1}$ gentamicin, 100 µg $ml^{-1}$ kanamycin).

**Heterologous expression in yeast and microsome preparation**

Yeast cells (WAT11) transformed with expression constructs including empty vector control were streaked on bacteriological agar (20 g $l^{-1}$) plates of SD-His medium (6.7 g $l^{-1}$ yeast nitrogen base without amino acids, 2 g $l^{-1}$ drop-out mix without histidine, 74 mg $l^{-1}$ adenine hemisulfate) containing 2% glucose (w/v) from glycerol stocks and incubated at 30 °C for 48 hours for colonies to appear. Colonies were inoculated in 10 ml SD-His + 2% glucose medium and grown overnight at 30 °C, 200 rpm. A cell density ($OD_{600}$) corresponding to 1.0 was sub-cultured into 100 ml of SD-His medium + 2% glucose and incubated for 28-34 hours at 30 °C, 200 rpm. Cells were harvested at 4000 × g, room temperature for 5 min. Harvested cells were resuspended in

100 ml of SD-His medium + 1.8% galactose (w/v) + 0.2 % glucose (w/v) and incubated for 18-24 hours at 30 °C, 200 rpm for protein induction. Yeast culturing and handling were performed while maintaining aseptic conditions to avoid contaminations.

After protein expression, cells were harvested by centrifugation (4000 × g, 4 °C, 10 min) for yeast microsome preparation (Pompon *et al.*, 1996). The cell pellet was resuspended in 10 ml (2 ml $g^{-1}$) of TEK buffer (50 mM Tris-HCl, 1 mM EDTA, 100 mM potassium chloride, pH 7.4) and incubated at room temperature for 5 min. Cells were then harvested by centrifugation as above, and the supernatant was discarded; the pellet was resuspended in 2 ml of ice-cold TES buffer (50 mM Tris-HCl, 1 mM EDTA, 600 mM sorbitol, 10 g $l^{-1}$ bovine serum albumin, pH 7.4). Cells were disrupted by glass beads (0.5 mm diameter, equal volume 2 ml) in a cold room (4 °C) using a Bead Genie (Scientific Industries) by pulsation at six rounds of 5000 rpm for 1 min on, 1 min off intervals. Following cell lysis, 5 ml of ice-cold TES buffer was added to the lysate/glass bead suspension, mixed well, and the supernatant was collected into a pre-chilled tube. The glass beads were washed three times and pooled for centrifugation at 8000 × g, 4 °C, 10 min. The supernatant was collected and ultra-centrifuged at 100,000 × g, 4 °C, 90 min. The supernatant was discarded, and the translucent pellet was washed first with 1 ml ice-cold TES buffer, then with 1 ml of TEG buffer (50 mM Tris-HCl, 1 mM EDTA, 20% glycerol, pH 7.4), and resuspended in 1 ml TEG buffer using a Dounce homogeniser. Microsomal protein preparations were stored at –80 °C for in vitro assays.

**SPE purification of products of enzymatic reactions**

Solid-phase extraction (SPE) was performed to remove impurities and concentrate the sample before HPLC isolation. For yeast and *N. benthamiana* sample clean-up, a 3 ml Chromabond-HLB (60 µm, 3 ml, 60 mg Macherey-Nagel) and a 6 ml Chromabond-HLB (60 µm, 6 mL, 200 mg, Macherey-Nagel) SPE cartridge was used, respectively. The sample, reconstituted in 10% methanol (aqueous), was applied to a conditioned SPE cartridge for processing. The SPE cartridge was first conditioned with three-column volumes of methanol and three-column volumes of 10% methanol (aqueous). The sample was then applied to the SPE cartridge, washed with three-column volumes of 20% methanol (aqueous), and dried under vacuum. Finally, the reaction product was eluted with one column volume of methanol, and the flow-through was evaporated to dryness for semi-preparative HPLC isolation.

**Semi-preparative HPLC isolation of enzymatic products**

Semi-preparative-scale reaction workups of (–)-pachysiphine, (–)-16OH-tabersonine, and (–)-16OH-pachysiphine were subjected to high-performance liquid chromatography (HPLC) for compound isolation. The *N. benthamiana* extract containing pseudotabersonine was also subjected to HPLC isolation. An Agilent 1260 Infinity II HPLC instrument connected to an autosampler, diode array detector (DAD), and fraction collector for compound detection and isolation. Chromatographic separation was performed using a Phenomenex Kinetex XB-C18 (5.0 µm, 100 Å, 100 × 2.1 mm) column maintained at 40 °C under gradient elution using reversed phase conditions. The mobile phases used for separation were water with 0.1% formic acid (A) and acetonitrile (B). The flow rate was set at 1.2 ml $min^{-1}$, and chromatographic separation was performed at 10% B for 2 min, followed by a linear gradient from 10% to 30% B in 12 min, 90% B for 3 min, 10% B for 3 min ($t_{total}$ 20 min). Prior to injection, the samples were diluted to 1 mg $ml^{-1}$ with methanol and filtered using a 0.22 µm PTFE syringe filter. The diluted samples were placed in the autosampler, 50 ul injections were performed, and fractions were collected by monitoring the UV 254 nm and 328 nm corresponding to aspidosperma alkaloid absorbance (Hisiger & Jolicoeur, 2007). Fractions were analyzed by LC-MS to confirm identity and verified fractions were pooled and evaporated to dryness. The isolated compounds were then submitted for NMR analysis. In the case of pseudotabersonine, the isolated compound was analyzed further by chiral LC-MS.

**LC-MS analysis.**
All samples were analyzed using an UltiMate 3000 (Thermo Scientific) ultra-high performance liquid chromatography (UHPLC) system coupled to an Impact II UHR-Q-ToF (Ultra-High-Resolution Quadrupole-Time-of-Flight) mass spectrometer (Bruker Daltonics). Metabolites were separated by reversed-phase liquid chromatography using a Phenomenex Kinetex XB-C18 (100 x 2.1mm, 2.6 µm, 100 Å) column at 40 °C. The mobile phases for metabolite separation were water with 0.1% formic acid (A) and acetonitrile (B). A flow rate of 0.6 ml $min^{-1}$ was used for the chromatography. The sample injection volume was 2 µl. The chromatographic separation started at 10% B for 1 min, and the linear gradient was from 10% to 30% B in 5 min, 90% B for 1.5 min, and 10% B for 2.5 min ($t_{total}$ 10 min). Authentic standards were prepared as 20 µM solutions in methanol, and 2 µl were injected under the chromatographic conditions described above. Mass spectrometry acquisition was performed in positive electrospray ionization (+ ESI) mode with a capillary voltage of 3500 V and an end plate offset of 500 V; a nebulizer pressure of 2.5 bar was used, with nitrogen at 250 °C and a flow of 11 l $min^{-1}$ as the drying gas. Acquisition was set at 12 Hz in the mass range from *m/z* 80 to 1000, with data-dependent $MS^2$ and an active exclusion after three spectra, released after 0.2 min. Fragmentation was triggered on an absolute threshold of 400 counts and limited to a total cycle time range of 0.5 seconds. The collision energies were applied dynamically and stepped from 20 to 50 eV. At the beginning of each sample run, a sodium formate-isopropanol calibration solution was directly infused into the source at 0.18 ml $h^{-1}$ using an external syringe pump to calibrate the MS spectra recorded. To avoid injection peak and salt contamination of the MS, the initial 1 min of the active chromatographic gradient of each run was discarded to waste, and the calibration solution was directed to the MS during this time. At the end of the active LC gradient (6 min), the MS valve was switched to waste, and during re-equilibration time (2.5 min), the calibration solution was directed to the MS. Data analysis was performed using Bruker Compass Data Analysis (Version 5.3) software.

**Structure optimization and ECD spectral calculation for (–)-pachysiphine.**
Electronic circular dichroism (ECD) spectra calculation was performed for (–)-pachysiphine. Based on the structure determined from NMR analysis, a molecular model was created in GaussView ver.6 (Semichem Inc., Shawnee, Kansas, USA) and optimized using the semi-empirical method PM6 in Gaussian (Gaussian Inc., Wallingford, Connecticut, USA). The resulting structure was used for conformer variation with the GMMX processor of the Gaussian program package. The resulting structures were DFT-optimized with Gaussian ver.16 (APFD/6-31G(d) level). A 4 kcal/mol cut-off level was used to select conformers subjected to another DFT optimization on a higher level (APFD/6-311G+(2d,p)). The lowest energy conformer from the high-level DFT calculation was used for the ROESY analysis. All structures up to a deviation of 2.5 kcal/mol from the lowest energy conformer were used to determine the ECD frequencies in a TD-SCF calculation on the same level as the former DFT optimization. The ECD curve was calculated from the Boltzmann-weighed contributions of all conformers with a cut-off level of two percent.
